# Causal discovery in biodiversity-mediated social-ecological systems

**DOI:** 10.1101/2024.05.26.595962

**Authors:** Maria J. Santos, Pengjuan Zu, Debra Zuppinger-Dingley, Maarten B. Eppinga, Anubhav Gupta, Frank Pennekamp, Cheng Li, Sarah Mayor, Camilla Stefanini, Yuji Tokumoto, Sofia J. van Moorsel, Marylaure Harpe, Martin O. Reader, Lidong Mo, Veruska Muccione, Meredith C. Schuman

**Affiliations:** Department of Geography, University of Zurich, Winterthurerstrasse 190, Zürich CH-8057, Switzerland; Department of Environmental Systems Science, ETH Zurich, Schmelzbergstrasse 9, CH-8092, Zürich, Switzerland; Department of Evolutionary Biology and Environmental Studies, University of Zurich, Winterthurerstrasse 190, Zürich CH-8057, Switzerland; Department of Biodiversity and Conservation Biology, Eidgenossische Forschungsanstalt fuer Wald, Schnee, und Landschaft (WSL), Zürcherstrasse 111, 8903 Birmensdorf, Switzerland; University of Miyazaki: Miyazaki, Japan; Department of Nature and Landscape, Canton of Graubünden Office for Nature and the Environment, Switzerland; Department of Chemistry, University of Zurich, Winterthurerstrasse 190, Zürich CH-8057, Switzerland

**Keywords:** causal inference, social-ecological interactions, big data, biodiversity and ecosystem function, ecosystem services

## Abstract

Global biodiversity loss and climate change exacerbate feedbacks within social-ecological systems, i.e., between ecosystems, their services and well-being of human societies. Our ability to mediate these feedbacks is hampered by incomplete understanding of the underlying causal links, which could benefit from interdisciplinary approaches to discover theoretical or empirical links from heterogeneous data characteristic of social-ecological studies. We propose a novel framework connecting literature-based causal knowledge with data-driven inference of causality. We test this framework for the highly biodiverse island of Borneo by conducting a systematic literature review of 7473 studies over 170 years, and a causal inference analysis for three conceptual causal diagrams connecting global change, socio-economics, ecosystem services, and biodiversity-ecosystem function using a set of 227 spatially explicit variables. We find that, while natural or social processes have been mostly studied independently, a set of studies already documents causal links across social-ecological domains for processes related to deforestation, food or energy. Causal discovery unveiled consistent negative causal links between global change, social-economic landscape, and biodiversity-ecosystem function, and positive causal links between global change and socio-economics, and these links were robust to indicator selection and addition. We detected few and weak links between social-economic landscape, global change, and ecosystem services. When comparing the data-driven *inferred* causal links to those *documented* by the literature, we find that links between biodiversity and ecosystem function with global change, and links between social-economic landscape and ecosystem services were also consistent, and causal analysis uncovered new (potential) causal links not yet described in the literature.

**Significance Statement:** Addressing climate change and biodiversity loss in the Anthropocene requires us to recognize that human societies and ecological systems are inherently interconnected in complex adaptive systems. Causal understanding in social-ecological systems enables understanding system dynamics and response to pressures and shocks. While promising, few studies have studied these systems using a combination of ‘big literature’ which provides the state-of-the-knowledge and ‘big data’ that provides the underlying information for causal discovery. With this framework, we can specify and rigorously test, causal links in biodiversity-mediated social-ecological processes under global change and examine potential interventions that lead to much needed sustainable outcomes.

## Introduction

Addressing climate change and biodiversity loss in the Anthropocene requires us to recognize that human societies and ecological systems are inherently interconnected in complex adaptive systems, i.e., social-ecological systems (SES) (1–3). Understanding SES feedbacks is crucial in predicting the functioning of our Earth System and society (4–10) and its responses to current global changes and challenges. Yet, research has mainly examined these processes by focusing on either ecological or social and institutional components, simplifying their interactions and long-term dynamics (11), and rarely inferring causality (12). Causal discovery, the identification of underlying causal relationships using observational data, provides an exciting avenue to overcome limitations and understand the feedbacks between global change, and the ecosystem processes and services that support human societies. Causal discovery has the potential to move forward our understanding of SES behavior from the current wealth of approaches that approximate causality (13). However, causal discovery in SES is complex and remains challenging (14) and hampered by the requirement to integrate across fields of knowledge (15–17), using very different types of information (14), analyses and methods (1, 18, 19), and vocabulary (20). Further, direct manipulation of SES is logistically impossible or unethical (6, 21, 22), and so establishing causal relationships in this setting requires new and creative ways of integrating knowledge, data and analyses.

The complexity of first- and higher-order interactions between variables that define processes in SES, and the hard-to-address assumptions of causality (8, 23), have limited our ability to infer causality. Yet, some pioneering examples have attempted to address this problem, for example through the investigation of causal links between acorn masting, deer, moths, and Lyme disease in humans (24, 25); earth system processes (4); ecosystem service cascades that link biodiversity and human well-being (26); and climate change (27, 28) and climate-driven economic downturn related to large-scale human crises in current (21) and historical periods (11). As such, causal analyses establish the parallel between a causal graph that defines the assumed relationships between drivers based on prior knowledge – for instance, expert knowledge and previous empirical assessment; and compares it to empirical or modeled results to identify consistencies, but also potential missing links or relationships between drivers, as well as rejecting assumed relationships (29). The underlying principle is that, as the cause happens *before* the effect, it carries unique information about the future values of its effect which can be extracted by the analysis of series of events. As many of these events may occur synchronously but in clusters, or asynchronously, throughout a given landscape, approaches such as space-for-time substitutions may also provide opportunities for conducting causal analyses using spatial data (30–32). Causal analyses are conceptually and philosophically well-established from experimental data in many research fields (33–35), and some methods for causal analyses are specific for some data types (30–32). Thus while individual methods may have limited support for data integration and method transferability (5, 6, 36, 37), the range of techniques (38) and the flexibility of many to deal with different data types and structures, make them a promising approach to derive new insights from interdisciplinary data (4, 39, 40) characteristic of SES.

The emergence of computer-assisted methods for research synthesis -- “big literature” and "big data" - herald the development of new complementary approaches needed for causal discovery in SES (4–10). Through the analysis of “big literature” (41), where literature synthesis results are weaved with their influence, as in research weaving, we can describe what is *documented* and discover connections between concepts, and knowledge transfer and influence across authors, institutions and disciplines. The outcomes of research weaving serve as a starting point for causal analysis: for causal inference, where robust identification of causality is conducted, but also for causal discovery, where the underlying causal structure is not known but is *inferred* purely based on the analysis of observational data. Integrating research weaving with causal discovery therefore combines what is *documented* with what is *inferred* (**Fig. 1**), taking advantage of novel tools that enable the examination of “big data”. “Big data” science (42) has accelerated the development of analytical methods which are at least partly agnostic of data type, and able to deal with large data sets (>10 000 observations and >100 variables) with the potential for directly linking data across disciplines. For example, big data approaches for the classification of articles based on their contents, i.e., systematic mapping, allied with meta-analytical methods (43, 44), can provide synthesis on knowledge and trends in vocabulary and disciplinary evolution (e.g. (45)). Causal discovery, recently has experienced major developments in algorithms and applications to observational data, which are becoming more readily available and have the potential to link data from social and ecological domains as required for causal discovery in SES (5, 21, 36, 46). These “big literature” and "big data" methods have recently been applied to several disciplines related to SES: ecology (38, 47–49), climate science (50, 51) uncertainty analyses (52), and sustainability (53), where they enable the combination of data types and spatial and temporal processes through causal links, and derive predictions of system behavior. This is useful for SES understanding as it might help to identify how, for example, biodiversity-dependent domains such as ecosystem services and global change have been studied, what we know about their interactions and the strength of evidence that support them (54), and where the gaps in our knowledge lie. Further, it might help to distinguish whether changes in biodiversity cause changes in the socio-economic landscape (55) or in ecosystem services (56), or unveil interactions across the domains of biodiversity-dependent SES.

**Figure 1.**
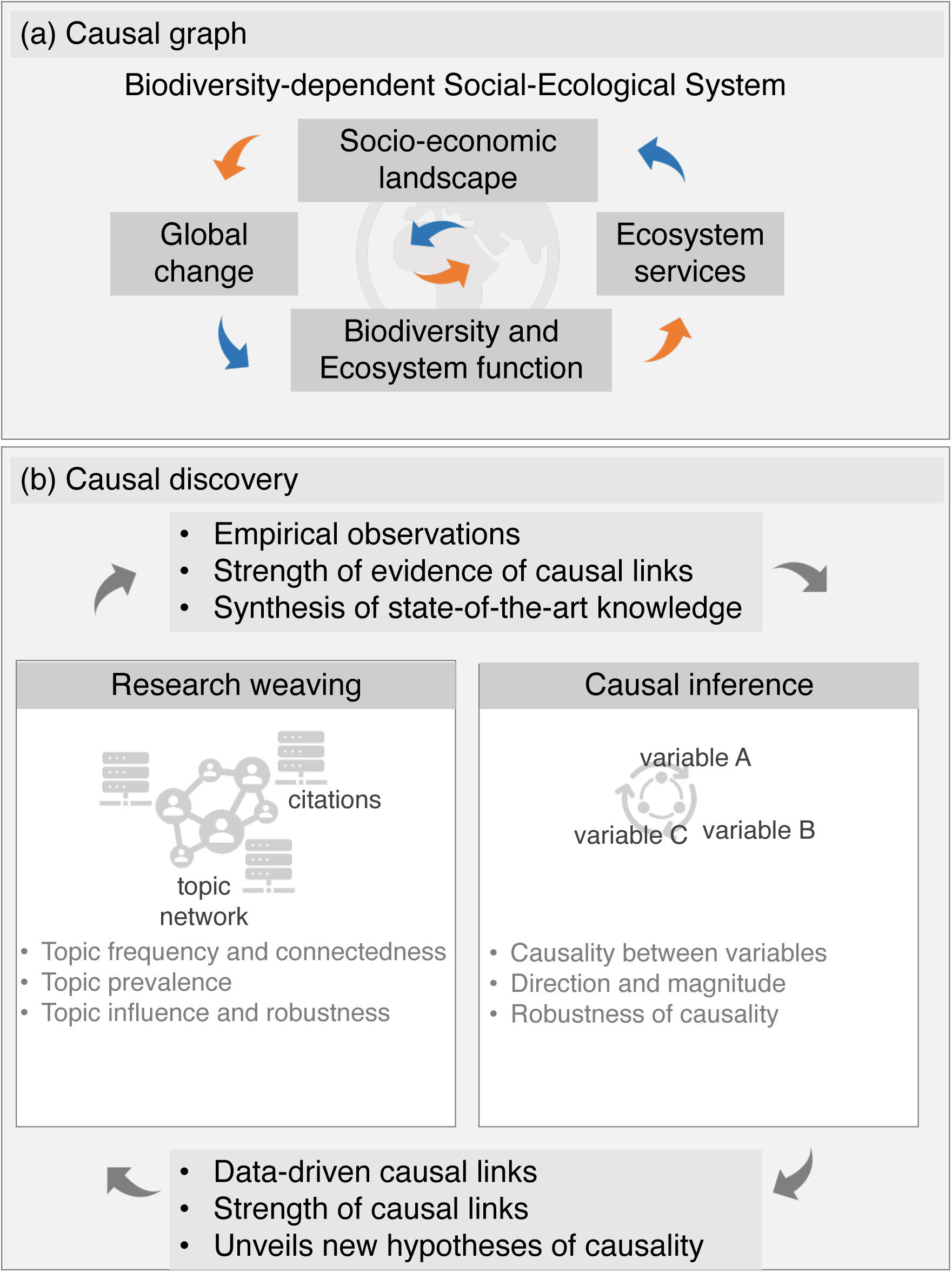
Causal discovery: combining research weaving (“big literature”) and causal inference (“big data”) to unveil causal link in social-ecological systems.

Here, we make use of these emergent techniques and combine research weaving with causal discovery to gain new insights about causal links in SES. Our proposed approach comprises a data-driven alternative and automated way to identify trends, advance hypotheses and highlight knowledge gaps in our understanding of SES feedbacks. We test our approach on the example of Borneo, an island hotspot of biodiversity with a long history of human presence. The Borneo island system has previously been used as an illustration of how interconnected SES enable system resilience (57). In the 21^st^ century, expansion of logging concessions in Borneo led to levels of deforestation (58) and forest degradation that precluded El Niño-driven processes of forest regeneration: El Niño current effects are destructive, with interruption of species reproductive cycles and fire patterns, and such changes are predicted to continue (59). We first examine the published literature about Borneo to reveal the evolution of social and ecological research in this system, and identify research gaps through research weaving (60, 61). With this approach, we identify the strength of evidence around *documented* links among the social and ecological dimensions of SES. We then conduct causal inference over a novel spatially explicit dataset with 227 social and ecological variables for Borneo to examine the probability of links across four broad domains: ‘Biodiversity-Ecosystem Function’, ‘Ecosystem Services’, ‘Socio- Economic Landscape’ and ‘Global Change’, and how much these links depend on the amount or identity of input data. We use LiNGAM, i.e., the Linear, Non-Gaussian, Acyclic causal Model framework, for causal discovery. LiNGAM derives structural equation models or linear Bayesian networks depicting the most likely causal relationships (*48*, *49*). Structural causal model methods such as LiNGAM that do not require prior graphs have the potential to be applied to cross-sectional data, as long as spatial autocorrelation is properly dealt with (30). We furthermore use a set of simulated data to test the performance of LiNGAM and to examine effects of parameter behavior, distribution shape, sample size, scaling and nesting of variables to understand the algorithm’s applicability to SES research. Finally, we compare the data-driven *inferred* causal links to those *documented* (i.e., those reported in the literature) and draw conclusions on the applicability of this framework to the study of social-ecological systems.

## Results

For the nearly 7500 scientific publications we examined across the four domains of an SES, about 70% investigated a single domain (n=5234 out of 7474 publications; **Fig. 2**; flow diagram of the systematic review in **Fig. S1**) and the number of domains investigated increased with publication year (1860-2020, **Fig. 2b**). The most frequently identified domain was ’Biodiversity-Ecosystem Function’ (54%, n=4024, significantly higher than the other domains: Χ_BiodEcosFunct- EcosServ_ = 4645, p < 0.0001, Χ_BiodEcosyFunct-SocioEcon_ = 3378, p < 0.0001, Χ_BiodEcosFunct-GlobChange_ = 4711, p < 0.0001), followed by ’Social-Economic Landscape’ (10%, n=717, Χ_SocioEcon-EcosServ_ = 232, p < 0.0001, Χ_SocioEcon-GlobChange_ = 259, p < 0.0001), while ’Ecosystem Services’ (n=257) and ’Global Change’ (n=236) occurred less frequently (and did not differ significantly in their frequencies Χ_EcosServ-GlobChange_ = 0.93, p = 0.34; **Figure 2a-c**). Among literature that covered multiple domains, publications combining two domains were overrepresented (randomization tests, all p < 0.0001; Supporting Information A), with the exception of entries combining ‘Social-Economic Landscape’ and ‘Global Change’ (randomization test, p = 0.56). Only 4% of publications examined three-way domain relationships (n=462), for which all combinations including ’Biodiversity-Ecosystem Function’ were underrepresented (randomization test: p < 0.0001, Supporting Information A). This is somewhat surprising, as almost half of the publications examining two domains covered both ’Biodiversity-Ecosystem Function’ and ‘Global Change’ (n=691; **Fig. 2a-c**). The remaining three-domain combination between ‘Social-Economic Landscape’, ‘Ecosystem Services’ and ‘Global Change’ was neither over- nor under- represented (randomization test: p = 0.24; Supporting Information A). One percent of the studies (n=87), examined the four domains in some combination (**Fig. 2c**; randomization test: p < 0.0001; Supporting Information A). This may reflect a growing trend of interdisciplinary research across domains, research networks and novel institutional architectures that facilitate integration. We found that the most frequent word stems were ‘speci’, ‘forest’, ‘borneo’, ‘use’, and ‘tree’, corroborating the dominance of ‘Biodiversity-Ecosystem Function’ literature (**Fig. 2g**). Finally, we conducted a citation and bibliometric analysis on the subset of papers that already addressed the four SES domains and find that the most frequently co-occurring words clustered into seven groups, with the main topics being conservation (n=128), biodiversity (n=36), deforestation (n=24), land use (n=11), and management (n=10), the latter being closer to ecosystem services and social-economic terminology (**Fig. 2h**). For this subset of papers, we found that a few authors commonly co-cited each other (**Fig. 2i**), and the average citations per article per year tended to increase over time, peaking at 8-12 (**Fig. 2j**).

**Figure 2.**
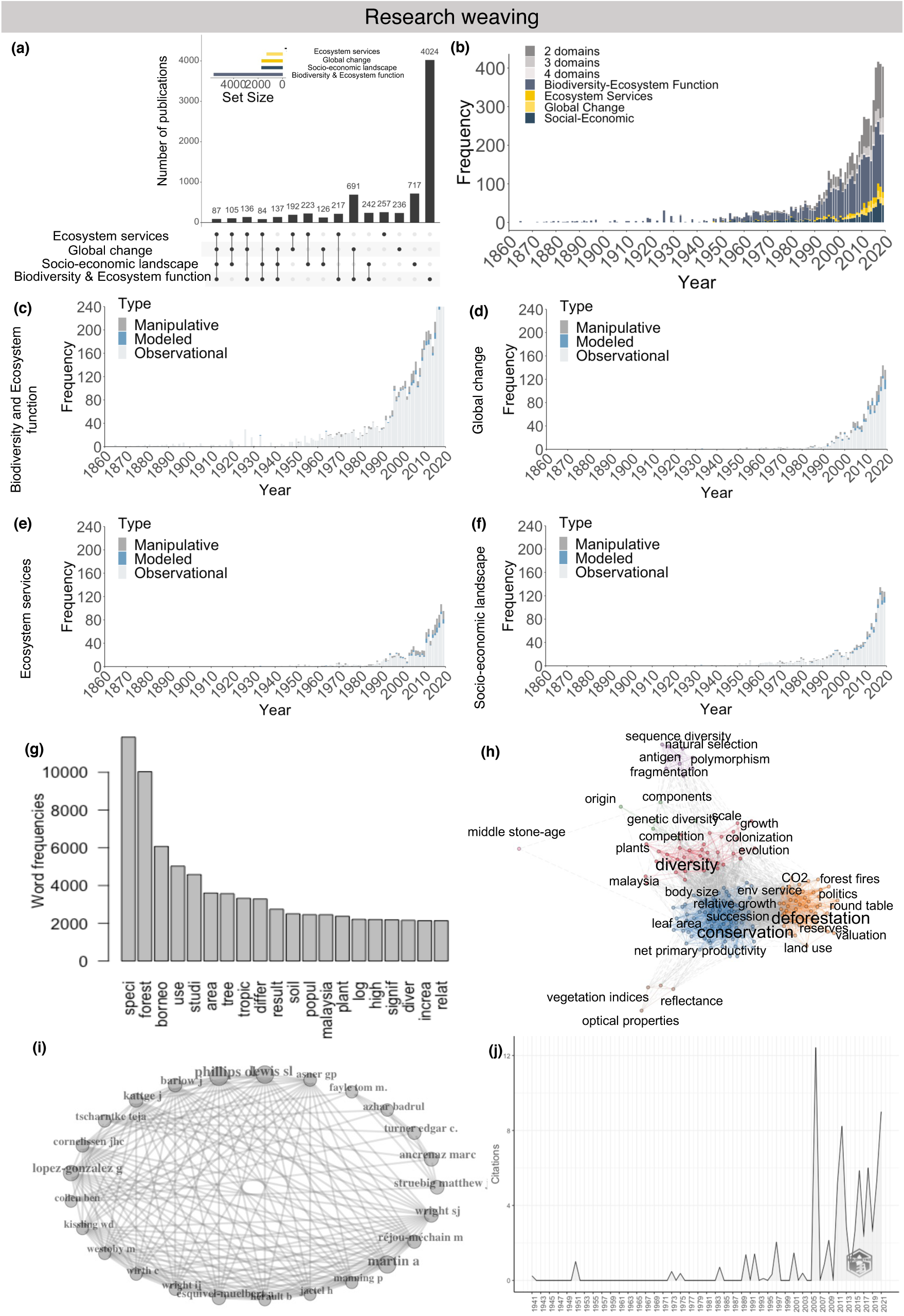
Research weaving results for the island of Borneo: (a) Number of publications per SES domain, (b) temporal trends in topic prevalence among the domains of SES, (c) temporal trends in ‘Biodiversity-Ecosystem Function’, (d) temporal trends in ‘Global Change’, (e) temporal trends in ‘Ecosystem Services’, (f) temporal trends in ‘Social-Economic Landscape’, (g) most common words in abstract text, (h) co-occurrence network of the top 200 words (larger font words are most common words per cluster), (i) author citation network for the subset addressing all four domains, and (j) average number of citations per article per year for the subset addressing all four domains.

We tested an approach to uncover causal links between the four domains of SES using LiNGAM, a well-researched method for causal discovery (64, 65). We began with three example causal graphs comprising four core variables (one variable from each domain). These were: ‘tree density’, ‘cropland’, ‘human development’ and ‘human footprint’ (causal graph 1), ‘species richness’, ‘food area’, ‘investment’ and ‘phosphorus loading’ (causal graph 2), and ‘species richness’, ‘soil water capacity’, ‘human population density’ and ‘forest loss’ (causal graph 3), chosen for their documented causal relationships in literature (**Fig. 3**; Supporting Information B). LiNGAM is suitable for discovering linear acyclic causal relationships from data with non-Gaussian error terms (64, 65) and so generally suitable for SES data (although in some cases non-linear methods are advantageous), and can be expanded to spatial data when addressing spatial autocorrelation (30) (see Supporting Information C for spatial autocorrelation). We found generally negative causal relationships of ‘Global Change’ and ‘Ecosystem Services’ domains on ’Biodiversity-Ecosystem Function’ (**Fig. 3**, blue lines with arrows). For example, we found that human development, human footprint, and cropland all had negative causal effects on tree density (**Fig. 3a**); food area and phosphorus loading reduced species richness (**Fig. 3b**); and forest loss decreased alpha diversity (**Fig. 3c**). These results from LiNGAM are in line with our expectations (Supporting Information B), except for the link between phosphorus loading and species richness, which is positive according to the literature (66, 67). In contrast, we found no positive causal relationships involving ‘Biodiversity-Ecosystem Function’ variables (**Fig. 3**), which is surprising as positive relationships have been reported for example with ecosystem services (68, 69). We then gradually added more variables from each domain to examine whether the revealed causal relationships were context-dependent. First, we added variables equally from each domain, from one to eight variables per domain (corresponding to the domain with the fewest variables) from a set of 227 variables spanning the four domains, each represented by 3289 co-geolocated gridded data points across Borneo (**Table S3** for variable details). We found that all causal relationships had intermediate robustness, i.e., could be eliminated when variables were added, and sometimes even transitioned to opposite causal direction; and this pattern was consistent across the three examples (**Fig. 3**, mix-colored bars between the causal variables).

**Figure 3.**
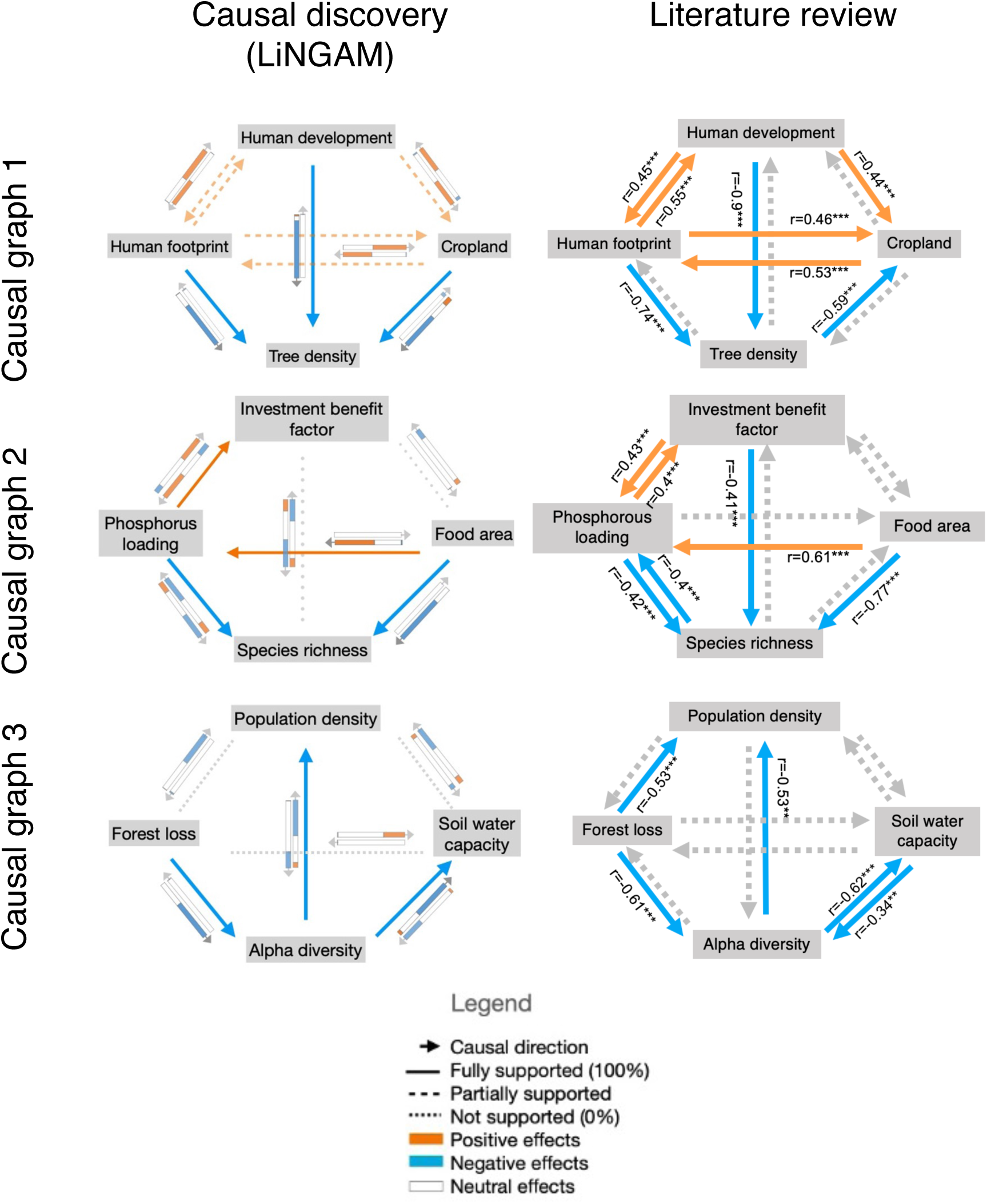
Causal relationships for the three example causal graphs. Top row: Results from LiNGAM: Lines with arrows show sign and direction of relationships when analyzing only core variables. Bars with arrows show the distribution of sign and direction of additional variables; the length of different colors in the bars indicate the proportions of cases that support the corresponding causal effects (see **Table SC3** and **Fig. SC3** for more details of the results when adding variables). Bottom row: Results from literature review. We display the number and count of words related to the core variables from the literary corpus examined (title and abstract) for the three example causal graphs. Arrows show direction and sign of the relationships: blue for negative and orange for positive. Colors match those of Fig. 2. Examples of word counts and associations are given in **Table 1**.

**Table 1.**
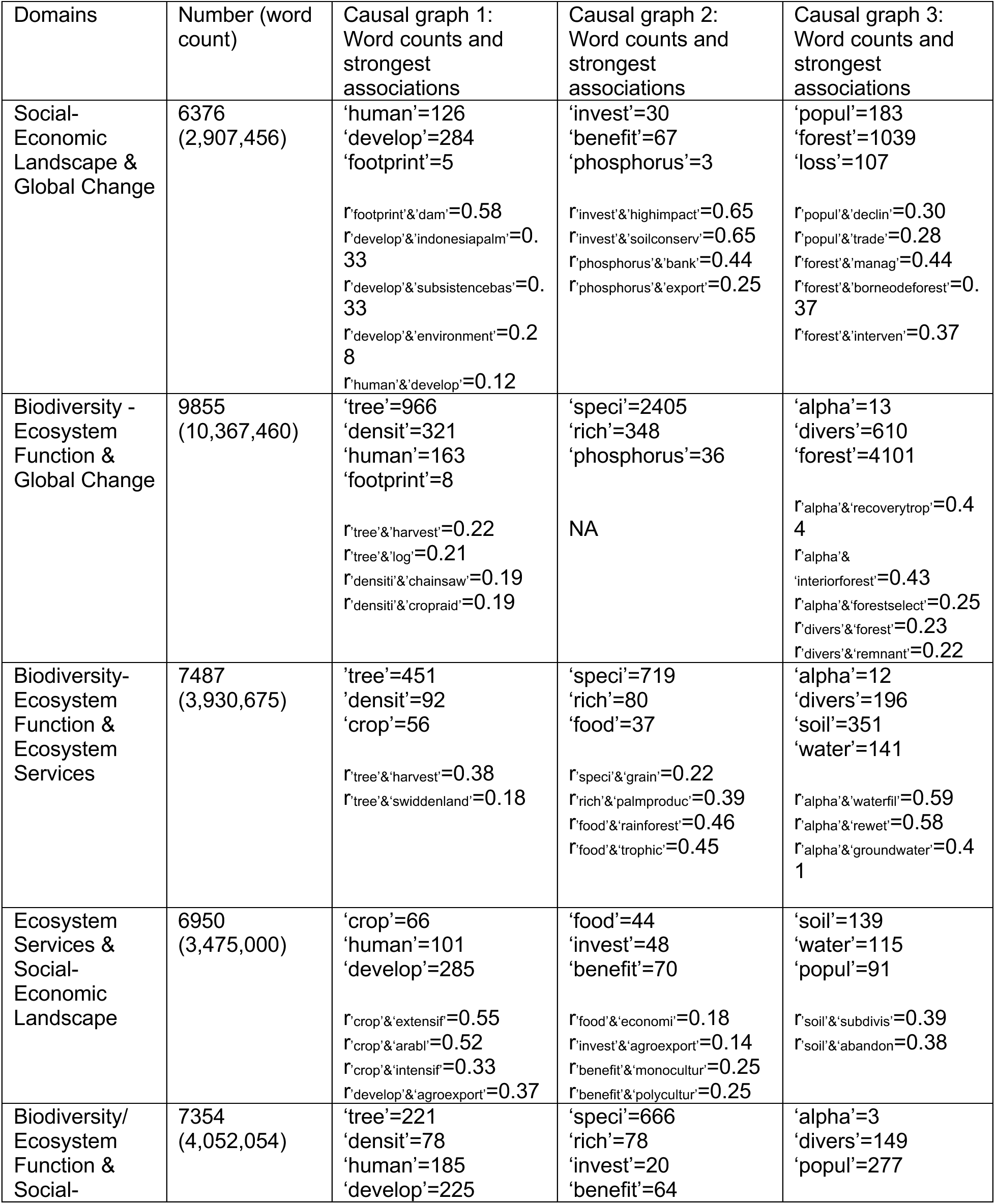
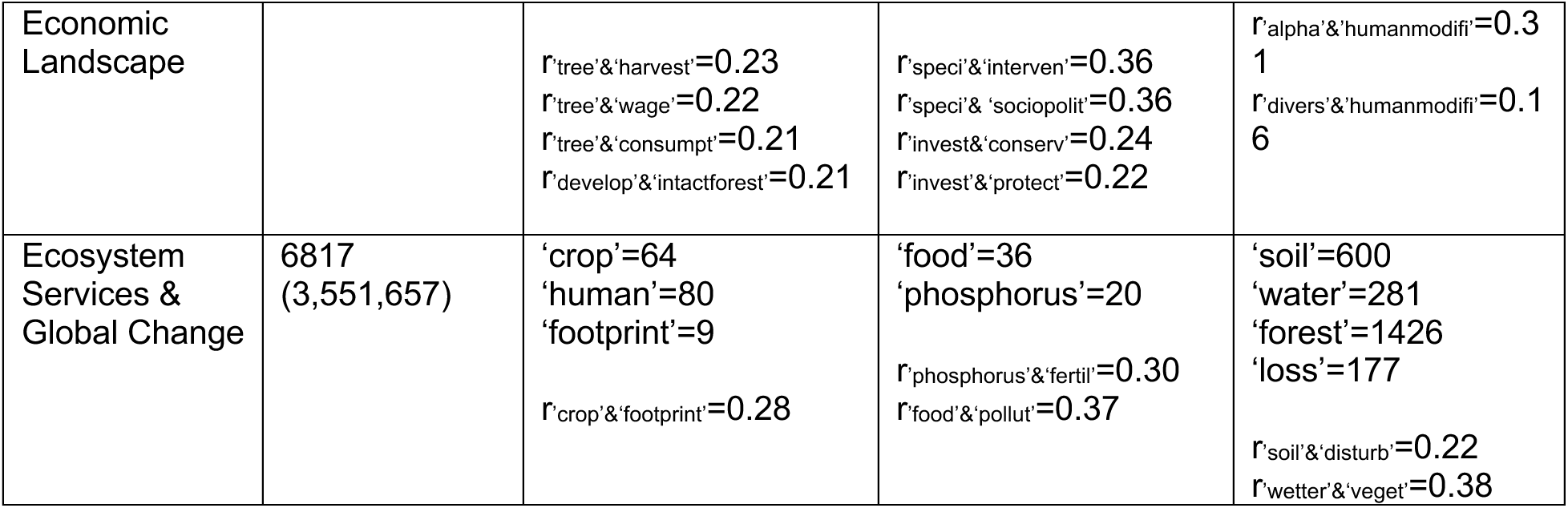
Causal discovery in SES: Links between research weaving and causal inference. We display the number and count of words related to the core variables from the literary corpus examined (title and abstract), and for each causal graph we show the top word associations ranked by the strength of the correlation coefficient. For analysis of the full corpus, see **Table SE1**.

This could be a result of LiNGAM not accounting for latent variables (4, 29, 62) and the fact that the inferred links are contingent on some other input variables. For example, cropland had a negative causal link with tree density when examining only the core causal graph (blue arrow, **Fig. 3a**), but as more variables were added, in ∼60% of cases this causal link remained negative, in ∼30% it changed to neutral, and in ∼10% it changed to positive (**Figs. 3a**, **S6**, **Table S3**).

Results were similar when we selected variables randomly in proportion to the representation of domains in the 227-variable dataset (i.e., 20 from ‘Biodiversity-Ecosystem Function’, 1 from ‘Ecosystem Services’, 1 from ‘Social-Economic Landscape’, 6 from ‘Global Change’) (**Fig. S6**).

We also used simulations to examine how well LiNGAM performs to retrieve unknown causal links depending on correlation strength (see methods for details and Supporting Information C and D for details on the results). LiNGAM performed better at identifying causal links when the absolute value of the LiNGAM coefficient B was between 0.3 and 4, versus above 5-10 (very strong), when predictive ability started to show multimodal distributions, and below which the predictive ability dropped. This corresponded to correlation coefficients between ca. 0.25 and below 1, representing strong associations between variables in ecological and social studies. The inability to predict high values may be because the algorithm underlying LiNGAM relies on asymmetry in the noise of correlations, which is likely to be low for very strong correlations (29, 62). LiNGAM performed equally well for imperfect gaussian, bimodal and skewed distributions.

Variable scaling deteriorated the lower threshold of B but showed very similar patterns. The lower threshold of B decreased by adding more samples, lowering the mean error and increasing the number of links in the causal graph.

Finally, we compared the data-driven *inferred* causal links to those *documented* in the literature to find that the identified causal links were mostly consistent with word associations from the literature review (**Fig. 3**, **Table 1**). We found strong associations between words about ‘Biodiversity-Ecosystem Function’ and ‘Global Change’ (e.g., ‘alpha’ and ‘recovery’, ‘diversity’ and ‘remnant’) and ‘Social-Economic Landscape’ (e.g., ‘tree’ and ‘harvest’, ‘species’ and ‘interventions’) and ‘Ecosystem Services’ (e.g., ‘tree’ and ‘harvest’, ‘rich’ and ‘palm production’, ‘alpha’ and ‘water’), but some strong word associations did not correspond to causal links identified by LiNGAM, and vice versa. For example, causal inference failed to identify well-established relationships between ‘Social-Economic Landscape’ factors and ‘Ecosystem Services’ (e.g. ‘Food area’-‘Investment benefit factor’, ‘Soil water capacity’-‘Population density’) and between ‘Global Change’ and ‘Ecosystem Services’ (e.g. ‘Soil water capacity’-‘Forest loss’). Our dataset contained fewer and less well-developed variables in these domains than for ‘Biodiversity-Ecosystem Function’. On the other hand, we found little evidence in the literature for the relationships of ‘Biodiversity-Ecosystem Function’ and ‘Ecosystem Services’ to ‘Global Change’, which were highlighted strongly in our causal inference results (e.g. ‘Species richness’- ‘Phosphorus loading’, ‘Tree density’-‘Human footprint’, ‘Tree density’-‘Human development’, ‘Cropland’-‘Human footprint’), potentially indicating causal links yet to be researched and new hypotheses to be tested.

## Discussion

Here, we show that a novel combination of research weaving with causal discovery allows us to test whether *documented* causal links from the literature are *inferred* by data-driven causal discovery in a large observational dataset. We find that existing literature on the system we chose, the hyperdiverse island of Borneo, rarely examined two or more SES domains, and is enriched with single domain studies on Biodiversity and Ecosystem Functioning and Global Change. These results may point to the lower data availability for ecosystem services and social-economic landscape variables, particularly in peer-reviewed international literature. Yet, we still found an emerging set of studies across the four domains, which generally cover aspects related to ecosystem services, political ecology and land use in the island (70–72), all natural resource management aspects that are better analyzed when taking a SES approach. For instance, Obidzinski et al. (72) showed that oil palm production would lead to social inequalities, as the benefits of the production were not evenly distributed and accompanied with restrictions on traditional land use rights and land losses, along with well-documented deforestation and secondary external impacts such as water pollution, soil erosion, and air pollution.

We furthermore discovered many interesting links using causal inference in a process that was fairly robust for the three example causal graphs we examined, i.e., causal links changed with the addition of new variables, but not much. The causal links we discovered were often consistent with those described in the literature (Supporting Information), and aligned well with the text analysis of the ∼7500 papers we reviewed (**Fig. 3**). For example, for causal graph 1, we generally found a negative effect of all domains on ‘tree density’, which is aligned with the many deforestation processes ongoing in Borneo (70–72). Similar relationships were retrieved with the text analysis, yet in literature-based associations we found a predominantly negative effect of tree density on cropland, whereas causal discovery identified a negative effect of cropland on tree density. Text analysis also suggested many positive relationships not captured in our causal inference study. We interpret these as new hypotheses to be examined that might contribute to fill gaps in theory and data collection on SES (14, 17, 73). This would ensure that ‘big data’ becomes closer to ‘right data’, and that ‘big literature’ encompasses more ‘needed knowledge’ or ‘generalized knowledge claims’ (14). Similarly, for the other two example causal graphs (#2 and 3), we retrieved the (mostly) negative causal links also identified in the literature. For example, we found negative effects of phosphorous loading and food area on species richness, as also indicated by the word association analysis. In this case, we know that phosphorous loading, especially after deforestation, greatly changes the composition and quality of the soil for subsequent forest or other land uses (66, 67, 74), corroborating this negative effect. Finally, and only for causal graph 2 (linking investment factor with food area, phosphorous loading, and species richness), we retrieved the positive effects reported in the literature between phosphorous loading and investment benefit factor, likely associated with the investments on fertilizer to produce better crop outputs.

While confident of these results, it is important to mention two aspects that are fundamental to our causal discovery analysis. First, we used an ensemble of opportunistic, static, spatially-explicit variables to infer causality; while useful as a first approach, longitudinal studies and time series data are likely to better support testing of causal relationships (75). Spatial processes captured through spatial autocorrelation analyses may still be present in the data and dilute our ability to identify the strength and directionality of the relationships (31, 32). As such, it would be interesting to compare our results with other datasets from census or other collection methodologies (qualitative and quantitative) to see whether relationships follow the same or different directions from those we identified, and also to try different algorithms such as a spatially explicit causal inference algorithm recently introduced by Gao and colleagues (32). Second, we use a method for causal discovery that is based on the correlation structure between variables to determine the probability of causal links. Correlation and causation are related to the extent that causality can be called a special (and especially complex) case of correlation. Our simulation analysis indicates that LiNGAM performs best when correlation values are between 0.25 and 1 and thus fails to identify associations of variables with weak linear correlations, which was also recently demonstrated on selected Earth observation datasets (32). In some important cases, causal relationships do not have a clear signature of linear correlation. This is elegantly demonstrated by Gao and colleagues (32) in their analysis of the relationship between copper pollution, industrial activity and night light, where neither Pearson correlation nor LiNGAM could detect causality that was detected using spatially explicit Geographical Convergent Cross Mapping. The assumptions of linear relationships and acyclic behavior between variables built in to LiNGAM are not always met in the real world, yet they provide interpretative simplicity and perhaps help to filter for relationships that can be fruitfully targeted to manage SES processes.

Data representing single timepoints or shorter time scales, which common in SES as in other fields (32), may often not carry interpretable signatures of feedback loops that alter system dynamics. Thus, the acyclic character of LiNGAM and many other causal discovery methods may help not to comprehensively discover causal relationships, but rather to preferentially discover relationships that are amenable to measurement and management.

While there may yet be a long road to design studies that address SES cross-domain interactions along with transparent methods that allow for integrative synthesis and comparative studies (14, 17, 73), our results suggest that combining research weaving with causal discovery may already provide an avenue to advance our understanding of interactions in SES. The proof-of-concept analyses we present here, comparing LiNGAM-extracted causal relationships with those documented in the literature, indicate robustness and highlight potential “low-hanging fruit” which could be harvested to fill existing knowledge gaps. As more sophisticated methods are emerging (65, 76, 77), more nuanced analyses are becoming possible for complex systems such as SES. Yet, our approach synthesized knowledge on Bornean SES (7, 8, 23) and could be applied to any system where literature and data on co-occurrence of ecosystem services (78, 79), land systems and archetypes (7, 80), and other SES components are available. Earth observation generates a growing number of such datasets, often in time series, which is an added advantage for causal inference, and these data have enabled the development of an Earth systems approach and are increasingly leveraged in SES research. Developing our capacity to infer and test causality from such observational data is fundamental to the attribution of change due to ongoing global change-driven biodiversity loss (81, 82), and to studying SES behavior in response to these pressures (83).

### Materials and Methods Research weaving

To explore published studies for the island of Borneo, we conducted a systematic search for English language articles from the oldest available date (1851) to September 2020. Specifically, we used a list of search terms related to ‘Biodiversity-Ecosystem Function’, ‘Ecosystem Services’, ‘Socio-Economic Landscape’, and ‘Global Change’ (**Fig. S1**) to retrieve the literature in Scopus (Elsevier) and ISI Web of Knowledge (v.5.34, Clarivate). To combine articles from the searches in Scopus and ISI, we retained and matched columns containing the DOI, year of publication, document type, title and author(s) of the publications. The resulting 50,198 publications were evaluated according to the relevance of each paper to the scope of the study using both automated (using a custom R script) and manual evaluation procedures as described below.

### Detailed list of search terms

We searched for combinations of terms that included: (Borneo OR Bornean) AND biodivers*; (Borneo OR Bornean) AND taxonom*; (Borneo OR Bornean) AND phylogen*; (Borneo OR Bornean) AND species; (Borneo OR Bornean) AND trait; (Borneo OR Bornean) AND function; (Borneo OR Bornean) AND soci*; (Borneo OR Bornean) AND econom*; (Borneo OR Bornean) AND (“climate change” OR “global warming” OR “global change”); and (Bornean OR Borneo) AND ("deforestation" OR "land use"). We then deleted all duplicates that shared either the same DOI and/or title, resulting in the exclusion of 8,202 duplicates. The outputs based on keywords "(Bornean OR Borneo) AND phylogeny*" and "(Bornean OR Borneo) AND taxonom*" mostly corresponded to descriptions of taxonomic entities, their revisions and were not related to our domains of interest so we excluded them from further analysis (n=20,464 articles).

We then developed a customized R script to systematically remove articles which were not directly related to social-ecological systems. This enabled us to remove an additional 7129 articles, which included the following terms in their title: "nouveau", "check-list", "review of the genus", "identity", "checklist", "neue", "studies on the", "new taxa", "new genera", "new records", "known species", "description", "new species", "new subspecies", "new genus", "new", "revision of the", "carcas", "fossil", "taxoplasma", "nouvelles", "taxonomy", "a record of", "records", "revision of", "a contribution to", "the genus", "notes on", "a catalog of", "revision", "lycaenopsis", "butterflies of", "genera of", "list", "remarks" and "notes". Finally, we merged the results from the two databases, which resulted in the further exclusion of 2617 duplicates. We then manually revisited the title and abstract of the remaining articles (n=11,786) and further excluded 4313 articles, and a set of 7473 articles was then used for the subsequent analyses.

The next was to manually code the articles for analyses, based on a set of criteria. We decided on the following attributes to be coded: Responsible (person coding), Keep (relevant for study), Domain (four domains and their combinations), Inappropriate (due to exclusion criteria), Scope and Type (**Table S1**). Before we proceeded with the manual coding of the articles that remained after the automatic filtering, we conducted a sensitivity test to establish whether the eleven different persons involved in the coding would assign the same domain to the same paper (see Coding Consistency Analysis). To this aim, we decided on a subset of 114 publications and individually assigned domains (as outlined below) and categories (**Table S1**) to each paper.

Overall, there was 65-80% agreement among coders on the domain identity assigned to any given publication, and on how many domains were assigned (see ***Coding consistency analysis*** below). Based on these results, we used the quantitative word frequency and co-occurrence analysis to compare with the results of coding, and coding results were also checked for consistency by a reduced team of three authors. We also asked coders to assign more specific categories to studies, but because a preliminary consistency analysis showed that the variability in these assignments was much higher than for assignment of domains, we decided to not attribute specific categories to the publications; however, the reduced team did use the categories assigned by individual coders to clarify which domains to attribute, especially in the case of multiple domains. In the following, we describe the attributes that we opted to include. All studies were classified as belonging to one of the following domains (**Table S1**):

- **Socio-Economic Landscape:** Results about human social and economic systems
- **Global Change:** Results about global change factors, including land- and sea-use change, direct exploitation of organisms, climate change, pollution, and invasion of alien species
- **Biodiversity-Ecosystem Function:** Results about biodiversity, including distribution, abundance, richness, and evenness of genotypes, species, other phylogenetic or taxonomic groups, or functional groups
- **Ecosystem Services:** Results about ecosystem services, including provisioning, regulating, supporting, and cultural services

Where more than one domain applied, we included all relevant domains, favoring inclusivity. For example, we coded results about conservation as Social-Economic Landscape and Biodiversity-Ecosystem Function, or results on palm oil plantations as Global Change, Biodiversity-Ecosystem Function and Ecosystem Services because they describe the connections between biodiversity, land-use, and provisioning as an ecosystem service. In addition to the domain, we coded scope and type of entry. Scope included “within” or “beyond” to designate studies which had been conducted in Borneo, of which the findings related only to Borneo (“within”) or studies that included information from Borneo/related to Borneo as well as other areas (“beyond”). The type of study was classified as “observational”, “manipulative” or “modeled”. With study type “observational”, we coded studies for which data were collected through observations, i.e., direct measures of the state of a property without manipulation. With study type “manipulative”, we coded studies for which data were collected through measurements of a property after a manipulation was introduced in the system (e.g., experiments at any level of the system). With study type "modelled" we coded studies for which data were produced as an output of a model, and these did not consider the results from statistical significance tests. If none of these types could be assigned, we classified the type as NA. We also excluded some studies and coded them as "inappropriate" when they were not of relevance to any of the four domains or interactions between them, or in general not within the scope of this paper. Specifically, we excluded all entries that were classified to belong to any of the categories listed below (**Table S1**, list of domains and categories for manual selection). All of these assignments were also curated by the reduced team of three coders.

This yielded 7473 publications (**Fig. S1**). After the coding process, we conducted one final manual clean-up of the data set with the same reduced team of three authors to remove spelling errors and coding mistakes as well as to standardize the order of domains when multiple domains were coded, but no further publications were excluded during this step.

#### Coding consistency analysis

To check for consistency in coding of the different SES domains by the eleven evaluators that initially conducted this task, we offered a set of 114 practice entries and recorded for each practice entry which and how many domains were assigned (1, 2, 3, 4 domains, unscored, or not analyzable) by how many evaluators. Of the practice entries, 89 were assigned to domains by most of the evaluators (mode of 9), but only 10 were assigned to domains by all 11 evaluators.

On average, evaluators who assigned domains agreed 65% (mean for the ten entries coded by all 11) to 80% (mean for the 89 entries) on their assignment. When considering all 114 practice entries, which also included 25 that could not be assigned to a domain by one or more evaluators, it was most often the case (37 entries) that six out of 11 evaluators agreed on the most frequently chosen domain, and the majority (six or more) of evaluators agreed on the most frequent domain assignment for 85 of the 114 entries (**Fig. S3a**). We repeated the analysis for the number of domains and found that most often (38 of 114 entries), six out of 11 evaluators agreed on the number of domains assigned, and overall, the majority of evaluators (six or more) agreed on the number of domains assigned to 96 out of 114 entries (**Fig. S3b**). For the full dataset, individual evaluators’ coding was checked for consistency by a reduced team of three authors (see above). The coding results were then compared to the quantitative analysis of word frequency and co-occurrence from the literature as an independent test of representativeness (see below). See also the source data and Python script for the coding consistency analysis with the data and code for this article.

#### Trends in SES domains reported in the literature for the island of Borneo

To evaluate numbers of single- and cross-domain studies, we calculated the numbers and frequencies for the four domains and their combinations using the R package ‘UpSetR_1.4.0’ (84). We then plotted the prevalence of domains over time for the total number of evaluated publications and also splitting them by Within and Beyond Borneo, as well as by SES domain. We used randomization tests to determine whether the observed frequencies were statistically non-random (Supporting Information).

We also retrieved words from the abstracts of the 7473 publications and conducted a word stem frequency and co-occurrence analysis using the R packages ‘tm’ (85), ‘SnowballC’ (86), and ‘wordcloud’ (87). We first cleaned-up the corpus text document to (i) replace punctuation characters with spaces (e.g., replacing “/”, “@” and “|” with space); (ii) remove unnecessary white space; (iii) convert to lower case; (iv) remove stop words using ‘English stopwords’ as defined by a glossary embedded within the R package; (v) remove numbers and punctuation; and (vi) text stemming to reduce words to their root form (e.g., ‘develop’, ‘densit’, etc.). We conducted the analysis for all the publications in the corpus and by splitting them by those mentioning combinations of SES domains according to the coding results (see above).

We analyzed the corpus data to calculate the frequency of studies which addressed each of the domains and multiple domains, and the frequency over time of the type of studies for each domain, i.e. manipulative, modeled or observational studies. We then analyzed the frequency of words as single entities and their co-occurrence by calculating the correlation coefficient between words present in the same sentence and in close proximity. Finally, and taking the subset of papers that were coded as addressing all four SES domains, we conducted bibliometric analysis using the package ‘bibliometrix’ (88). We restricted the analyses to the papers that had full bibliometric data (77 out of 87), and calculated the co-occurrence of terms within this set of articles to identify the top 200 keyword co-occurrence pairs. Finally, we uncovered the author citation network for this subset of articles, to identify connections among these as well as the number of citations per article per year to assess the influence of these papers in the field.

### Causal discovery

To unveil potential causal relationships among the four SES domains, we used global-gridded (i.e., raster) data accompanied by sufficient meta-data detailing the spatial location from which a square region capturing Borneo was extracted (**Fig. S4**). This provided a snapshot at one point in time for particular variables. While the exact time of data collection differed between variables, they were all collected between 2010-2020. The data were gathered by the co-authors following due diligence to retrieve all available relevant spatial data using a combination of searches and expert knowledge (**Table S3**). We contextualize our dataset as retrieved by a geographically and gender diverse group of authors across many phases of the academic career in the natural sciences and with some affinity for social sciences. However, our approach does not assume a complete dataset and we acknowledge that even all available data would leave many gaps in the spatial and temporal characterization of each of the domains we investigate.

In contrast to most variables, longitudinal climate data is publicly available, allowing the calculation of temporal trends in bioclimatic variables over time. Specifically, we used monthly climatic data from the CHELSAcruts timeseries (89, 90) to calculate the widely used BIOCLIM variables (91) for two time periods: 1957 - 1986, and 1987-2016. Subsequently, climatic change variables could be calculated as the absolute difference between the two time periods.

The spatial resolution of obtained variables ranged between 30 m by 30 m and 1 degree by 1 degree (111 km by 111 km at the equator). Therefore, all variables were upscaled (in most cases) or downscaled to a standard resolution of 2.5 by 2.5 arc minutes. Downscaling was done for a set of ecosystem service variables, which tended to have the coarsest spatial resolution. For continuous variables, rescaling was done through 2-dimensional linear interpolation, while for categorical variables, the interpolated values of the standardized grid were based on the value at the nearest location in the actual dataset. Interpolation was done using the ‘interp2’ function as implemented in Matlab (92).

In the standardized grid described above, the Bornean land mass consisted of 17,136 pixels. However, many of the variables contained missing values for a proportion of these pixels. This posed a problem, in that the algorithms used for causal discovery (see below) were not able to handle missing data values. Hence, we selected a subset of 3289 pixels within the standardized grid that did not contain missing values for a selected set of 227 variables that covered the entire island of Borneo (**Fig. S4**), although data availability was lower for the center part of the island. The 227 variables were then categorized into the four domains described in the main text, with categories agreed to by five co-authors by running a pilot coding exercise and when there were coding disagreements, the co-authors discussed these disagreements and agreed on a final code. Overall, the dataset consisted of 162 variables for ‘Biodiversity-Ecosystem Function’, 11 for ‘Ecosystem Services’, 46 for ‘Global Change’, and 8 for ‘Social-Economic Landscape’ (see **Table S3** for details).

To discover causal relationships among domains variables and to further examine the robustness of these causal relationships with increasing knowledge (or new data accumulation), we first chose three example causal graphs, each consisting of four core variables (one variable from each domain). These were: ‘tree density’, ‘cropland’, ‘human development’ and ‘human footprint’ (causal graph 1), ‘species richness’, ‘food area’, ‘investment’ and ‘phosphorus loading’ (causal graph 2), and ‘species richness’, ‘soil water capacity’, ‘human population density’ and ‘forest loss’ (causal graph 3; **Table S2**).

We then evaluated causal relationships among these four core variables with and without additional variables from the pool of 227 variables. Due to the unequal number of variables in the four domains, we explored the effects of additional variables to the core causal graph in two ways: (1) balanced addition experiments where equal numbers of variables from each domain were added to the core causal graph, i.e., one from each domain, two from each domain, up to 7 from each domain (7 was the maximum number of additional variables we could retrieve for the ‘Social-Economic Landscape’ domain, which had fewest variables in our dataset). (2) an un- balanced addition experiment where the numbers of variables added were proportional to the total number of the variables in each domain (i.e., 20 from ‘Biodiversity-Ecosystem Function’, 1 from ‘Ecosystem Services’, 1 from ‘Social-Economic Landscape’, 6 from ‘Global Change’). For each of the variable-adding experiments, we randomly chose the designated number of variables from each domain, and replicated this process 10,000 times. Once the variables for one causal analysis were set, we conducted causal analyses using 95% of the observations (or pixels) for 100 rounds of resampling.

We used linear non-Gaussian acyclic modeling (LiNGAM, implemented with the R-package ‘rlingam’: https://github.com/gkikuchi/rlingam) for causal analysis. Specifically, we used direct LiNGAM (62, 64). Direct LiNGAM was chosen because (1) our data were non-Gaussian distributed; (2) LiNGAM requires no prior knowledge on the causal directions and strengths as other conventional methods do; (3) our data were heterogenous, and direct LiNGAM requires no algorithmic parameters and is guaranteed to converge compared to ICA (independent component analysis)-based LiNGAM, which is not scale-invariant and may not always converge (64). The numbers and frequencies of positive, negative and neutral (absent) causal relationships were calculated for the original causal graphs and for each of the variable adding experiments (resampling threshold for causal relation, then counting). These were used to determine the probability of a positive, negative, or no causal relationship among each pair of variables, for each of the two directions between them.

Finally, we also tested whether the input variables were spatially autocorrelated, as this would potentially have an effect on LiNGAM outputs and the detection of unwarranted causal links. To investigate the impact of spatial autocorrelation on our spatially-based causal discovery, we conducted an analysis to determine the maximum distance at which the absolute value of spatial autocorrelation (Moran’s I) decreased to smaller than 0.1. Specifically, in each analysis, we treated one of the 227 variables as the dependent variable and the remaining covariates as independent variables to construct a linear model. We then calculated the changes in spatial autocorrelation from 0 to 500 km based on the linear model. Additionally, we calculated the pair- wise distances for all the 3289 observed points involved in the analysis. Overall, our analysis indicates that spatial autocorrelation has only a marginal effect on our spatial analysis. Results are reported in Supporting Information.

### Causal discovery simulation

We also conducted a simulation analysis to assess how well LiNGAM can retrieve correlations and causal relationships under a set of conditions likely to occur when studying SES relationships. To better understand the limitations of the LINGAM approach, we conducted simulations that varied 1) the strength of the causal relationships in a causal network (mean coefficient and standard deviation of the coefficient B), 2) the distribution (i.e., shape) of the variables, 3) scaling of the variables and 4) sample size. Detailed descriptions and the results can be found in Supporting Information.

## Acknowledgments

We thank B. Schmid, K. Shimizu, and O. Petchey, and participants of the URPP-GCB retreat workshop for their comments which improved the study and manuscript, and M. E. Schaepman for supporting the work through the NOMIS foundation project Remotely Sensing Ecological Genomics co-funding C.L., C.S., M.C.S., and M. dlH. We thank the University of Zurich, University Research Priority Program in Global Change and Biodiversity for supporting the project and funding P.Z., and the University of Zurich for supporting all authors.

## Author Contributions

CRediT taxonomy roles are listed with authors in alphabetical order. Conceptualization: D.Z.-D., M.C.S., M.B.E., M.J.S., P.Z., V.M.; Project Administration: M.C.S., M.J.S.; Supervision: F.P., M.C.S., M.J.S.; Data Curation: C.L., C.S., L.M., M.C.S., M.B.E., M.O.R., S.M., Y.T.; Formal Analysis: A.G., C.L., C.S., F.P., M.C.S., M.B.E., M.J.S., P.Z.; Funding Acquisition and Resources: M.C.S., M.J.S.; Investigation: A.G., C.L., C.S., D.Z.-D., F.P., M.C.S., M.D.L.H., M.B.E., M.J.S., P.Z., S.V.M., V.M., Y.T.; Methodology: A.G., F.P., M.C.S., M.B.E., M.J.S., P.Z.; Visualization: A.G., C.L., C.S., M.B.E., M.J.S., P.Z.; Writing - Original Draft: M.J.S., P.Z.; Writing - Review & Editing: A.G., C.L., C.S., D.Z.-D., F.P., M.C.S., M.D.L.H., M.B.E., M.J.S., P.Z., S.V.M., V.M., Y.T..

