## Supplementary Information for "Causal discovery in biodiversity-mediated social-ecological systems"

### Source data files and code

Data sources are available upon request to the authors and code is available at https://github.com/ESS-uzh/borneo_urppgcb.

**Supporting Information Text**

**PRISMA diagram for the literature review and additional analyses**

*Structure of the literature review:* We followed the PRISMA guidelines (https://www.prisma-statement.org/) for literature search and the resulting results and selection procedure is described in **Figure S1**.

For the final selection of articles to analyze (~7500) we coded each article according to defined domains and categories that describe the four fundamental domains of a SES (**Fig. 1**). We list these terms and describe them in **Table S1**.

*Randomization tests for literature review:* Focusing on studies covering multiple domains, the main text describes which combinations of domains were over- and under-represented. Over- and underrepresentation of each domain combination was determined using randomization tests (*3*). Specifically, we created 10,000 randomizations of the observed domain frequencies. This yielded for each domain combination an average number of occurrences, as well as the standard deviation of occurrences. The degree of over- or under-representation of a domain combination in the dataset can then be expressed as the standardized effect size:

$$SES = \frac{\theta_{obs} - \bar{\theta}_{rand}}{\sigma_{rand}}$$

In which $\theta_{obs}$ is the frequency of a domain combination in the dataset, $\bar{\theta}_{rand}$ is the average frequency at which this combination occurred in randomized replicates and $\sigma_{rand}$ its standard deviation. The probability of exceedance for observing the SES under the null hypothesis of randomly distributed domain frequencies can then be determined non-parametrically through the number of randomized replicates that exceed the (absolute value of the) observed effect size. In general, however, the standardized effect sizes > 2 indicate probabilities of exceedance below p = 0.05 (**Fig. S2**).

*Coding Consistency Analysis:* We conducted a coding consistency analysis (see description in Methods). We found a high degree of consistency on domain assignment by co-authors coding the articles retrieved by the literature search, and agreement on the number of domains assigned (**Fig. S3**).

**Literature used to support the example causal graphs**

We chose three example causal graphs based on our expert knowledge and supported by literature that examined the pair-wise relationships between each of the variables representing each of the four domains of the hypothesized SES for each of the causal graphs. In **Table S2** we provide the references used to justify the example causal graphs for the island of Borneo as well as a description of the analysis conducted in these studies, the direction of the relationship as reported and its strength.

**LiNGAM data, structure and additional results**

We used a set of 227 variables as input to LiNGAM for causal discovery in SES. The description of the data and sources is provided in **Table S3**. The data was prepared following the workflow described in **Fig. S4**.

*Spatial autocorrelation:* Our analysis indicates that 95% of the covariates decrease to an absolute value of spatial autocorrelation smaller than 0.1 at a distance of less than 23.4 km (**Fig. S5a**), while 97.5% of the distance pairs among the 3289 observations are longer than 26.7 km (**Fig. S5b**). Furthermore, the 12 variables used in the example causal graphs, decreased to an absolute spatial autocorrelation of less than 0.1 at a distance shorter than 9.8 km (range = 1.7-23.4 km).

In addition to the results in **Fig. 3** in the main text, we provide in **Fig. S6** the results of the balanced and unbalanced addition of variables to the core causal graphs we examined as examples. It can be observed that as more variables are added the causal links become more stable, i.e., similar results every time a new variable is added, but not always converging to a consensus in the causal direction. We also find that the unbalanced addition of new variables results in similar patterns. Further, in **Table S3** we provide the proportional details underlying the graphs presented in the LiNGAM results in **Fig. 3** in the main text.

**Simulation with LiNGAM**

To better understand the limitations of the LiNGAM approach, we conducted simulations that varied 1) the strength of the causal relationships in a causal network (mean coefficient and standard deviation of the coefficient B), 2) the distribution (i.e., shape) of the variables, 3) scaling of the variables and 4) sample size.

*Strength of the causal relationships in a causal network:* It is known that the power of LiNGAM to determine causal relationships is related to the strength of its coefficient B, which is not the same as, but related to, the Pearson correlation coefficient between two variables. By simulation, we found that a too-weak B (corresponding to a Pearson correlation coefficient of less than 0.25) or a too-strong B (corresponding to a Pearson correlation coefficient close to 1) can lead to frequent false causal predictions in networks where the causality is known (i.e., pre-determined for the sake of simulation; **Fig. S7**). We expect this is related to LiNGAM's reliance on differences in the error of correlation; if the error is much greater than the relationship perhaps no causal link can be determined, and if the relationship is much greater than the error the directionality may be difficult to determine. We accept the limitation of not being able to determine weak causal relationships, and given the nature of correlations in ecology and social sciences we do not expect a too-strong relationship to often be a problem.

Furthermore, we investigated the effect of variation in the coefficient B and the ability of LiNGAM to recover the causal network. Usually, higher variation in B led to a higher true positive/negative rate (**Fig. S7**). We explain this slightly counterintuitive result with the higher probability of having stronger values of B in networks with more variation, which LiNGAM is better able to detect.

*Shape of distribution:* LiNGAM showed high accuracy in uncovering causal links with bimodal, imperfect Gaussian and skewed variables (**Fig. S8**). Higher B coefficients resulted in higher accuracy for all three distribution shapes. The bimodal distribution led to higher variation in the true positive rate at high standard deviation of B. Since bimodal, imperfect Gaussian and skewed distributions make up the majority of the variable shapes in the empirical analysis, we are confident that LiNGAM should perform well.

*Scaling variables:* We encountered the unanticipated problem that inputting unscaled data often caused the LiNGAM analysis to fail. This was solved by scaling the data, initially done by dividing each dataset by its standard deviation (an imperfect approximation of the variance as these are non-

Gaussian distributions). This resulted in data on a similar scale, but simulations showed that this can result in completely different outcomes of causal analysis. In **Fig. S9** we show a comparison of the observed core causal graph (panel A) versus that estimated using unscaled (B) and scaled (C) data. We find that when using the unscaled data, the observed core causal graph is retrieved, while no causality or different direction of causality emerges when using scaled data (**Fig. S9c**), and the results of empirical analysis using scaled data were much less stable. Specifically, the results using scaled empirical data, like the simulation on scaled data, showed changes in direction, lower likelihood to identify causal relationships, and a disappearance of causality with inclusion of additional variables, in comparison to results from unscaled data, which tended to consistently identify likelihood of directional causality regardless of variable addition. These different outcomes could be related to changes in the modeled coefficient B resulting from scaling. In simulations wherein data which originally had a difference in scale were scaled to the same range, B became very small. We consider this artificially deflating B. However, when data have too great a difference in scale -- we estimate more than 1 order of magnitude -- the analysis fails for reasons we do not yet understand, but potentially related to too-large B. Based on our findings, we recommend to find out which variables cause LiNGAM to fail. If these are the variables with extreme differences in range compared to the core dataset, then (3) scale only the variables with an extreme range, by some minimum amount so as to allow LiNGAM to proceed; try to determine this minimum amount by simulation.

*Sample size:* As expected, larger sample size led to higher fractions of true positives and negatives. However, already with a sample size of five, almost 90% of the true positives and more than 99% of the true negatives were correctly recovered. This gives us confidence that with the sample size of our empirical analyses we should be able to uncover all but the weakest causal links.

**Word frequency of occurrence and causal graphs**

In the main text we present a subset of word frequency of occurrence and association and the results for the entire corpus are described in **Table S4**.

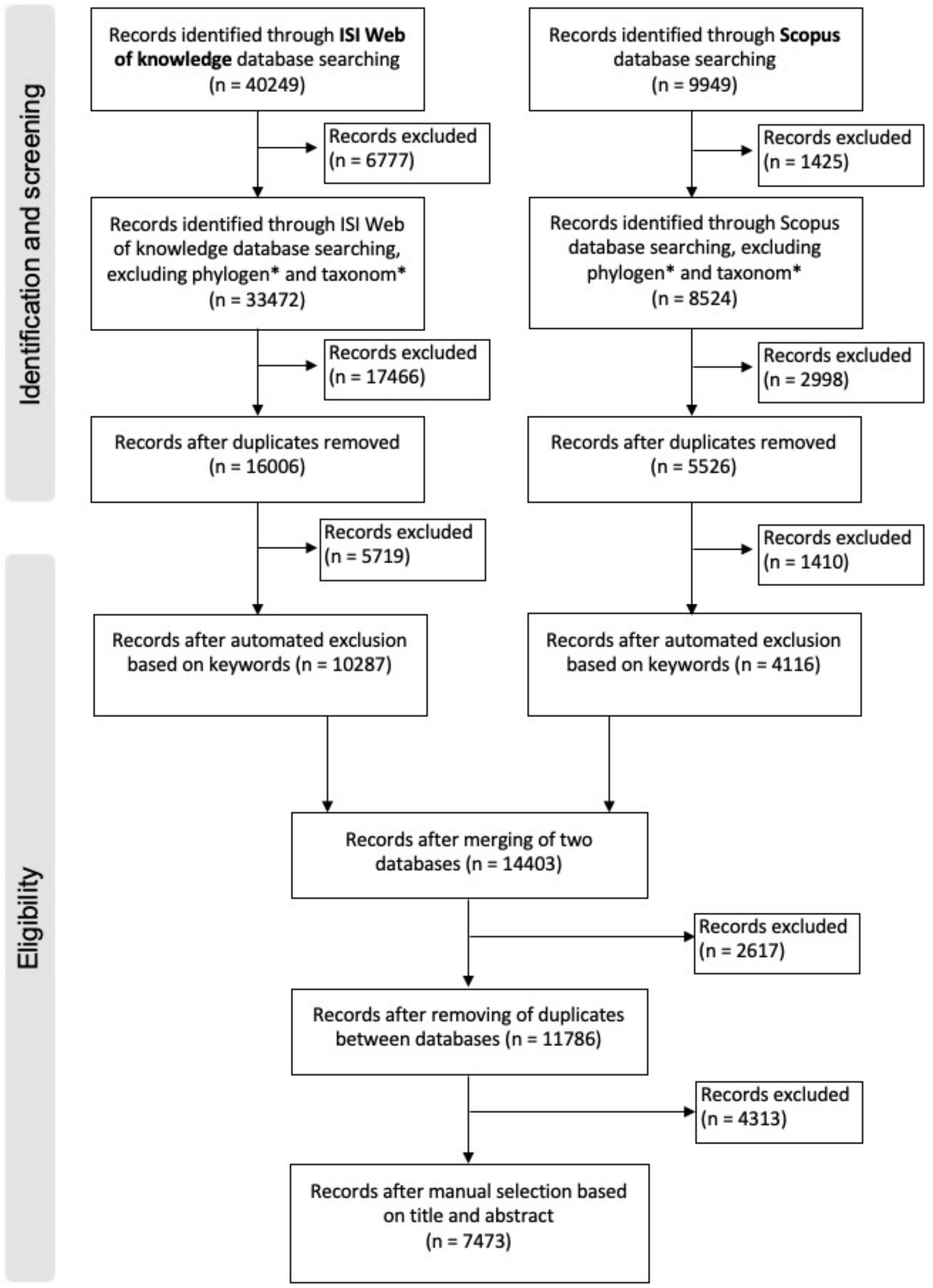


**Figure S1:** PRISMA diagram: Flow diagram of the systematic review, including search terms, results, dates, numbers, numbers removed based on criteria, etc. - usual flow chart. This flow diagram and systematic review path follows the recommendations by (*1*, *2*).

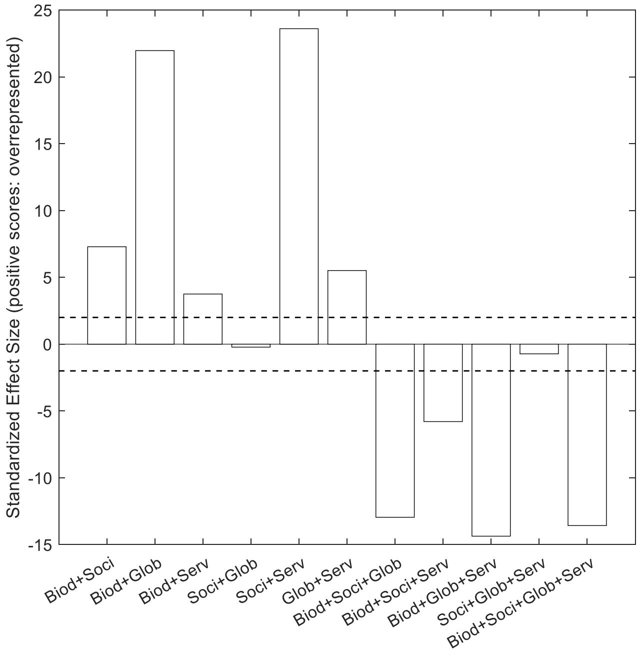


**Figure S2:** Standardized effect sizes indicating whether specific combinations of domains were overrepresented (positive scores) or underrepresented (negative scores) in the corpus of studies analyzed.

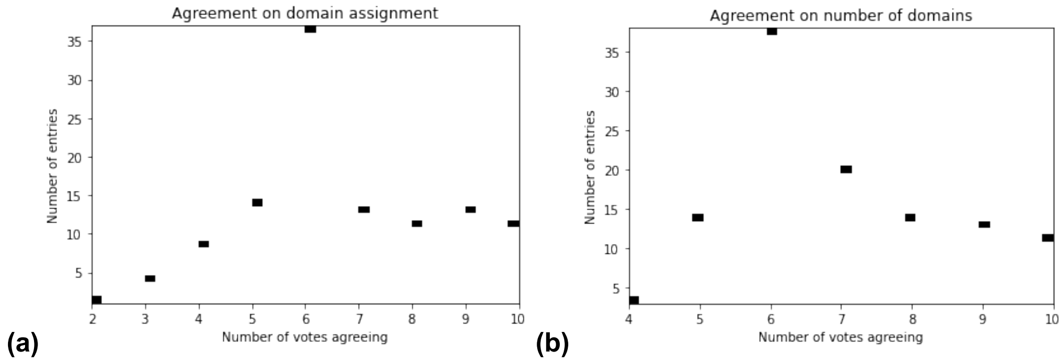


**Figure S3:** (a) Agreement on domain assignment by co-authors coding the articles retrieved by the literature search, and (b) agreement on the number of domains assigned.

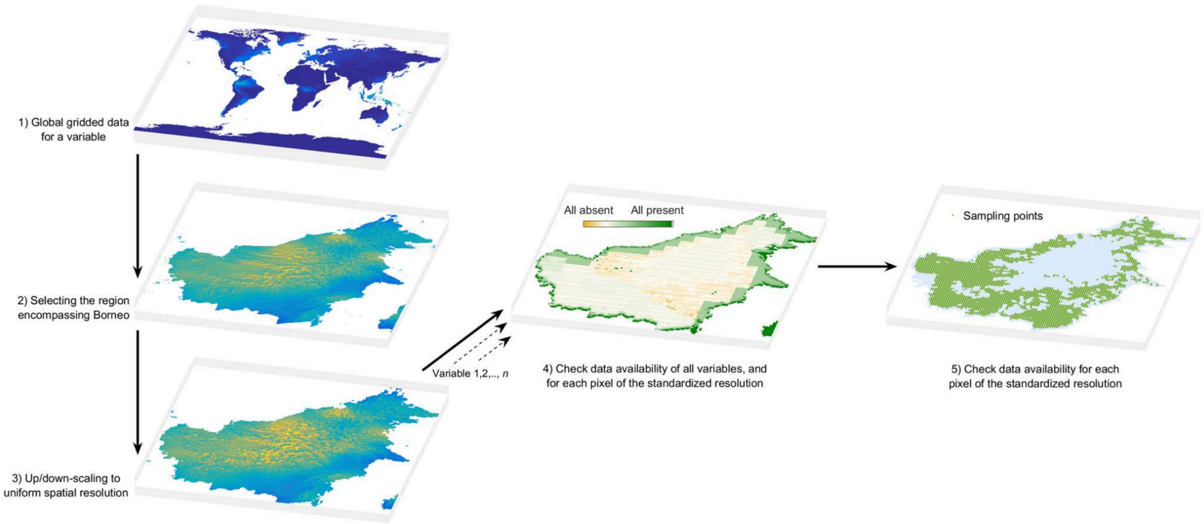


**Figure S4.** Workflow for data preparation for causal inference.

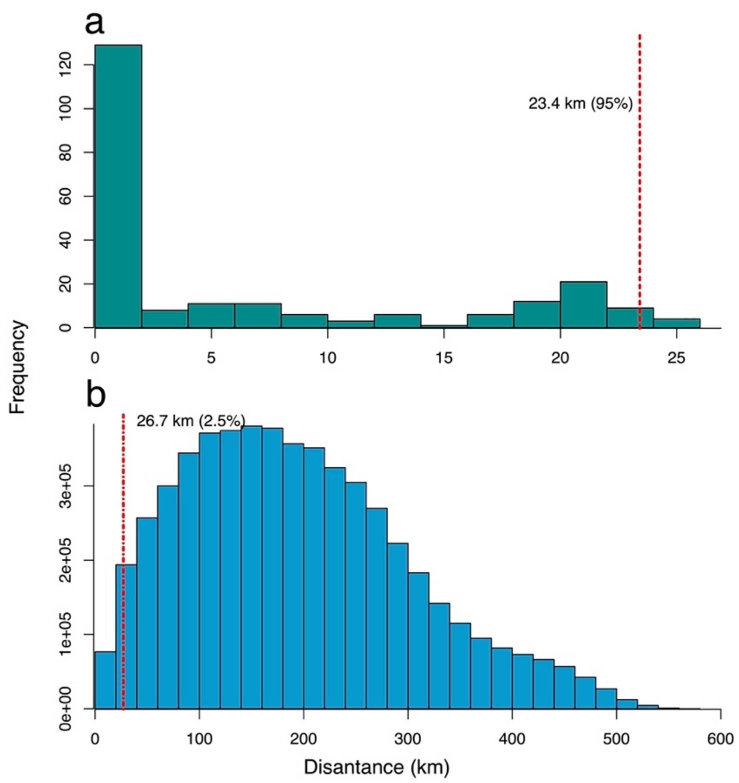


**Figure S5.** Spatial autocorrelation distribution of the 227 covariates and the spatial distances among the 3289 observations. **(a)** Distribution of distances at which spatial autocorrelation decreases to an absolute value smaller than 0.1, with the red dashed line indicating the 95% quantile at 23.4 km. **(b)** Distribution of distances among all 3289 observations used in our analysis, showing that 97.5% of the distances are greater than 26.7 km, which exceeds the 23.4 km threshold in panel **(a)**.

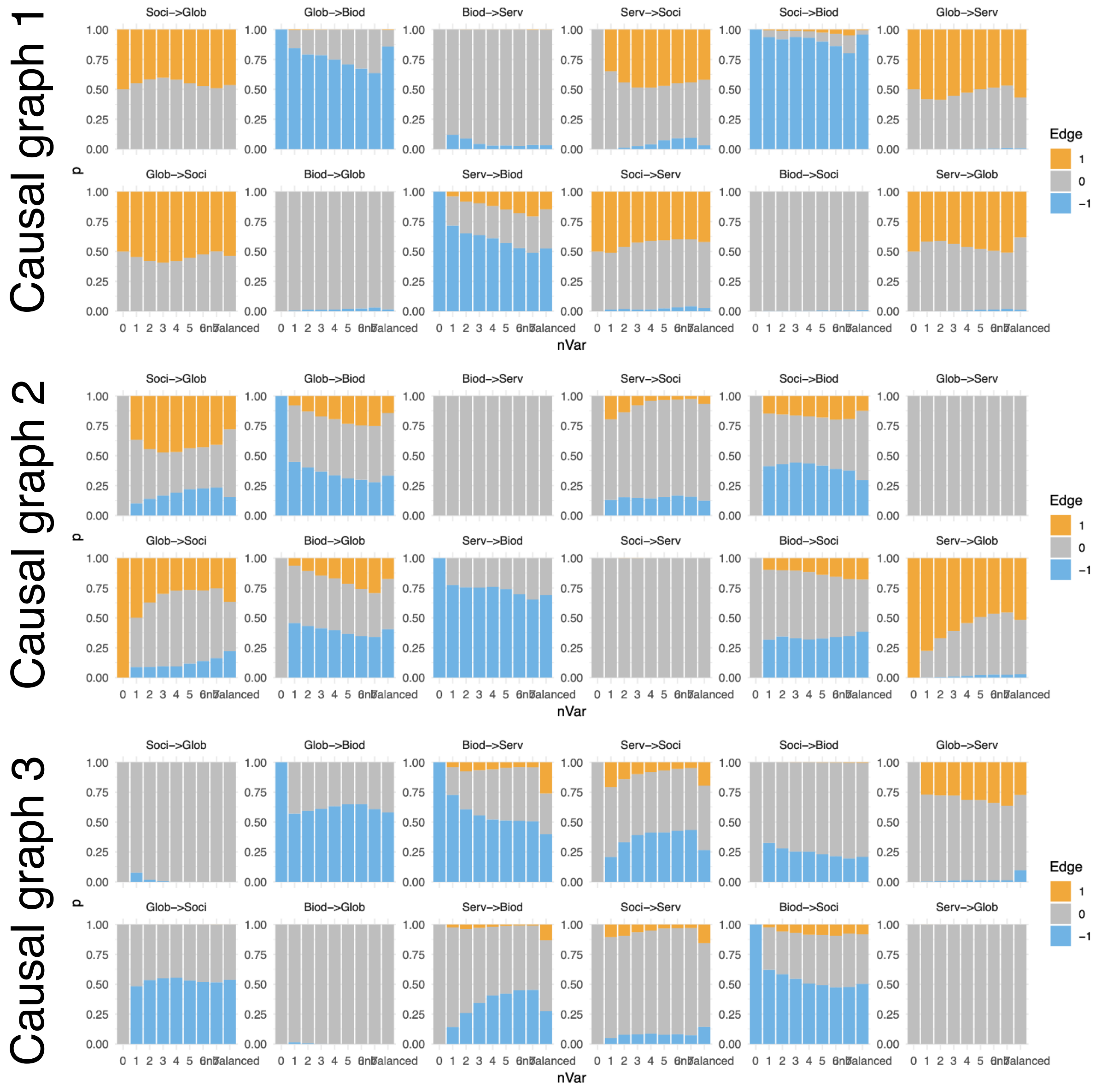


**Figure S6.** Causal analyses results when adding variables in a balanced (adding 1 up to 7 additional variables to the core causal graph) and unbalanced (randomly selecting among the 227 variables without considering the number of variables in each SES domain). Colors indicate positive, neutral or negative causal link.

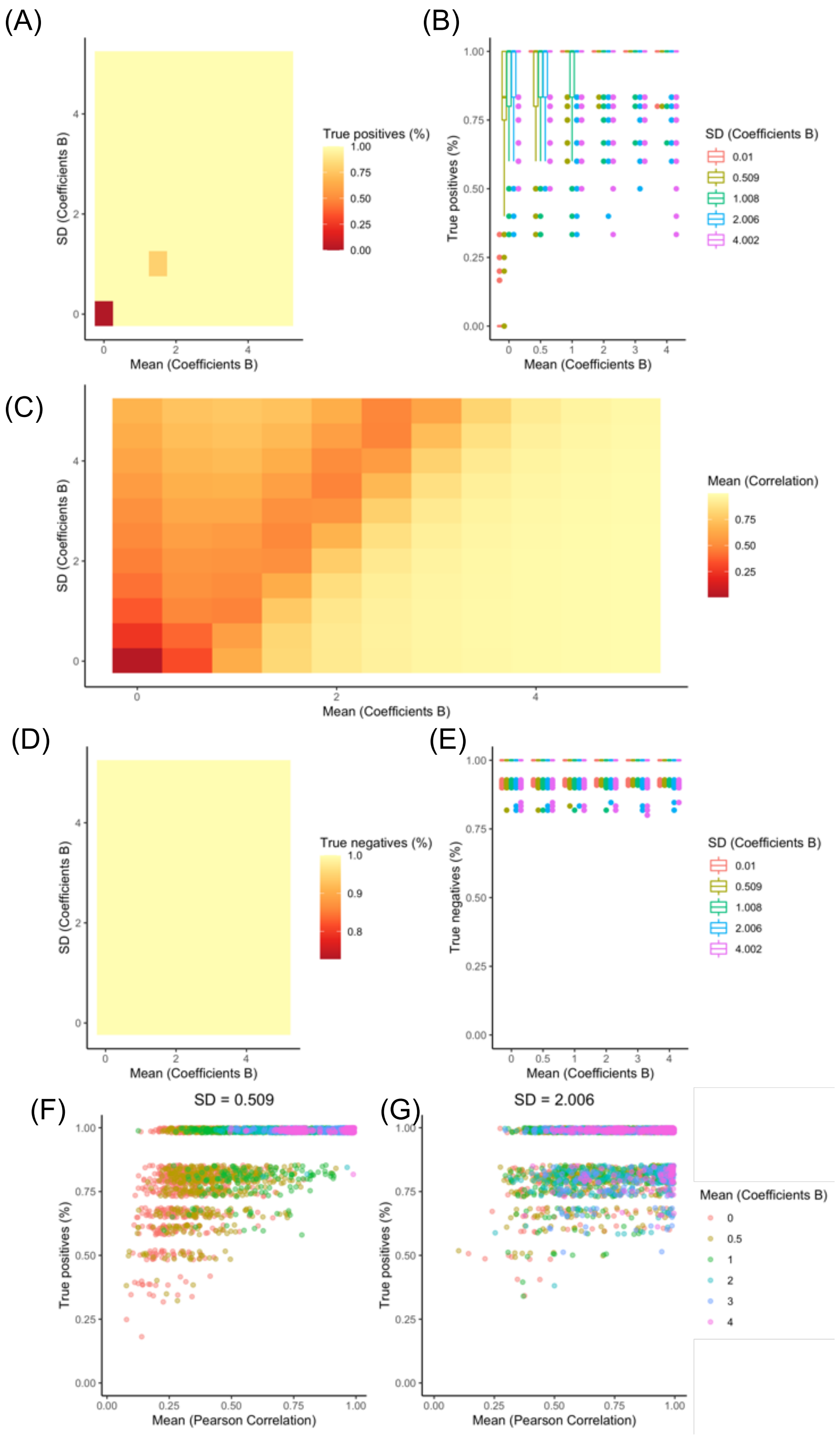


**Figure S7:** Results of the randomization experiments to test for the effects of the strength of the causal relationship for both true positives and negatives: (A) True positives as a function of the mean strength of B and its standard deviation (SD), (B) Fraction of true positives distributed by B coefficient ranges and their SD, (C) Correlation between the strength of the B coefficient and its variation in effect, (D) False positives as a function of the mean strength of B and its standard deviation (SD), (E) Fraction of false positives distributed by B coefficient ranges and their SD, (F) True positives as a function of Pearson Correlation at different values of Coefficient B for low SD, and (G) True positives as a function of Pearson Correlation at different values of Coefficient B for high SD.

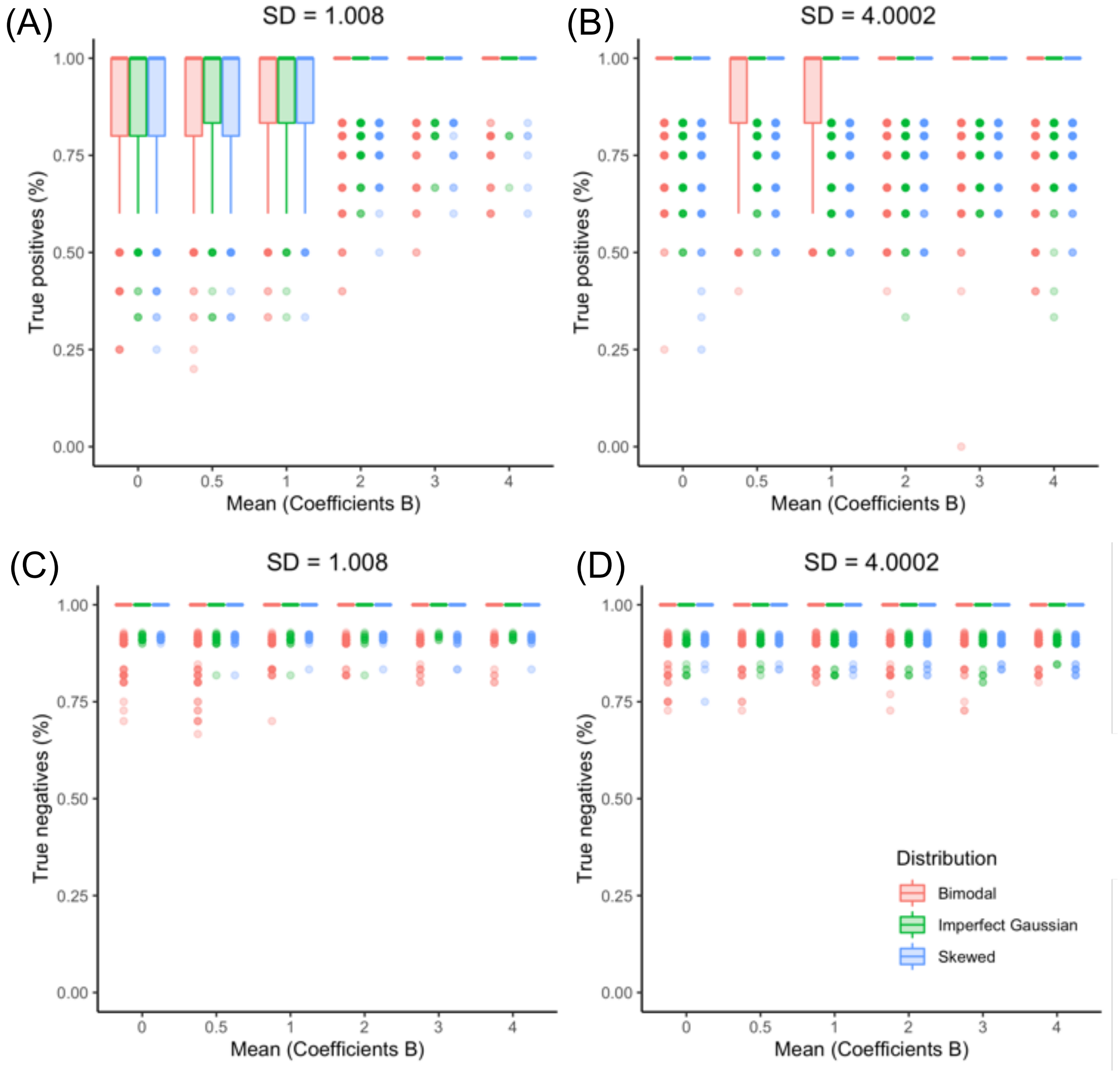


**Fig. S8:** The fraction of true positives and negatives are shown as a function of the mean strength of B and its standard deviation (SD) and the shape of the distributions (bimodal, imperfect Gaussian and skewed Gaussian).

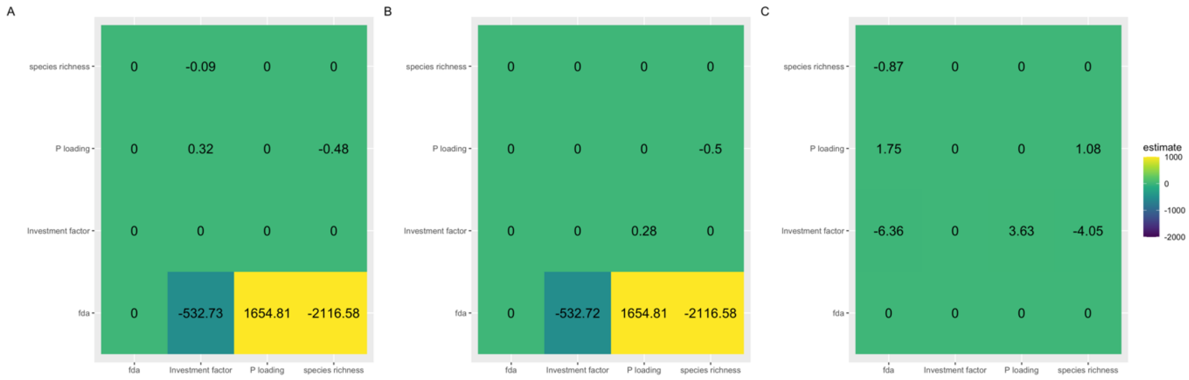


**Figure S9:** The effect of scaling on the estimated coefficient B: (A) The observed causal graph, (B) estimated causal graph when using simulated unscaled data, and (C) estimated causal graph when using simulated scaled data.

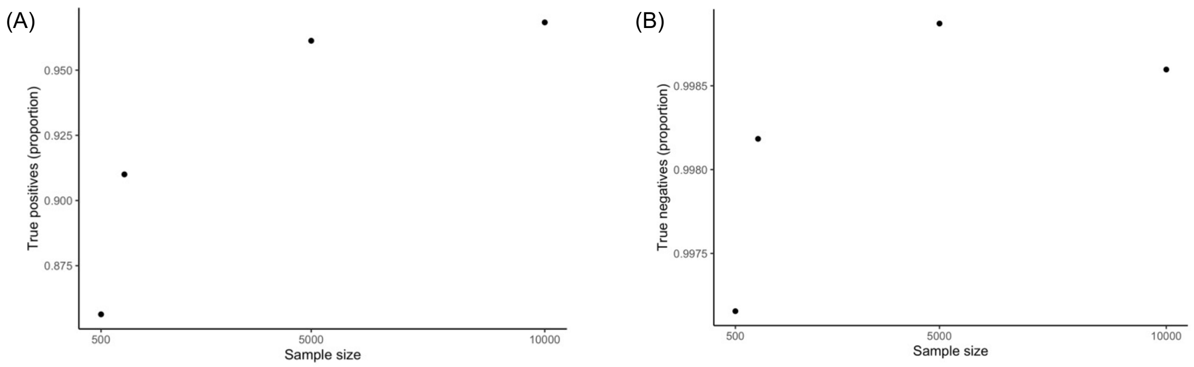


**Figure S10:** The effect of sample size on proportion of (A) true positives, and (B) true negatives.

**Table S1:** SES domains and categories for the manual selection of the publications to include in the research weaving analysis.

| | Score 1 (include) | | | | Score 0 (exclude) |
|---|---|---|---|---|---|
| **Domain** | **Social-Economic** | **Global Change** | **Biodiversity - Ecosystem Function** | **Ecosystem Services** | **Inappropriate** |
| **Description** | Human social and economic systems | Land- and sea-use change, direct exploitation of organisms, climate change, pollution, and species invasions | Species distribution, abundance, richness, and evenness of genotypes, species, other phylogenetic or taxonomic groups, or functional groups | Provisioning, regulating, supporting, and cultural services | Related to methodology, topic without connection to the four domains listed on the left, credibility of publication |
| **Examples** | Culture | Climate | Genetic composition-Co-ancestry | Habitat creation and maintenance | Methodology: |
| | Demographics | Soil | Genetic composition-Allelic diversity | Pollination and propagule dispersal | Captivity |
| | Health | Water | Population genetic differentiation | Regulation of air quality | Proof of principle |
| | Cultivation (e.g. agriculture, horticulture, silviculture) | Land use change (includes conservation) | Breed and variety diversity | Regulation of climate | Simulation |
| | Energy | Exploitation of organisms | Species distribution | Regulation of ocean acidification | Theory |
| | Technology and innovation | Climate change | Population abundance | Regulation of freshwater quantity, location, timing | Taxonomic revisions |
| | Economics (including poverty and economic development) | Pollution | Population structure by age/size class | Regulation of freshwater and coastal water quality | Anatomy |
| | Livelihoods | Invasions | Species traits-Phenology | Formation, protection, decontamination of soils and sediments | Geology (e.g. ancient tectonic plate movements, paleogeography, hydrology). |
| | Living conditions | | Species traits-Morphology | Regulation of hazards and extreme events | Wrong geographic location (not Borneo) |

| | | | | | |
|---|---|---|---|---|---|
| | Education and knowledge | | Species traits-Reproduction | Regulation of detrimental organisms and biological processes | Source (not credible, news article, not a scientific paper) |
| | Human rights | | Species traits-Physiology | Energy | Duplicate |
| | Violence and war | | Species traits-Movement | Food and feed | Corrigendum, Erratum, Expression of concern |
| | | | Taxonomic diversity | Materials, companionship and labor | Response paper |
| | | | Species interactions | Medicinal, biochemical and genetic resources | Review: exclusively reviews of existing literature that do not report new results based on e.g. synthesis or re-analysis of those results. |
| | | | Net primary productivity | Learning and inspiration | Conference proceedings, e.g. describing a meeting |
| | | | Secondary productivity | Physical and psychological experiences | Theory |
| | | | Nutrient retention | Supporting identities | Language (i.e., we are only considering English literature) |
| | | | Disturbance regime | Maintenance of options | Insufficient information (e.g. missing DOI, abstract) |
| | | | Habitat structure | | |
| | | | Ecosystem extent and fragmentation | | |
| | | | Ecosystem composition by functional type | | |

**Table S2.** Examples of SES with known causal relations informed by the literature that were used in our analyses to test the causal inference algorithm. We provide the references where causality between each pair of variables was described, and the strength and directionality of the relationship when possible.

| Causal graph | | Reference | Details |
|---|---|---|---|
| Causal graph 1 | 'Human Development' WITH 'Human Footprint' | (*4*) | What: Human development explains human footprint.<br>Strength: not reported<br>Direction: negative |
| | 'Human Footprint' WITH 'Tree density' | (*5*) | What: Reports changes in woodland area and wood cut density as a function of human footprint.<br>Strength: strong<br>Direction: negative |
| | 'Tree density' WITH 'Cropland' | (*6*) | What: Examines tree density recovery post cropland abandonment.<br>Strength: strong<br>Direction: negative |
| | 'Cropland' WITH 'Human development' | (*7*) | What: Links land use changes over Borneo to human development<br>Strength: strong<br>Direction: positive |
| Causal graph 2 | 'Investment benefit factor' WITH 'Phosphorus loading' | (*8*) | What: Shows that higher leaf phosphorus is linked to length of investment (Indirect effect on phosphorus loading)<br>Strength: not reported<br>Direction: positive |
| | 'Phosphorus loading' with 'Species richness' | (*9*, *10*) | What: Shows relationship between tropical forest diversity and phosphorus concentration/availability<br>Strength: strong<br>Direction: positive |
| | 'Species richness' WITH 'Food area' | (*11*) | What: Shows that ecosystem services depend on species richness<br>Strength: strong<br>Direction: negative |

| | | | |
|---|---|---|---|
| | ‘Food area’ WITH ‘Investment benefit factor’ | (*12*) | What: examine relationship between ecosystem services and investment<br>Strength: not reported<br>Direction: positive |
| Causal graph 3 | ‘Population density’ WITH ‘Forest loss’ | (*13*) | What: examine relationship of forest loss with population density<br>Strength: strong<br>Direction: positive |
| | ‘Forest loss’ WITH ‘alpha diversity’ | (*14*) | What: examine relationship of forest loss with diversity<br>Strength: strong<br>Direction: negative |
| | ‘alpha diversity’ WITH ‘soil water capacity’ | (*15*) | What: examine relationship between soil biota and soil properties<br>Strength: strong<br>Direction: both negative and positive |
| | ‘Soil water capacity’ WITH ‘Population density’ | (*16*) | What: examine flood risk and population parameters<br>Strength: weak<br>Direction: not reported |

**Table S3.** SES domains and variables available to include in the causal inference analysis**.**

| Variable name | Reference | Unit | Source | Description | Domain |
|---|---|---|---|---|---|
| Annual Soil Respiration | (*17*) | g C / m^2 * y | https://daac.ornl.gov/CMS/guides/CMS_Global_Soil_Respiration.html | (*17*) fitted a Quantile Regression Forest algorithm to over 2,500-point observations from the Soil Respiration Database and corresponding 1-km spatially distributed climate and vegetation covariates, of which were selected: mean annual temperature, mean annual precipitation, MODIS EVI and mean precipitation from November to January. | Biodiversity-Ecosystem Function |
| Percentage evergreen broadleaf trees | (*18*) | percentage (0-100) | https://www.earthenv.org/landcover | Percentage of the pixel area covered by evergreen broadleaf trees. (*18*) created a global 1-km scale consensus land cover product, integrating the DISCover (*19*), GLC2000 (*20*) (Mayaux et al. 2006), MODIS2005 (Friedl et al. 2010) and GlobCover (Bicheron et al. 2008) land cover products | Biodiversity-Ecosystem Function |
| Proportion of trees associating with arbuscular mycorrhizal fungi | (*21*) | Unitless (proportion) | https://static-content.springer.com/esm/art%3A10.1038%2Fs41586-019-1128-0/MediaObjects/41586_2019_1128_MOESM4_ESM.zip | (*21*) used forest inventory data to calculate the proportion of AM trees, and then created a global map using a model including as predictor variables: Decomposition of the Warmest quarter (using the Yasso07 model, which is a function of mean temperature and precipitation), Temperature seasonality, Decomposition of the wettest quarter and Temperature of the wettest quarter | Biodiversity-Ecosystem Function |
| Proportion of trees associating with ecto-mycorrhizal fungi | (*21*) | Unitless (proportion) | https://static-content.springer.com/esm/art%3A10.1038%2Fs41586-019-1128-0/MediaObjects/41586_2019_1128_MOESM4_ESM.zip | (*21*) used forest inventory data to calculate the proportion of AM trees, and then created a global map using a model including as predictor variables: Decomposition of the Warmest quarter (using the Yasso07 model, which is a function of mean temperature and precipitation), Temperature seasonality, Temperature range and Decomposition of the wettest quarter | Biodiversity-Ecosystem Function |

| | | | | | |
|---|---|---|---|---|---|
| Proportion of nitrogen-fixing trees | (*21*) | Unitless (proportion) | https://static-content.springer.com/esm/art%3A10.1038%2Fs41586-019-1128-0/MediaObjects/41586_2019_1128_MOESM4_ESM.zip | (*21*) used forest inventory data to calculate the proportion of AM trees, and then created a global map using a model including as predictor variables: Maximum temperature, Soil PH (in H2O), Temperature of the Warmest quarter and Temperature of the wettest quarter | Biodiversity-Ecosystem Function |
| Density of bacterivorous nematodes in the soil | (*22*) | Number/ 100 g soil | https://gitlab.ethz.ch/devinrouth/crowther_lab_nematodes | (*22*) compiled nematode compositions from a global dataset of soil samples, then created a global map using an iterative process that included 73 environmental co-variates, the most important ones being: Sand content (15cm), Organic CStock5-15cm, CEC 15 cm, MODIS Nadir reflectance bands 1 and 7, Annual precipitation, pH (15 cm), Aridity index, Temperature seasonality and Precipitation seasonality | Biodiversity-Ecosystem Function |
| Density of fungivorous nematodes in the soil | (*22*) | Number/ 100 g soil | https://gitlab.ethz.ch/devinrouth/crowther_lab_nematodes | (*22*) compiled nematode compositions from a global dataset of soil samples, then created a global map using an iterative process that included 73 environmental co-variates, the most important ones being: Sand content (15cm), Organic CStock5-15cm, CEC 15 cm, MODIS Nadir reflectance bands 1 and 7, Annual precipitation, pH (15 cm), Aridity index, Temperature seasonality and Precipitation seasonality | Biodiversity-Ecosystem Function |
| Density of herbivorous nematodes in the soil | (*22*) | Number/ 100 g soil | https://gitlab.ethz.ch/devinrouth/crowther_lab_nematodes | (*22*) compiled nematode compositions from a global dataset of soil samples, then created a global map using an iterative process that included 73 environmental co-variates, the most important ones being: Sand content (15cm), Organic CStock5-15cm, CEC 15 cm, MODIS Nadir reflectance bands 1 and 7, Annual precipitation, pH (15 cm), Aridity index, Temperature seasonality and Precipitation seasonality | Biodiversity-Ecosystem Function |
| Density of omnivorous nematodes in the soil | (*22*) | Number/ 100 g soil | https://gitlab.ethz.ch/devinrouth/crowther_lab_nematodes | (*22*) compiled nematode compositions from a global dataset of soil samples, then created a global map using an iterative process that included 73 environmental co-variates, the most important ones being: Sand content (15cm), Organic CStock5-15cm, CEC 15 cm, MODIS Nadir | Biodiversity-Ecosystem Function |

| | | | | | |
|---|---|---|---|---|---|
| | | | | reflectance bands 1 and 7, Annual precipitation, pH (15 cm), Aridity index, Temperature seasonality and Precipitation seasonality | |
| Density of predatory nematodes in the soil | (*22*) | Number/ 100 g soil | https://gitlab.ethz.ch/devinrouth/crowther_lab_nematodes | (*22*) compiled nematode compositions from a global dataset of soil samples, then created a global map using an iterative process that included 73 environmental co-variates, the most important ones being: Sand content (15cm), Organic CStock5-15cm, CEC 15 cm, MODIS Nadir reflectance bands 1 and 7, Annual precipitation, pH (15 cm), Aridity index, Temperature seasonality and Precipitation seasonality | Biodiversity-Ecosystem Function |
| Total density of nematodes in the soil | (*22*) | Number/ 100 g soil | https://gitlab.ethz.ch/devinrouth/crowther_lab_nematodes | (*22*) compiled nematode compositions from a global dataset of soil samples, then created a global map using an iterative process that included 73 environmental co-variates, the most important ones being: Sand content (15cm), Organic CStock5-15cm, CEC 15 cm, MODIS Nadir reflectance bands 1 and 7, Annual precipitation, pH (15 cm), Aridity index, Temperature seasonality and Precipitation seasonality | Biodiversity-Ecosystem Function |
| Soil Carbon to Nitrogen ratio from 0.00m-0.05m | (*23*) | g.g^-1 | https://www.soilgrids.org ; https://www.isric.org/explore/wosis/accessing-wosis-derived-datasets | (*23*) used data on 150,000 soil profiles worldwide to create a global map, integrating observations within profile to obtain datapoints at fixed depth.A machine learning approach was used, with input data on: Digital Elevation Model-derived characteristics (using the merged DEMS SRTMGL3 and GMTED, as described in (*24*)), long-term average and sd of MODIS-based EVI, bands 4 and 7, long-term monthly average and standard deviation in land surface temperature (MODIS LST), long-term averaged mean monthly hours of snow cover (MODIS 8-day snow occurrence), land cover (using GlobCover30, described in (*25*), who classified Landsat data from 2000 and 2010), precipitation data (averaging WorldClim and GPCP), lithology (using GLiM, (*26*)), Global water table depth (*27*), long-term averaged mean monthly MODIS Flood Water, Landsat-based distribution of mangroves (*28*) and average soil and sedimentary-deposit thickness (*29*) | Biodiversity-Ecosystem Function |
| Soil Carbon to Nitrogen ratio from 0.00m-2.00m | (*23*) | g.g^-1 | https://www.soilgrids.org ; https://www.isric.org/explore/wosis/accessing-wosis- | (*23*) used data on 150,000 soil profiles worldwide to create a global map, integrating observations within profile to obtain datapoints at fixed depth.A machine learning approach was used, with input data on: Digital Elevation Model-derived characteristics (using the merged DEMS SRTMGL3 and GMTED, as described in (*24*)), long-term average and sd of MODIS-based EVI, bands 4 and 7, long-term monthly average and standard | Biodiversity-Ecosystem Function |

| | | | | | |
|---|---|---|---|---|---|
| | | | derived-datasets | deviation in land surface temperature (MODIS LST), long-term averaged mean monthly hours of snow cover (MODIS 8-day snow occurrence), land cover (using GlobCover30, described in (*25*), who classified Landsat data from 2000 and 2010), precipitation data (averaging WorldClim and GPCP), lithology (using GLiM, (*26*)), Global water table depth (*27*), long-term averaged mean monthly MODIS Flood Water, Landsat-based distribution of mangroves (*28*) and average soil and sedimentary-deposit thickness (*29*) | |
| Soil Carbon to Nitrogen ratio from 0.05m-0.15m | (*23*) | g.g^-1 | https://www.soilgrids.org ; https://www.isric.org/explore/wosis/accessing-wosis-derived-datasets | (*23*) used data on 150,000 soil profiles worldwide to create a global map, integrating observations within profile to obtain datapoints at fixed depth.A machine learning approach was used, with input data on: Digital Elevation Model-derived characteristics (using the merged DEMS SRTMGL3 and GMTED, as described in (*24*)), long-term average and sd of MODIS-based EVI, bands 4 and 7, long-term monthly average and standard deviation in land surface temperature (MODIS LST), long-term averaged mean monthly hours of snow cover (MODIS 8-day snow occurrence), land cover (using GlobCover30, described in (*25*), who classified Landsat data from 2000 and 2010), precipitation data (averaging WorldClim and GPCP), lithology (using GLiM, (*26*)), Global water table depth (*27*), long-term averaged mean monthly MODIS Flood Water, Landsat-based distribution of mangroves (*28*) and average soil and sedimentary-deposit thickness (*29*) | Biodiversity-Ecosystem Function |
| Soil Carbon to Nitrogen ratio from 0.15m-0.30m | (*23*) | g.g^-1 | https://www.soilgrids.org ; https://www.isric.org/explore/wosis/accessing-wosis-derived-datasets | (*23*) used data on 150,000 soil profiles worldwide to create a global map, integrating observations within profile to obtain datapoints at fixed depth.A machine learning approach was used, with input data on: Digital Elevation Model-derived characteristics (using the merged DEMS SRTMGL3 and GMTED, as described in (*24*)), long-term average and sd of MODIS-based EVI, bands 4 and 7, long-term monthly average and standard deviation in land surface temperature (MODIS LST), long-term averaged mean monthly hours of snow cover (MODIS 8-day snow occurrence), land cover (using GlobCover30, described in (*25*), who classified Landsat data from 2000 and 2010), precipitation data (averaging WorldClim and GPCP), lithology (using GLiM, (*26*)), Global water table depth (*27*), long-term averaged mean monthly MODIS Flood Water, Landsat-based | Biodiversity-Ecosystem Function |

| | | | | | |
|---|---|---|---|---|---|
| | | | | distribution of mangroves (*28*) and average soil and sedimentary-deposit thickness (*29*) | |
| Soil Carbon to Nitrogen ratio from 0.30m-0.60m | (*23*) | g.g^-1 | https://www.soilgrids.org ; https://www.isric.org/explore/wosis/accessing-wosis-derived-datasets | (*23*) used data on 150,000 soil profiles worldwide to create a global map, integrating observations within profile to obtain datapoints at fixed depth.A machine learning approach was used, with input data on: Digital Elevation Model-derived characteristics (using the merged DEMS SRTMGL3 and GMTED, as described in (*24*)), long-term average and sd of MODIS-based EVI, bands 4 and 7, long-term monthly average and standard deviation in land surface temperature (MODIS LST), long-term averaged mean monthly hours of snow cover (MODIS 8-day snow occurrence), land cover (using GlobCover30, described in (*25*), who classified Landsat data from 2000 and 2010), precipitation data (averaging WorldClim and GPCP), lithology (using GLiM, (*26*)), Global water table depth (*27*), long-term averaged mean monthly MODIS Flood Water, Landsat-based distribution of mangroves (*28*) and average soil and sedimentary-deposit thickness (*29*) | Biodiversity-Ecosystem Function |
| Soil Carbon to Nitrogen ratio from 0.60m-1.00m | (*23*) | g.g^-1 | https://www.soilgrids.org ; https://www.isric.org/explore/wosis/accessing-wosis-derived-datasets | (*23*) used data on 150,000 soil profiles worldwide to create a global map, integrating observations within profile to obtain datapoints at fixed depth.A machine learning approach was used, with input data on: Digital Elevation Model-derived characteristics (using the merged DEMS SRTMGL3 and GMTED, as described in (*24*)), long-term average and sd of MODIS-based EVI, bands 4 and 7, long-term monthly average and standard deviation in land surface temperature (MODIS LST), long-term averaged mean monthly hours of snow cover (MODIS 8-day snow occurrence), land cover (using GlobCover30, described in (*25*), who classified Landsat data from 2000 and 2010), precipitation data (averaging WorldClim and GPCP), lithology (using GLiM, (*26*)), Global water table depth (*27*), long-term averaged mean monthly MODIS Flood Water, Landsat-based distribution of mangroves (*28*) and average soil and sedimentary-deposit thickness (*29*) | Biodiversity-Ecosystem Function |
| Soil Carbon to Nitrogen ratio from 1.00m-2.00m | (*23*) | g.g^-1 | https://www.soilgrids.org ; https://www.isric.org/explore/wosis/accessin | (*23*) used data on 150,000 soil profiles worldwide to create a global map, integrating observations within profile to obtain datapoints at fixed depth.A machine learning approach was used, with input data on: Digital Elevation Model-derived characteristics (using the merged DEMS SRTMGL3 and GMTED, as described in (*24*)), long-term average and sd of MODIS- | Biodiversity-Ecosystem Function |

| | | | | | |
|---|---|---|---|---|---|
| | | | g-wosis-derived-datasets | based EVI, bands 4 and 7, long-term monthly average and standard deviation in land surface temperature (MODIS LST), long-term averaged mean monthly hours of snow cover (MODIS 8-day snow occurrence), land cover (using GlobCover30, described in (*25*), who classified Landsat data from 2000 and 2010), precipitation data (averaging WorldClim and GPCP), lithology (using GLiM, (*26*)), Global water table depth (*27*), long-term averaged mean monthly MODIS Flood Water, Landsat-based distribution of mangroves (*28*) and average soil and sedimentary-deposit thickness (*29*) | |
| Soil Nitrogen content from 0.00m-2.00m | (*23*) | g.kg^-1 | https://www.soilgrids.org ; https://www.isric.org/explore/wosis/accessing-wosis-derived-datasets | (*23*) used data on 150,000 soil profiles worldwide to create a global map, integrating observations within profile to obtain datapoints at fixed depth.A machine learning approach was used, with input data on: Digital Elevation Model-derived characteristics (using the merged DEMS SRTMGL3 and GMTED, as described in (*24*)), long-term average and sd of MODIS-based EVI, bands 4 and 7, long-term monthly average and standard deviation in land surface temperature (MODIS LST), long-term averaged mean monthly hours of snow cover (MODIS 8-day snow occurrence), land cover (using GlobCover30, described in (*25*), who classified Landsat data from 2000 and 2010), precipitation data (averaging WorldClim and GPCP), lithology (using GLiM, (*26*)), Global water table depth (*27*), long-term averaged mean monthly MODIS Flood Water, Landsat-based distribution of mangroves (*28*) and average soil and sedimentary-deposit thickness (*29*) | Biodiversity-Ecosystem Function |
| Soil Nitrogen content from 0.00m-0.05m | (*23*) | g.kg^-1 | https://www.soilgrids.org ; https://www.isric.org/explore/wosis/accessing-wosis-derived-datasets | (*23*) used data on 150,000 soil profiles worldwide to create a global map, integrating observations within profile to obtain datapoints at fixed depth.A machine learning approach was used, with input data on: Digital Elevation Model-derived characteristics (using the merged DEMS SRTMGL3 and GMTED, as described in (*24*)), long-term average and sd of MODIS-based EVI, bands 4 and 7, long-term monthly average and standard deviation in land surface temperature (MODIS LST), long-term averaged mean monthly hours of snow cover (MODIS 8-day snow occurrence), land cover (using GlobCover30, described in (*25*), who classified Landsat data from 2000 and 2010), precipitation data (averaging WorldClim and GPCP), lithology (using GLiM, (*26*)), Global water table depth (*27*), long-term averaged mean monthly MODIS Flood Water, Landsat-based | Biodiversity-Ecosystem Function |

| | | | | | |
|---|---|---|---|---|---|
| | | | | distribution of mangroves (*28*) and average soil and sedimentary-deposit thickness (*29*) | |
| Soil Nitrogen content from 0.05m-0.15m | (*23*) | g.kg^-1 | https://www.soilgrids.org ; https://www.isric.org/explore/wosis/accessing-wosis-derived-datasets | (*23*) used data on 150,000 soil profiles worldwide to create a global map, integrating observations within profile to obtain datapoints at fixed depth.A machine learning approach was used, with input data on: Digital Elevation Model-derived characteristics (using the merged DEMS SRTMGL3 and GMTED, as described in (*24*)), long-term average and sd of MODIS-based EVI, bands 4 and 7, long-term monthly average and standard deviation in land surface temperature (MODIS LST), long-term averaged mean monthly hours of snow cover (MODIS 8-day snow occurrence), land cover (using GlobCover30, described in (*25*), who classified Landsat data from 2000 and 2010), precipitation data (averaging WorldClim and GPCP), lithology (using GLiM, (*26*)), Global water table depth (*27*), long-term averaged mean monthly MODIS Flood Water, Landsat-based distribution of mangroves (*28*) and average soil and sedimentary-deposit thickness (*29*) | Biodiversity-Ecosystem Function |
| Soil Nitrogen content from 0.15m-0.30m | (*23*) | g.kg^-1 | https://www.soilgrids.org ; https://www.isric.org/explore/wosis/accessing-wosis-derived-datasets | (*23*) used data on 150,000 soil profiles worldwide to create a global map, integrating observations within profile to obtain datapoints at fixed depth.A machine learning approach was used, with input data on: Digital Elevation Model-derived characteristics (using the merged DEMS SRTMGL3 and GMTED, as described in (*24*)), long-term average and sd of MODIS-based EVI, bands 4 and 7, long-term monthly average and standard deviation in land surface temperature (MODIS LST), long-term averaged mean monthly hours of snow cover (MODIS 8-day snow occurrence), land cover (using GlobCover30, described in (*25*), who classified Landsat data from 2000 and 2010), precipitation data (averaging WorldClim and GPCP), lithology (using GLiM, (*26*)), Global water table depth (*27*), long-term averaged mean monthly MODIS Flood Water, Landsat-based distribution of mangroves (*28*) and average soil and sedimentary-deposit thickness (*29*) | Biodiversity-Ecosystem Function |
| Soil Nitrogen content from 0.30m-0.60m | (*23*) | g.kg^-1 | https://www.soilgrids.org ; https://www.isric.org/explore/wosis/accessin | (*23*) used data on 150,000 soil profiles worldwide to create a global map, integrating observations within profile to obtain datapoints at fixed depth.A machine learning approach was used, with input data on: Digital Elevation Model-derived characteristics (using the merged DEMS SRTMGL3 and GMTED, as described in (*24*)), long-term average and sd of MODIS- | Biodiversity-Ecosystem Function |

| | | | | | |
|---|---|---|---|---|---|
| | | | g-wosis-derived-datasets | based EVI, bands 4 and 7, long-term monthly average and standard deviation in land surface temperature (MODIS LST), long-term averaged mean monthly hours of snow cover (MODIS 8-day snow occurrence), land cover (using GlobCover30, described in (*25*), who classified Landsat data from 2000 and 2010), precipitation data (averaging WorldClim and GPCP), lithology (using GLiM, (*26*)), Global water table depth (*27*), long-term averaged mean monthly MODIS Flood Water, Landsat-based distribution of mangroves (*28*) and average soil and sedimentary-deposit thickness (*29*) | |
| Soil Nitrogen content from 0.60m-1.00m | (*23*) | g.kg^-1 | https://www.soilgrids.org ; https://www.isric.org/explore/wosis/accessing-wosis-derived-datasets | (*23*) used data on 150,000 soil profiles worldwide to create a global map, integrating observations within profile to obtain datapoints at fixed depth.A machine learning approach was used, with input data on: Digital Elevation Model-derived characteristics (using the merged DEMS SRTMGL3 and GMTED, as described in (*24*)), long-term average and sd of MODIS-based EVI, bands 4 and 7, long-term monthly average and standard deviation in land surface temperature (MODIS LST), long-term averaged mean monthly hours of snow cover (MODIS 8-day snow occurrence), land cover (using GlobCover30, described in (*25*), who classified Landsat data from 2000 and 2010), precipitation data (averaging WorldClim and GPCP), lithology (using GLiM, (*26*)), Global water table depth (*27*), long-term averaged mean monthly MODIS Flood Water, Landsat-based distribution of mangroves (*28*) and average soil and sedimentary-deposit thickness (*29*) | Biodiversity-Ecosystem Function |
| Soil Nitrogen content from 1.00m-2.00m | (*23*) | g.kg^-1 | https://www.soilgrids.org ; https://www.isric.org/explore/wosis/accessing-wosis-derived-datasets | (*23*) used data on 150,000 soil profiles worldwide to create a global map, integrating observations within profile to obtain datapoints at fixed depth.A machine learning approach was used, with input data on: Digital Elevation Model-derived characteristics (using the merged DEMS SRTMGL3 and GMTED, as described in (*24*)), long-term average and sd of MODIS-based EVI, bands 4 and 7, long-term monthly average and standard deviation in land surface temperature (MODIS LST), long-term averaged mean monthly hours of snow cover (MODIS 8-day snow occurrence), land cover (using GlobCover30, described in (*25*), who classified Landsat data from 2000 and 2010), precipitation data (averaging WorldClim and GPCP), lithology (using GLiM, (*26*)), Global water table depth (*27*), long-term averaged mean monthly MODIS Flood Water, Landsat-based | Biodiversity-Ecosystem Function |

| | | | | | |
|---|---|---|---|---|---|
| | | | | distribution of mangroves (*28*) and average soil and sedimentary-deposit thickness (*29*) | |
| Number of trees per km^2 | (*30*) | # trees km$^{-2}$ | https://www.nature.com/articles/sdata20166 9#Sec12 | (*30*) used ~ 430,000 observations of tree density, to create generalized linear models of tree density for each biome, including as predictors topographic variables, climatic variables (Temperature seasonality, Mean annual temperature, Annual precipitation, Precipitation in driest quarter, Precipitation in driest month, Precipitation seasonality, Aridity and Evapotranspiration) and Vegetation indicators (LAI, EVI (including the texture EVI-based metrics: Angular second moment, contrast and dissimilarity)) | Biodiversity-Ecosystem Function |
| Occurrence of tropical montane cloud forests | (*31*) | Relative occurrence rate (ranging between 0 and 1) | https://www.earthenv.org/cloud | Relative occurrence rate of tropical montane cloud forests estimated using an inhomogeneous point process model. (*31*) used 15 years of MODIS data (the MOD09 atmospherically corrected reflectance product) to create a global dataset of the relative occurrence rate of clouds per pixel | Biodiversity-Ecosystem Function |
| Proportion of trees that is between 11 and 20 years old | (*32*) | Unitless (proportion) | https://doi.pangaea.de/10.1594/PANGAEA.889943 | Four Plant Functional Types from MODIS Collection 5.1 land cover dataset are used, crosswalking land cover types to PFT fractions. Age distributions are based on country-level forest inventories for temperate and high latitude regions, and based on biomass for tropical regions. | Biodiversity-Ecosystem Function |
| Proportion of trees that is between 21 and 30 years old | (*32*) | Unitless (proportion) | https://doi.pangaea.de/10.1594/PANGAEA.889943 | Four Plant Functional Types from MODIS Collection 5.1 land cover dataset are used, crosswalking land cover types to PFT fractions. Age distributions are based on country-level forest inventories for temperate and high latitude regions, and based on biomass for tropical regions. | Biodiversity-Ecosystem Function |
| Proportion of trees that is between 31 and 40 years old | (*32*) | Unitless (proportion) | https://doi.pangaea.de/10.1594/PANGAEA.889943 | Four Plant Functional Types from MODIS Collection 5.1 land cover dataset are used, crosswalking land cover types to PFT fractions. Age distributions are based on country-level forest inventories for temperate and high latitude regions, and based on biomass for tropical regions. | Biodiversity-Ecosystem Function |
| Proportion of trees that is between 41 and 50 years old | (*32*) | Unitless (proportion) | https://doi.pangaea.de/10.1594/PANGAEA.889943 | Four Plant Functional Types from MODIS Collection 5.1 land cover dataset are used, crosswalking land cover types to PFT fractions. Age distributions are based on country-level forest inventories for temperate and high latitude regions, and based on biomass for tropical regions. | Biodiversity-Ecosystem Function |

| | | | | | |
|---|---|---|---|---|---|
| Proportion of trees that is between 51 and 60 years old | (*32*) | Unitless (proportion) | https://doi.pangaea.de/10.1594/PANGAEA.889943 | Four Plant Functional Types from MODIS Collection 5.1 land cover dataset are used, crosswalking land cover types to PFT fractions. Age distributions are based on country-level forest inventories for temperate and high latitude regions, and based on biomass for tropical regions. | Biodiversity-Ecosystem Function |
| Proportion of trees that is between 61 and 70 years old | (*32*) | Unitless (proportion) | https://doi.pangaea.de/10.1594/PANGAEA.889943 | Four Plant Functional Types from MODIS Collection 5.1 land cover dataset are used, crosswalking land cover types to PFT fractions. Age distributions are based on country-level forest inventories for temperate and high latitude regions, and based on biomass for tropical regions. | Biodiversity-Ecosystem Function |
| Proportion of trees that is between 71 and 80 years old | (*32*) | Unitless (proportion) | https://doi.pangaea.de/10.1594/PANGAEA.889943 | Four Plant Functional Types from MODIS Collection 5.1 land cover dataset are used, crosswalking land cover types to PFT fractions. Age distributions are based on country-level forest inventories for temperate and high latitude regions, and based on biomass for tropical regions. | Biodiversity-Ecosystem Function |
| Proportion of trees that older than 140 years | (*32*) | Unitless (proportion) | https://doi.pangaea.de/10.1594/PANGAEA.889943 | Four Plant Functional Types from MODIS Collection 5.1 land cover dataset are used, crosswalking land cover types to PFT fractions. Age distributions are based on country-level forest inventories for temperate and high latitude regions, and based on biomass for tropical regions. | Biodiversity-Ecosystem Function |
| Aboveground biomass (for the year 2010) | (*33*) | Mg/ha | https://globbiomass.org/wp-content/uploads/GB_Maps/Globbiomass_global_dataset.html | (*33*) derived Aboveground Biomass values from their Growing Stock Volume estimates with a set of Biomass Expansion and Conversion Factors (BCEF) following approaches to extend on ground estimates of wood density and stem-to-total biomass expansion factors to obtain a global raster dataset. | Biodiversity-Ecosystem Function |
| Volume of all living trees > 10cm diameter at breast height | (*33*) | m3/ha | https://globbiomass.org/wp-content/uploads/GB_Maps/Globbiomass_global_dataset.html | (*33*) derived Growing Stock Volume estimates from spaceborne SAR (ALOS PALSAR, Envisat ASAR), optical (Landsat-7), LiDAR (ICESAT) and auxiliary datasets with multiple estimation procedures. | Biodiversity-Ecosystem Function |
| Tree cover in the year 2000 | (*34*) | % | https://data.globalforestwatch | (*34*) derived global tree cover from Landsat data | Biodiversity-Ecosystem Function |

| | | | | | |
|---|---|---|---|---|---|
| | | | .org/datasets/14228e6347c44f5691572169e9e107ad | | |
| Tree cover in the year 2010 | (*34*) | % | https://glad.umd.edu/Potapov/TCC_2010/ | (*34*) derived global tree cover from Landsat data | Biodiversity-Ecosystem Function |
| Enhanced Vegetation Index (EVI) | https://lpdaac.usgs.gov/products/myd13q1v006/ | | https://explorer.earthengine.google.com/#detail/MODIS%2F006%2FMYD13Q1 | MYD13Q1.006 Vegetation Indices 16-Day Global 250m: Enhanced Vegetation Index (EVI). Scaled units (0.0001); cloud composite; No data from Antarctica; default: averaged from the calendar years 2015-2019 | Biodiversity-Ecosystem Function |
| Gross Primary Production (GPP) | (*35*) | 0.0001 * kg*C/m^2 | https://explorer.earthengine.google.com/#detail/MODIS%2F055%2FMOD17A3 | MOD17A3.055: Terra Gross Primary Production Yearly Global 1km. Yearly data; default: averaged for the years 2010-2014 | Biodiversity-Ecosystem Function |
| Leaf Area Index (LAI) | https://lpdaac.usgs.gov/products/mcd15a3hv006/ | m^2 / m^2 | https://explorer.earthengine.google.com/#detail/MODIS%2F006%2FMCD15A3H | MCD15A3H.006 Terra+Aqua Leaf Area Index. Masks substantial quantities of arid areas; default: averaged from the calendar years 2015-2019 | Biodiversity-Ecosystem Function |
| Normalized Difference Vegetation Index (NDVI) | https://lpdaac.usgs.gov/products/myd13q1v006/ | unitless | https://explorer.earthengine.google.com/#detail/MODIS%2F006%2FMYD13Q1 | MYD13Q1.006 Vegetation Indices 16-Day Global 250m: NDVI. Scaled units (0.0001); cloud composite; No data from Antarctica; default: averaged from the calendar years 2015-2019 | Biodiversity-Ecosystem Function |

| | | | | | |
|---|---|---|---|---|---|
| Net Primary Production (NPP) | (*35*) | 0.0001 * kg*C/m^2 | https://explorer.earthengine.google.com/#detail/MODIS%2F055%2FMOD17A3 | MOD17A3.055: Terra Net Primary Production Yearly Global 1km. Scaled units (0.0001); Yearly data; default: averaged for the years 2010-2014 | Biodiversity-Ecosystem Function |
| Forest canopy height | (*36*) | m | https://glad.umd.edu/dataset/gedi | (*36*) integrated Landsat data with GEDI Lidar forest structure measurements | Biodiversity-Ecosystem Function |
| Absolute Depth to Bedrock | (*23*) | cm | https://www.soilgrids.org ; https://www.isric.org/explore/wosis/accessing-wosis-derived-datasets | (*23*) used data on 150,000 soil profiles worldwide to create a global map, integrating observations within profile to obtain datapoints at fixed depth.A machine learning approach was used, with input data on: Digital Elevation Model-derived characteristics (using the merged DEMS SRTMGL3 and GMTED, as described in (*24*)), long-term average and sd of MODIS-based EVI, bands 4 and 7, long-term monthly average and standard deviation in land surface temperature (MODIS LST), long-term averaged mean monthly hours of snow cover (MODIS 8-day snow occurrence), land cover (using GlobCover30, described in (*25*), who classified Landsat data from 2000 and 2010), precipitation data (averaging WorldClim and GPCP), lithology (using GLiM, (*26*)), Global water table depth (*27*), long-term averaged mean monthly MODIS Flood Water, Landsat-based distribution of mangroves (*28*) and average soil and sedimentary-deposit thickness (*29*) | Biodiversity-Ecosystem Function |
| Bulk density (fine earth) 0.00m | (*23*) | kg / cubic–meter | https://www.soilgrids.org ; https://www.isric.org/explore/wosis/accessing-wosis-derived-datasets | (*23*) used data on 150,000 soil profiles worldwide to create a global map, integrating observations within profile to obtain datapoints at fixed depth.A machine learning approach was used, with input data on: Digital Elevation Model-derived characteristics (using the merged DEMS SRTMGL3 and GMTED, as described in (*24*)), long-term average and sd of MODIS-based EVI, bands 4 and 7, long-term monthly average and standard deviation in land surface temperature (MODIS LST), long-term averaged mean monthly hours of snow cover (MODIS 8-day snow occurrence), land cover (using GlobCover30, described in (*25*), who classified Landsat data from 2000 and 2010), precipitation data (averaging WorldClim and GPCP), lithology (using GLiM, (*26*)), Global water table depth (*27*), long-term averaged mean monthly MODIS Flood Water, Landsat-based | Biodiversity-Ecosystem Function |

| | | | | | |
|---|---|---|---|---|---|
| | | | | distribution of mangroves (*28*) and average soil and sedimentary-deposit thickness (*29*) | |
| Bulk density (fine earth) at 0.05m | (*23*) | kg / cubic–meter | https://www.soilgrids.org ; https://www.isric.org/explore/wosis/accessing-wosis-derived-datasets | (*23*) used data on 150,000 soil profiles worldwide to create a global map, integrating observations within profile to obtain datapoints at fixed depth.A machine learning approach was used, with input data on: Digital Elevation Model-derived characteristics (using the merged DEMS SRTMGL3 and GMTED, as described in (*24*)), long-term average and sd of MODIS-based EVI, bands 4 and 7, long-term monthly average and standard deviation in land surface temperature (MODIS LST), long-term averaged mean monthly hours of snow cover (MODIS 8-day snow occurrence), land cover (using GlobCover30, described in (*25*), who classified Landsat data from 2000 and 2010), precipitation data (averaging WorldClim and GPCP), lithology (using GLiM, (*26*)), Global water table depth (*27*), long-term averaged mean monthly MODIS Flood Water, Landsat-based distribution of mangroves (*28*) and average soil and sedimentary-deposit thickness (*29*) | Biodiversity-Ecosystem Function |
| Bulk density (fine earth) at 0.15m | (*23*) | kg / cubic–meter | https://www.soilgrids.org ; https://www.isric.org/explore/wosis/accessing-wosis-derived-datasets | (*23*) used data on 150,000 soil profiles worldwide to create a global map, integrating observations within profile to obtain datapoints at fixed depth.A machine learning approach was used, with input data on: Digital Elevation Model-derived characteristics (using the merged DEMS SRTMGL3 and GMTED, as described in (*24*)), long-term average and sd of MODIS-based EVI, bands 4 and 7, long-term monthly average and standard deviation in land surface temperature (MODIS LST), long-term averaged mean monthly hours of snow cover (MODIS 8-day snow occurrence), land cover (using GlobCover30, described in (*25*), who classified Landsat data from 2000 and 2010), precipitation data (averaging WorldClim and GPCP), lithology (using GLiM, (*26*)), Global water table depth (*27*), long-term averaged mean monthly MODIS Flood Water, Landsat-based distribution of mangroves (*28*) and average soil and sedimentary-deposit thickness (*29*) | Biodiversity-Ecosystem Function |
| Bulk density (fine earth) at 0.30m | (*23*) | kg / cubic–meter | https://www.soilgrids.org ; https://www.isric.org/explore/wosis/accessin | (*23*) used data on 150,000 soil profiles worldwide to create a global map, integrating observations within profile to obtain datapoints at fixed depth.A machine learning approach was used, with input data on: Digital Elevation Model-derived characteristics (using the merged DEMS SRTMGL3 and GMTED, as described in (*24*)), long-term average and sd of MODIS- | Biodiversity-Ecosystem Function |

| | | | | | |
|---|---|---|---|---|---|
| | | | g-wosis-derived-datasets | based EVI, bands 4 and 7, long-term monthly average and standard deviation in land surface temperature (MODIS LST), long-term averaged mean monthly hours of snow cover (MODIS 8-day snow occurrence), land cover (using GlobCover30, described in (*25*), who classified Landsat data from 2000 and 2010), precipitation data (averaging WorldClim and GPCP), lithology (using GLiM, (*26*)), Global water table depth (*27*), long-term averaged mean monthly MODIS Flood Water, Landsat-based distribution of mangroves (*28*) and average soil and sedimentary-deposit thickness (*29*) | |
| Bulk density (fine earth) at 0.60m | (*23*) | kg / cubic–meter | https://www.soilgrids.org ; https://www.isric.org/explore/wosis/accessing-wosis-derived-datasets | (*23*) used data on 150,000 soil profiles worldwide to create a global map, integrating observations within profile to obtain datapoints at fixed depth.A machine learning approach was used, with input data on: Digital Elevation Model-derived characteristics (using the merged DEMS SRTMGL3 and GMTED, as described in (*24*)), long-term average and sd of MODIS-based EVI, bands 4 and 7, long-term monthly average and standard deviation in land surface temperature (MODIS LST), long-term averaged mean monthly hours of snow cover (MODIS 8-day snow occurrence), land cover (using GlobCover30, described in (*25*), who classified Landsat data from 2000 and 2010), precipitation data (averaging WorldClim and GPCP), lithology (using GLiM, (*26*)), Global water table depth (*27*), long-term averaged mean monthly MODIS Flood Water, Landsat-based distribution of mangroves (*28*) and average soil and sedimentary-deposit thickness (*29*) | Biodiversity-Ecosystem Function |
| Bulk density (fine earth) at 1.00m | (*23*) | kg / cubic–meter | https://www.soilgrids.org ; https://www.isric.org/explore/wosis/accessing-wosis-derived-datasets | (*23*) used data on 150,000 soil profiles worldwide to create a global map, integrating observations within profile to obtain datapoints at fixed depth.A machine learning approach was used, with input data on: Digital Elevation Model-derived characteristics (using the merged DEMS SRTMGL3 and GMTED, as described in (*24*)), long-term average and sd of MODIS-based EVI, bands 4 and 7, long-term monthly average and standard deviation in land surface temperature (MODIS LST), long-term averaged mean monthly hours of snow cover (MODIS 8-day snow occurrence), land cover (using GlobCover30, described in (*25*), who classified Landsat data from 2000 and 2010), precipitation data (averaging WorldClim and GPCP), lithology (using GLiM, (*26*)), Global water table depth (*27*), long-term averaged mean monthly MODIS Flood Water, Landsat-based | Biodiversity-Ecosystem Function |

| | | | | | |
|---|---|---|---|---|---|
| | | | | distribution of mangroves (*28*) and average soil and sedimentary-deposit thickness (*29*) | |
| Bulk density (fine earth) at 2.00m | (*23*) | kg / cubic–meter | https://www.soilgrids.org ; https://www.isric.org/explore/wosis/accessing-wosis-derived-datasets | (*23*) used data on 150,000 soil profiles worldwide to create a global map, integrating observations within profile to obtain datapoints at fixed depth.A machine learning approach was used, with input data on: Digital Elevation Model-derived characteristics (using the merged DEMS SRTMGL3 and GMTED, as described in (*24*)), long-term average and sd of MODIS-based EVI, bands 4 and 7, long-term monthly average and standard deviation in land surface temperature (MODIS LST), long-term averaged mean monthly hours of snow cover (MODIS 8-day snow occurrence), land cover (using GlobCover30, described in (*25*), who classified Landsat data from 2000 and 2010), precipitation data (averaging WorldClim and GPCP), lithology (using GLiM, (*26*)), Global water table depth (*27*), long-term averaged mean monthly MODIS Flood Water, Landsat-based distribution of mangroves (*28*) and average soil and sedimentary-deposit thickness (*29*) | Biodiversity-Ecosystem Function |
| Cation exchange capacity of soil at 0.00m | (*23*) | cmolc/kg | https://www.soilgrids.org ; https://www.isric.org/explore/wosis/accessing-wosis-derived-datasets | (*23*) used data on 150,000 soil profiles worldwide to create a global map, integrating observations within profile to obtain datapoints at fixed depth.A machine learning approach was used, with input data on: Digital Elevation Model-derived characteristics (using the merged DEMS SRTMGL3 and GMTED, as described in (*24*)), long-term average and sd of MODIS-based EVI, bands 4 and 7, long-term monthly average and standard deviation in land surface temperature (MODIS LST), long-term averaged mean monthly hours of snow cover (MODIS 8-day snow occurrence), land cover (using GlobCover30, described in (*25*), who classified Landsat data from 2000 and 2010), precipitation data (averaging WorldClim and GPCP), lithology (using GLiM, (*26*)), Global water table depth (*27*), long-term averaged mean monthly MODIS Flood Water, Landsat-based distribution of mangroves (*28*) and average soil and sedimentary-deposit thickness (*29*) | Biodiversity-Ecosystem Function |
| Cation exchange capacity of soil at 0.05m | (*23*) | cmolc/kg | https://www.soilgrids.org ; https://www.isric.org/explore/wosis/accessin | (*23*) used data on 150,000 soil profiles worldwide to create a global map, integrating observations within profile to obtain datapoints at fixed depth.A machine learning approach was used, with input data on: Digital Elevation Model-derived characteristics (using the merged DEMS SRTMGL3 and GMTED, as described in (*24*)), long-term average and sd of MODIS- | Biodiversity-Ecosystem Function |

| | | | | | |
|---|---|---|---|---|---|
| | | | g-wosis-derived-datasets | based EVI, bands 4 and 7, long-term monthly average and standard deviation in land surface temperature (MODIS LST), long-term averaged mean monthly hours of snow cover (MODIS 8-day snow occurrence), land cover (using GlobCover30, described in (*25*), who classified Landsat data from 2000 and 2010), precipitation data (averaging WorldClim and GPCP), lithology (using GLiM, (*26*)), Global water table depth (*27*), long-term averaged mean monthly MODIS Flood Water, Landsat-based distribution of mangroves (*28*) and average soil and sedimentary-deposit thickness (*29*) | |
| Cation exchange capacity of soil at 0.15m | (*23*) | cmolc/kg | https://www.soilgrids.org ; https://www.isric.org/explore/wosis/accessing-wosis-derived-datasets | (*23*) used data on 150,000 soil profiles worldwide to create a global map, integrating observations within profile to obtain datapoints at fixed depth.A machine learning approach was used, with input data on: Digital Elevation Model-derived characteristics (using the merged DEMS SRTMGL3 and GMTED, as described in (*24*)), long-term average and sd of MODIS-based EVI, bands 4 and 7, long-term monthly average and standard deviation in land surface temperature (MODIS LST), long-term averaged mean monthly hours of snow cover (MODIS 8-day snow occurrence), land cover (using GlobCover30, described in (*25*), who classified Landsat data from 2000 and 2010), precipitation data (averaging WorldClim and GPCP), lithology (using GLiM, (*26*)), Global water table depth (*27*), long-term averaged mean monthly MODIS Flood Water, Landsat-based distribution of mangroves (*28*) and average soil and sedimentary-deposit thickness (*29*) | Biodiversity-Ecosystem Function |
| Cation exchange capacity of soil at 0.30m | (*23*) | cmolc/kg | https://www.soilgrids.org ; https://www.isric.org/explore/wosis/accessing-wosis-derived-datasets | (*23*) used data on 150,000 soil profiles worldwide to create a global map, integrating observations within profile to obtain datapoints at fixed depth.A machine learning approach was used, with input data on: Digital Elevation Model-derived characteristics (using the merged DEMS SRTMGL3 and GMTED, as described in (*24*)), long-term average and sd of MODIS-based EVI, bands 4 and 7, long-term monthly average and standard deviation in land surface temperature (MODIS LST), long-term averaged mean monthly hours of snow cover (MODIS 8-day snow occurrence), land cover (using GlobCover30, described in (*25*), who classified Landsat data from 2000 and 2010), precipitation data (averaging WorldClim and GPCP), lithology (using GLiM, (*26*)), Global water table depth (*27*), long-term averaged mean monthly MODIS Flood Water, Landsat-based | Biodiversity-Ecosystem Function |

| | | | | | |
|---|---|---|---|---|---|
| | | | | distribution of mangroves (*28*) and average soil and sedimentary-deposit thickness (*29*) | |
| Cation exchange capacity of soil at 0.60m | (*23*) | cmolc/kg | https://www.soilgrids.org ; https://www.isric.org/explore/wosis/accessing-wosis-derived-datasets | (*23*) used data on 150,000 soil profiles worldwide to create a global map, integrating observations within profile to obtain datapoints at fixed depth.A machine learning approach was used, with input data on: Digital Elevation Model-derived characteristics (using the merged DEMS SRTMGL3 and GMTED, as described in (*24*)), long-term average and sd of MODIS-based EVI, bands 4 and 7, long-term monthly average and standard deviation in land surface temperature (MODIS LST), long-term averaged mean monthly hours of snow cover (MODIS 8-day snow occurrence), land cover (using GlobCover30, described in (*25*), who classified Landsat data from 2000 and 2010), precipitation data (averaging WorldClim and GPCP), lithology (using GLiM, (*26*)), Global water table depth (*27*), long-term averaged mean monthly MODIS Flood Water, Landsat-based distribution of mangroves (*28*) and average soil and sedimentary-deposit thickness (*29*) | Biodiversity-Ecosystem Function |
| Cation exchange capacity of soil at 1.00m | (*23*) | cmolc/kg | https://www.soilgrids.org ; https://www.isric.org/explore/wosis/accessing-wosis-derived-datasets | (*23*) used data on 150,000 soil profiles worldwide to create a global map, integrating observations within profile to obtain datapoints at fixed depth.A machine learning approach was used, with input data on: Digital Elevation Model-derived characteristics (using the merged DEMS SRTMGL3 and GMTED, as described in (*24*)), long-term average and sd of MODIS-based EVI, bands 4 and 7, long-term monthly average and standard deviation in land surface temperature (MODIS LST), long-term averaged mean monthly hours of snow cover (MODIS 8-day snow occurrence), land cover (using GlobCover30, described in (*25*), who classified Landsat data from 2000 and 2010), precipitation data (averaging WorldClim and GPCP), lithology (using GLiM, (*26*)), Global water table depth (*27*), long-term averaged mean monthly MODIS Flood Water, Landsat-based distribution of mangroves (*28*) and average soil and sedimentary-deposit thickness (*29*) | Biodiversity-Ecosystem Function |
| Cation exchange capacity of soil at 2.00m | (*23*) | cmolc/kg | https://www.soilgrids.org ; https://www.isric.org/explore/wosis/accessin | (*23*) used data on 150,000 soil profiles worldwide to create a global map, integrating observations within profile to obtain datapoints at fixed depth.A machine learning approach was used, with input data on: Digital Elevation Model-derived characteristics (using the merged DEMS SRTMGL3 and GMTED, as described in (*24*)), long-term average and sd of MODIS- | Biodiversity-Ecosystem Function |

| | | | | | |
|---|---|---|---|---|---|
| | | | g-wosis-derived-datasets | based EVI, bands 4 and 7, long-term monthly average and standard deviation in land surface temperature (MODIS LST), long-term averaged mean monthly hours of snow cover (MODIS 8-day snow occurrence), land cover (using GlobCover30, described in (*25*), who classified Landsat data from 2000 and 2010), precipitation data (averaging WorldClim and GPCP), lithology (using GLiM, (*26*)), Global water table depth (*27*), long-term averaged mean monthly MODIS Flood Water, Landsat-based distribution of mangroves (*28*) and average soil and sedimentary-deposit thickness (*29*) | |
| Clay content (0-2 micro meter) at 0.00m | (*23*) | mass fraction in % | https://www.soilgrids.org ; https://www.isric.org/explore/wosis/accessing-wosis-derived-datasets | (*23*) used data on 150,000 soil profiles worldwide to create a global map, integrating observations within profile to obtain datapoints at fixed depth.A machine learning approach was used, with input data on: Digital Elevation Model-derived characteristics (using the merged DEMS SRTMGL3 and GMTED, as described in (*24*)), long-term average and sd of MODIS-based EVI, bands 4 and 7, long-term monthly average and standard deviation in land surface temperature (MODIS LST), long-term averaged mean monthly hours of snow cover (MODIS 8-day snow occurrence), land cover (using GlobCover30, described in (*25*), who classified Landsat data from 2000 and 2010), precipitation data (averaging WorldClim and GPCP), lithology (using GLiM, (*26*)), Global water table depth (*27*), long-term averaged mean monthly MODIS Flood Water, Landsat-based distribution of mangroves (*28*) and average soil and sedimentary-deposit thickness (*29*) | Biodiversity-Ecosystem Function |
| Clay content (0-2 micro meter) at 0.05m | (*23*) | mass fraction in % | https://www.soilgrids.org ; https://www.isric.org/explore/wosis/accessing-wosis-derived-datasets | (*23*) used data on 150,000 soil profiles worldwide to create a global map, integrating observations within profile to obtain datapoints at fixed depth.A machine learning approach was used, with input data on: Digital Elevation Model-derived characteristics (using the merged DEMS SRTMGL3 and GMTED, as described in (*24*)), long-term average and sd of MODIS-based EVI, bands 4 and 7, long-term monthly average and standard deviation in land surface temperature (MODIS LST), long-term averaged mean monthly hours of snow cover (MODIS 8-day snow occurrence), land cover (using GlobCover30, described in (*25*), who classified Landsat data from 2000 and 2010), precipitation data (averaging WorldClim and GPCP), lithology (using GLiM, (*26*)), Global water table depth (*27*), long-term averaged mean monthly MODIS Flood Water, Landsat-based | Biodiversity-Ecosystem Function |

| | | | | | |
|---|---|---|---|---|---|
| | | | | distribution of mangroves (*28*) and average soil and sedimentary-deposit thickness (*29*) | |
| Clay content (0-2 micro meter) at 0.15m | (*23*) | mass fraction in % | https://www.soilgrids.org ; https://www.isric.org/explore/wosis/accessing-wosis-derived-datasets | (*23*) used data on 150,000 soil profiles worldwide to create a global map, integrating observations within profile to obtain datapoints at fixed depth.A machine learning approach was used, with input data on: Digital Elevation Model-derived characteristics (using the merged DEMS SRTMGL3 and GMTED, as described in (*24*)), long-term average and sd of MODIS-based EVI, bands 4 and 7, long-term monthly average and standard deviation in land surface temperature (MODIS LST), long-term averaged mean monthly hours of snow cover (MODIS 8-day snow occurrence), land cover (using GlobCover30, described in (*25*), who classified Landsat data from 2000 and 2010), precipitation data (averaging WorldClim and GPCP), lithology (using GLiM, (*26*)), Global water table depth (*27*), long-term averaged mean monthly MODIS Flood Water, Landsat-based distribution of mangroves (*28*) and average soil and sedimentary-deposit thickness (*29*) | Biodiversity-Ecosystem Function |
| Clay content (0-2 micro meter) at 0.30m | (*23*) | mass fraction in % | https://www.soilgrids.org ; https://www.isric.org/explore/wosis/accessing-wosis-derived-datasets | (*23*) used data on 150,000 soil profiles worldwide to create a global map, integrating observations within profile to obtain datapoints at fixed depth.A machine learning approach was used, with input data on: Digital Elevation Model-derived characteristics (using the merged DEMS SRTMGL3 and GMTED, as described in (*24*)), long-term average and sd of MODIS-based EVI, bands 4 and 7, long-term monthly average and standard deviation in land surface temperature (MODIS LST), long-term averaged mean monthly hours of snow cover (MODIS 8-day snow occurrence), land cover (using GlobCover30, described in (*25*), who classified Landsat data from 2000 and 2010), precipitation data (averaging WorldClim and GPCP), lithology (using GLiM, (*26*)), Global water table depth (*27*), long-term averaged mean monthly MODIS Flood Water, Landsat-based distribution of mangroves (*28*) and average soil and sedimentary-deposit thickness (*29*) | Biodiversity-Ecosystem Function |
| Clay content (0-2 micro meter) at 0.60m | (*23*) | mass fraction in % | https://www.soilgrids.org ; https://www.isric.org/explore/wosis/accessin | (*23*) used data on 150,000 soil profiles worldwide to create a global map, integrating observations within profile to obtain datapoints at fixed depth.A machine learning approach was used, with input data on: Digital Elevation Model-derived characteristics (using the merged DEMS SRTMGL3 and GMTED, as described in (*24*)), long-term average and sd of MODIS- | Biodiversity-Ecosystem Function |

| | | | | | |
|---|---|---|---|---|---|
| | | | g-wosis-derived-datasets | based EVI, bands 4 and 7, long-term monthly average and standard deviation in land surface temperature (MODIS LST), long-term averaged mean monthly hours of snow cover (MODIS 8-day snow occurrence), land cover (using GlobCover30, described in (*25*), who classified Landsat data from 2000 and 2010), precipitation data (averaging WorldClim and GPCP), lithology (using GLiM, (*26*)), Global water table depth (*27*), long-term averaged mean monthly MODIS Flood Water, Landsat-based distribution of mangroves (*28*) and average soil and sedimentary-deposit thickness (*29*) | |
| Clay content (0-2 micro meter) at 1.00m | (*23*) | mass fraction in % | https://www.soilgrids.org ; https://www.isric.org/explore/wosis/accessing-wosis-derived-datasets | (*23*) used data on 150,000 soil profiles worldwide to create a global map, integrating observations within profile to obtain datapoints at fixed depth.A machine learning approach was used, with input data on: Digital Elevation Model-derived characteristics (using the merged DEMS SRTMGL3 and GMTED, as described in (*24*)), long-term average and sd of MODIS-based EVI, bands 4 and 7, long-term monthly average and standard deviation in land surface temperature (MODIS LST), long-term averaged mean monthly hours of snow cover (MODIS 8-day snow occurrence), land cover (using GlobCover30, described in (*25*), who classified Landsat data from 2000 and 2010), precipitation data (averaging WorldClim and GPCP), lithology (using GLiM, (*26*)), Global water table depth (*27*), long-term averaged mean monthly MODIS Flood Water, Landsat-based distribution of mangroves (*28*) and average soil and sedimentary-deposit thickness (*29*) | Biodiversity-Ecosystem Function |
| Clay content (0-2 micro meter) at 2.00m | (*23*) | mass fraction in % | https://www.soilgrids.org ; https://www.isric.org/explore/wosis/accessing-wosis-derived-datasets | (*23*) used data on 150,000 soil profiles worldwide to create a global map, integrating observations within profile to obtain datapoints at fixed depth.A machine learning approach was used, with input data on: Digital Elevation Model-derived characteristics (using the merged DEMS SRTMGL3 and GMTED, as described in (*24*)), long-term average and sd of MODIS-based EVI, bands 4 and 7, long-term monthly average and standard deviation in land surface temperature (MODIS LST), long-term averaged mean monthly hours of snow cover (MODIS 8-day snow occurrence), land cover (using GlobCover30, described in (*25*), who classified Landsat data from 2000 and 2010), precipitation data (averaging WorldClim and GPCP), lithology (using GLiM, (*26*)), Global water table depth (*27*), long-term averaged mean monthly MODIS Flood Water, Landsat-based | Biodiversity-Ecosystem Function |

| | | | | | |
|---|---|---|---|---|---|
| | | | | distribution of mangroves (*28*) and average soil and sedimentary-deposit thickness (*29*) | |
| Coarse fragments volumetric at 0.00m | (*23*) | % | https://www.soilgrids.org ; https://www.isric.org/explore/wosis/accessing-wosis-derived-datasets | (*23*) used data on 150,000 soil profiles worldwide to create a global map, integrating observations within profile to obtain datapoints at fixed depth.A machine learning approach was used, with input data on: Digital Elevation Model-derived characteristics (using the merged DEMS SRTMGL3 and GMTED, as described in (*24*)), long-term average and sd of MODIS-based EVI, bands 4 and 7, long-term monthly average and standard deviation in land surface temperature (MODIS LST), long-term averaged mean monthly hours of snow cover (MODIS 8-day snow occurrence), land cover (using GlobCover30, described in (*25*), who classified Landsat data from 2000 and 2010), precipitation data (averaging WorldClim and GPCP), lithology (using GLiM, (*26*)), Global water table depth (*27*), long-term averaged mean monthly MODIS Flood Water, Landsat-based distribution of mangroves (*28*) and average soil and sedimentary-deposit thickness (*29*) | Biodiversity-Ecosystem Function |
| Coarse fragments volumetric at 0.05m | (*23*) | % | https://www.soilgrids.org ; https://www.isric.org/explore/wosis/accessing-wosis-derived-datasets | (*23*) used data on 150,000 soil profiles worldwide to create a global map, integrating observations within profile to obtain datapoints at fixed depth.A machine learning approach was used, with input data on: Digital Elevation Model-derived characteristics (using the merged DEMS SRTMGL3 and GMTED, as described in (*24*)), long-term average and sd of MODIS-based EVI, bands 4 and 7, long-term monthly average and standard deviation in land surface temperature (MODIS LST), long-term averaged mean monthly hours of snow cover (MODIS 8-day snow occurrence), land cover (using GlobCover30, described in (*25*), who classified Landsat data from 2000 and 2010), precipitation data (averaging WorldClim and GPCP), lithology (using GLiM, (*26*)), Global water table depth (*27*), long-term averaged mean monthly MODIS Flood Water, Landsat-based distribution of mangroves (*28*) and average soil and sedimentary-deposit thickness (*29*) | Biodiversity-Ecosystem Function |
| Coarse fragments volumetric at 0.15m | (*23*) | % | https://www.soilgrids.org ; https://www.isric.org/explore/wosis/accessin | (*23*) used data on 150,000 soil profiles worldwide to create a global map, integrating observations within profile to obtain datapoints at fixed depth.A machine learning approach was used, with input data on: Digital Elevation Model-derived characteristics (using the merged DEMS SRTMGL3 and GMTED, as described in (*24*)), long-term average and sd of MODIS- | Biodiversity-Ecosystem Function |

| | | | | | |
|---|---|---|---|---|---|
| | | | g-wosis-derived-datasets | based EVI, bands 4 and 7, long-term monthly average and standard deviation in land surface temperature (MODIS LST), long-term averaged mean monthly hours of snow cover (MODIS 8-day snow occurrence), land cover (using GlobCover30, described in (*25*), who classified Landsat data from 2000 and 2010), precipitation data (averaging WorldClim and GPCP), lithology (using GLiM, (*26*)), Global water table depth (*27*), long-term averaged mean monthly MODIS Flood Water, Landsat-based distribution of mangroves (*28*) and average soil and sedimentary-deposit thickness (*29*) | |
| Coarse fragments volumetric at 0.30m | (*23*) | % | https://www.soilgrids.org ; https://www.isric.org/explore/wosis/accessing-wosis-derived-datasets | (*23*) used data on 150,000 soil profiles worldwide to create a global map, integrating observations within profile to obtain datapoints at fixed depth.A machine learning approach was used, with input data on: Digital Elevation Model-derived characteristics (using the merged DEMS SRTMGL3 and GMTED, as described in (*24*)), long-term average and sd of MODIS-based EVI, bands 4 and 7, long-term monthly average and standard deviation in land surface temperature (MODIS LST), long-term averaged mean monthly hours of snow cover (MODIS 8-day snow occurrence), land cover (using GlobCover30, described in (*25*), who classified Landsat data from 2000 and 2010), precipitation data (averaging WorldClim and GPCP), lithology (using GLiM, (*26*)), Global water table depth (*27*), long-term averaged mean monthly MODIS Flood Water, Landsat-based distribution of mangroves (*28*) and average soil and sedimentary-deposit thickness (*29*) | Biodiversity-Ecosystem Function |
| Coarse fragments volumetric at 0.60m | (*23*) | % | https://www.soilgrids.org ; https://www.isric.org/explore/wosis/accessing-wosis-derived-datasets | (*23*) used data on 150,000 soil profiles worldwide to create a global map, integrating observations within profile to obtain datapoints at fixed depth.A machine learning approach was used, with input data on: Digital Elevation Model-derived characteristics (using the merged DEMS SRTMGL3 and GMTED, as described in (*24*)), long-term average and sd of MODIS-based EVI, bands 4 and 7, long-term monthly average and standard deviation in land surface temperature (MODIS LST), long-term averaged mean monthly hours of snow cover (MODIS 8-day snow occurrence), land cover (using GlobCover30, described in (*25*), who classified Landsat data from 2000 and 2010), precipitation data (averaging WorldClim and GPCP), lithology (using GLiM, (*26*)), Global water table depth (*27*), long-term averaged mean monthly MODIS Flood Water, Landsat-based | Biodiversity-Ecosystem Function |

| | | | | | |
|---|---|---|---|---|---|
| | | | | distribution of mangroves (*28*) and average soil and sedimentary-deposit thickness (*29*) | |
| Coarse fragments volumetric at 1.00m | (*23*) | % | https://www.soilgrids.org ; https://www.isric.org/explore/wosis/accessing-wosis-derived-datasets | (*23*) used data on 150,000 soil profiles worldwide to create a global map, integrating observations within profile to obtain datapoints at fixed depth.A machine learning approach was used, with input data on: Digital Elevation Model-derived characteristics (using the merged DEMS SRTMGL3 and GMTED, as described in (*24*)), long-term average and sd of MODIS-based EVI, bands 4 and 7, long-term monthly average and standard deviation in land surface temperature (MODIS LST), long-term averaged mean monthly hours of snow cover (MODIS 8-day snow occurrence), land cover (using GlobCover30, described in (*25*), who classified Landsat data from 2000 and 2010), precipitation data (averaging WorldClim and GPCP), lithology (using GLiM, (*26*)), Global water table depth (*27*), long-term averaged mean monthly MODIS Flood Water, Landsat-based distribution of mangroves (*28*) and average soil and sedimentary-deposit thickness (*29*) | Biodiversity-Ecosystem Function |
| Coarse fragments volumetric at 2.00m | (*23*) | % | https://www.soilgrids.org ; https://www.isric.org/explore/wosis/accessing-wosis-derived-datasets | (*23*) used data on 150,000 soil profiles worldwide to create a global map, integrating observations within profile to obtain datapoints at fixed depth.A machine learning approach was used, with input data on: Digital Elevation Model-derived characteristics (using the merged DEMS SRTMGL3 and GMTED, as described in (*24*)), long-term average and sd of MODIS-based EVI, bands 4 and 7, long-term monthly average and standard deviation in land surface temperature (MODIS LST), long-term averaged mean monthly hours of snow cover (MODIS 8-day snow occurrence), land cover (using GlobCover30, described in (*25*), who classified Landsat data from 2000 and 2010), precipitation data (averaging WorldClim and GPCP), lithology (using GLiM, (*26*)), Global water table depth (*27*), long-term averaged mean monthly MODIS Flood Water, Landsat-based distribution of mangroves (*28*) and average soil and sedimentary-deposit thickness (*29*) | Biodiversity-Ecosystem Function |
| Depth to Bedrock (up to 200cm) | (*23*) | cm (up to 200) | https://www.soilgrids.org ; https://www.isric.org/explore/wosis/accessin | (*23*) used data on 150,000 soil profiles worldwide to create a global map, integrating observations within profile to obtain datapoints at fixed depth.A machine learning approach was used, with input data on: Digital Elevation Model-derived characteristics (using the merged DEMS SRTMGL3 and GMTED, as described in (*24*)), long-term average and sd of MODIS- | Biodiversity-Ecosystem Function |

| | | | | | |
|---|---|---|---|---|---|
| | | | g-wosis-derived-datasets | based EVI, bands 4 and 7, long-term monthly average and standard deviation in land surface temperature (MODIS LST), long-term averaged mean monthly hours of snow cover (MODIS 8-day snow occurrence), land cover (using GlobCover30, described in (*25*), who classified Landsat data from 2000 and 2010), precipitation data (averaging WorldClim and GPCP), lithology (using GLiM, (*26*)), Global water table depth (*27*), long-term averaged mean monthly MODIS Flood Water, Landsat-based distribution of mangroves (*28*) and average soil and sedimentary-deposit thickness (*29*) | |
| Soil organic carbon content (fine earth fraction) at 0.00m | (*23*) | g per kg | https://www.soilgrids.org ; https://www.isric.org/explore/wosis/accessing-wosis-derived-datasets | (*23*) used data on 150,000 soil profiles worldwide to create a global map, integrating observations within profile to obtain datapoints at fixed depth.A machine learning approach was used, with input data on: Digital Elevation Model-derived characteristics (using the merged DEMS SRTMGL3 and GMTED, as described in (*24*)), long-term average and sd of MODIS-based EVI, bands 4 and 7, long-term monthly average and standard deviation in land surface temperature (MODIS LST), long-term averaged mean monthly hours of snow cover (MODIS 8-day snow occurrence), land cover (using GlobCover30, described in (*25*), who classified Landsat data from 2000 and 2010), precipitation data (averaging WorldClim and GPCP), lithology (using GLiM, (*26*)), Global water table depth (*27*), long-term averaged mean monthly MODIS Flood Water, Landsat-based distribution of mangroves (*28*) and average soil and sedimentary-deposit thickness (*29*) | Biodiversity-Ecosystem Function |
| Soil organic carbon content (fine earth fraction) at 0.05m | (*23*) | g per kg | https://www.soilgrids.org ; https://www.isric.org/explore/wosis/accessing-wosis-derived-datasets | (*23*) used data on 150,000 soil profiles worldwide to create a global map, integrating observations within profile to obtain datapoints at fixed depth.A machine learning approach was used, with input data on: Digital Elevation Model-derived characteristics (using the merged DEMS SRTMGL3 and GMTED, as described in (*24*)), long-term average and sd of MODIS-based EVI, bands 4 and 7, long-term monthly average and standard deviation in land surface temperature (MODIS LST), long-term averaged mean monthly hours of snow cover (MODIS 8-day snow occurrence), land cover (using GlobCover30, described in (*25*), who classified Landsat data from 2000 and 2010), precipitation data (averaging WorldClim and GPCP), lithology (using GLiM, (*26*)), Global water table depth (*27*), long-term averaged mean monthly MODIS Flood Water, Landsat-based | Biodiversity-Ecosystem Function |

| | | | | | |
|---|---|---|---|---|---|
| | | | | distribution of mangroves (*28*) and average soil and sedimentary-deposit thickness (*29*) | |
| Soil organic carbon content (fine earth fraction) at 0.15m | (*23*) | g per kg | https://www.soilgrids.org ; https://www.isric.org/explore/wosis/accessing-wosis-derived-datasets | (*23*) used data on 150,000 soil profiles worldwide to create a global map, integrating observations within profile to obtain datapoints at fixed depth.A machine learning approach was used, with input data on: Digital Elevation Model-derived characteristics (using the merged DEMS SRTMGL3 and GMTED, as described in (*24*)), long-term average and sd of MODIS-based EVI, bands 4 and 7, long-term monthly average and standard deviation in land surface temperature (MODIS LST), long-term averaged mean monthly hours of snow cover (MODIS 8-day snow occurrence), land cover (using GlobCover30, described in (*25*), who classified Landsat data from 2000 and 2010), precipitation data (averaging WorldClim and GPCP), lithology (using GLiM, (*26*)), Global water table depth (*27*), long-term averaged mean monthly MODIS Flood Water, Landsat-based distribution of mangroves (*28*) and average soil and sedimentary-deposit thickness (*29*) | Biodiversity-Ecosystem Function |
| Soil organic carbon content (fine earth fraction) at 0.30m | (*23*) | g per kg | https://www.soilgrids.org ; https://www.isric.org/explore/wosis/accessing-wosis-derived-datasets | (*23*) used data on 150,000 soil profiles worldwide to create a global map, integrating observations within profile to obtain datapoints at fixed depth.A machine learning approach was used, with input data on: Digital Elevation Model-derived characteristics (using the merged DEMS SRTMGL3 and GMTED, as described in (*24*)), long-term average and sd of MODIS-based EVI, bands 4 and 7, long-term monthly average and standard deviation in land surface temperature (MODIS LST), long-term averaged mean monthly hours of snow cover (MODIS 8-day snow occurrence), land cover (using GlobCover30, described in (*25*), who classified Landsat data from 2000 and 2010), precipitation data (averaging WorldClim and GPCP), lithology (using GLiM, (*26*)), Global water table depth (*27*), long-term averaged mean monthly MODIS Flood Water, Landsat-based distribution of mangroves (*28*) and average soil and sedimentary-deposit thickness (*29*) | Biodiversity-Ecosystem Function |
| Soil organic carbon content (fine earth | (*23*) | g per kg | https://www.soilgrids.org ; https://www.isric.org/explore/wosis/accessin | (*23*) used data on 150,000 soil profiles worldwide to create a global map, integrating observations within profile to obtain datapoints at fixed depth.A machine learning approach was used, with input data on: Digital Elevation Model-derived characteristics (using the merged DEMS SRTMGL3 and GMTED, as described in (*24*)), long-term average and sd of MODIS- | Biodiversity-Ecosystem Function |

| | | | | | |
|---|---|---|---|---|---|
| fraction) at 0.60m | | | g-wosis-derived-datasets | based EVI, bands 4 and 7, long-term monthly average and standard deviation in land surface temperature (MODIS LST), long-term averaged mean monthly hours of snow cover (MODIS 8-day snow occurrence), land cover (using GlobCover30, described in (*25*), who classified Landsat data from 2000 and 2010), precipitation data (averaging WorldClim and GPCP), lithology (using GLiM, (*26*)), Global water table depth (*27*), long-term averaged mean monthly MODIS Flood Water, Landsat-based distribution of mangroves (*28*) and average soil and sedimentary-deposit thickness (*29*) | |
| Soil organic carbon content (fine earth fraction) at 1.00m | (*23*) | g per kg | https://www.soilgrids.org ; https://www.isric.org/explore/wosis/accessing-wosis-derived-datasets | (*23*) used data on 150,000 soil profiles worldwide to create a global map, integrating observations within profile to obtain datapoints at fixed depth.A machine learning approach was used, with input data on: Digital Elevation Model-derived characteristics (using the merged DEMS SRTMGL3 and GMTED, as described in (*24*)), long-term average and sd of MODIS-based EVI, bands 4 and 7, long-term monthly average and standard deviation in land surface temperature (MODIS LST), long-term averaged mean monthly hours of snow cover (MODIS 8-day snow occurrence), land cover (using GlobCover30, described in (*25*), who classified Landsat data from 2000 and 2010), precipitation data (averaging WorldClim and GPCP), lithology (using GLiM, (*26*)), Global water table depth (*27*), long-term averaged mean monthly MODIS Flood Water, Landsat-based distribution of mangroves (*28*) and average soil and sedimentary-deposit thickness (*29*) | Biodiversity-Ecosystem Function |
| Soil organic carbon content (fine earth fraction) at 2.00m | (*23*) | g per kg | https://www.soilgrids.org ; https://www.isric.org/explore/wosis/accessing-wosis-derived-datasets | (*23*) used data on 150,000 soil profiles worldwide to create a global map, integrating observations within profile to obtain datapoints at fixed depth.A machine learning approach was used, with input data on: Digital Elevation Model-derived characteristics (using the merged DEMS SRTMGL3 and GMTED, as described in (*24*)), long-term average and sd of MODIS-based EVI, bands 4 and 7, long-term monthly average and standard deviation in land surface temperature (MODIS LST), long-term averaged mean monthly hours of snow cover (MODIS 8-day snow occurrence), land cover (using GlobCover30, described in (*25*), who classified Landsat data from 2000 and 2010), precipitation data (averaging WorldClim and GPCP), lithology (using GLiM, (*26*)), Global water table depth (*27*), long-term averaged mean monthly MODIS Flood Water, Landsat-based | Biodiversity-Ecosystem Function |

| | | | | | |
|---|---|---|---|---|---|
| | | | | distribution of mangroves (*28*) and average soil and sedimentary-deposit thickness (*29*) | |
| Soil organic carbon density in kg per cubic-m at depth 0.00 m | (*23*) | kg / cubic–meter at depth | https://www.soilgrids.org ; https://www.isric.org/explore/wosis/accessing-wosis-derived-datasets | (*23*) used data on 150,000 soil profiles worldwide to create a global map, integrating observations within profile to obtain datapoints at fixed depth.A machine learning approach was used, with input data on: Digital Elevation Model-derived characteristics (using the merged DEMS SRTMGL3 and GMTED, as described in (*24*)), long-term average and sd of MODIS-based EVI, bands 4 and 7, long-term monthly average and standard deviation in land surface temperature (MODIS LST), long-term averaged mean monthly hours of snow cover (MODIS 8-day snow occurrence), land cover (using GlobCover30, described in (*25*), who classified Landsat data from 2000 and 2010), precipitation data (averaging WorldClim and GPCP), lithology (using GLiM, (*26*)), Global water table depth (*27*), long-term averaged mean monthly MODIS Flood Water, Landsat-based distribution of mangroves (*28*) and average soil and sedimentary-deposit thickness (*29*) | Biodiversity-Ecosystem Function |
| Soil organic carbon density in kg per cubic-m at depth 0.05 m | (*23*) | kg / cubic–meter at depth | https://www.soilgrids.org ; https://www.isric.org/explore/wosis/accessing-wosis-derived-datasets | (*23*) used data on 150,000 soil profiles worldwide to create a global map, integrating observations within profile to obtain datapoints at fixed depth.A machine learning approach was used, with input data on: Digital Elevation Model-derived characteristics (using the merged DEMS SRTMGL3 and GMTED, as described in (*24*)), long-term average and sd of MODIS-based EVI, bands 4 and 7, long-term monthly average and standard deviation in land surface temperature (MODIS LST), long-term averaged mean monthly hours of snow cover (MODIS 8-day snow occurrence), land cover (using GlobCover30, described in (*25*), who classified Landsat data from 2000 and 2010), precipitation data (averaging WorldClim and GPCP), lithology (using GLiM, (*26*)), Global water table depth (*27*), long-term averaged mean monthly MODIS Flood Water, Landsat-based distribution of mangroves (*28*) and average soil and sedimentary-deposit thickness (*29*) | Biodiversity-Ecosystem Function |
| Soil organic carbon density in kg per cubic- | (*23*) | kg / cubic–meter at depth | https://www.soilgrids.org ; https://www.isric.org/explore/wosis/accessin | (*23*) used data on 150,000 soil profiles worldwide to create a global map, integrating observations within profile to obtain datapoints at fixed depth.A machine learning approach was used, with input data on: Digital Elevation Model-derived characteristics (using the merged DEMS SRTMGL3 and GMTED, as described in (*24*)), long-term average and sd of MODIS- | Biodiversity-Ecosystem Function |

| | | | | | |
|---|---|---|---|---|---|
| m at depth 0.15 m | | | g-wosis-derived-datasets | based EVI, bands 4 and 7, long-term monthly average and standard deviation in land surface temperature (MODIS LST), long-term averaged mean monthly hours of snow cover (MODIS 8-day snow occurrence), land cover (using GlobCover30, described in (*25*), who classified Landsat data from 2000 and 2010), precipitation data (averaging WorldClim and GPCP), lithology (using GLiM, (*26*)), Global water table depth (*27*), long-term averaged mean monthly MODIS Flood Water, Landsat-based distribution of mangroves (*28*) and average soil and sedimentary-deposit thickness (*29*) | |
| Soil organic carbon density in kg per cubic-m at depth 0.30 m | (*23*) | kg / cubic–meter at depth | https://www.soilgrids.org ; https://www.isric.org/explore/wosis/accessing-wosis-derived-datasets | (*23*) used data on 150,000 soil profiles worldwide to create a global map, integrating observations within profile to obtain datapoints at fixed depth.A machine learning approach was used, with input data on: Digital Elevation Model-derived characteristics (using the merged DEMS SRTMGL3 and GMTED, as described in (*24*)), long-term average and sd of MODIS-based EVI, bands 4 and 7, long-term monthly average and standard deviation in land surface temperature (MODIS LST), long-term averaged mean monthly hours of snow cover (MODIS 8-day snow occurrence), land cover (using GlobCover30, described in (*25*), who classified Landsat data from 2000 and 2010), precipitation data (averaging WorldClim and GPCP), lithology (using GLiM, (*26*)), Global water table depth (*27*), long-term averaged mean monthly MODIS Flood Water, Landsat-based distribution of mangroves (*28*) and average soil and sedimentary-deposit thickness (*29*) | Biodiversity-Ecosystem Function |
| Soil organic carbon density in kg per cubic-m at depth 0.60 m | (*23*) | kg / cubic–meter at depth | https://www.soilgrids.org ; https://www.isric.org/explore/wosis/accessing-wosis-derived-datasets | (*23*) used data on 150,000 soil profiles worldwide to create a global map, integrating observations within profile to obtain datapoints at fixed depth.A machine learning approach was used, with input data on: Digital Elevation Model-derived characteristics (using the merged DEMS SRTMGL3 and GMTED, as described in (*24*)), long-term average and sd of MODIS-based EVI, bands 4 and 7, long-term monthly average and standard deviation in land surface temperature (MODIS LST), long-term averaged mean monthly hours of snow cover (MODIS 8-day snow occurrence), land cover (using GlobCover30, described in (*25*), who classified Landsat data from 2000 and 2010), precipitation data (averaging WorldClim and GPCP), lithology (using GLiM, (*26*)), Global water table depth (*27*), long-term averaged mean monthly MODIS Flood Water, Landsat-based | Biodiversity-Ecosystem Function |

| | | | | | |
|---|---|---|---|---|---|
| | | | | distribution of mangroves (*28*) and average soil and sedimentary-deposit thickness (*29*) | |
| Soil organic carbon density in kg per cubic-m at depth 1.00 m | (*23*) | kg / cubic–meter at depth | https://www.soilgrids.org ; https://www.isric.org/explore/wosis/accessing-wosis-derived-datasets | (*23*) used data on 150,000 soil profiles worldwide to create a global map, integrating observations within profile to obtain datapoints at fixed depth.A machine learning approach was used, with input data on: Digital Elevation Model-derived characteristics (using the merged DEMS SRTMGL3 and GMTED, as described in (*24*)), long-term average and sd of MODIS-based EVI, bands 4 and 7, long-term monthly average and standard deviation in land surface temperature (MODIS LST), long-term averaged mean monthly hours of snow cover (MODIS 8-day snow occurrence), land cover (using GlobCover30, described in (*25*), who classified Landsat data from 2000 and 2010), precipitation data (averaging WorldClim and GPCP), lithology (using GLiM, (*26*)), Global water table depth (*27*), long-term averaged mean monthly MODIS Flood Water, Landsat-based distribution of mangroves (*28*) and average soil and sedimentary-deposit thickness (*29*) | Biodiversity-Ecosystem Function |
| Soil organic carbon density in kg per cubic-m at depth 2.00 m | (*23*) | kg / cubic–meter at depth | https://www.soilgrids.org ; https://www.isric.org/explore/wosis/accessing-wosis-derived-datasets | (*23*) used data on 150,000 soil profiles worldwide to create a global map, integrating observations within profile to obtain datapoints at fixed depth.A machine learning approach was used, with input data on: Digital Elevation Model-derived characteristics (using the merged DEMS SRTMGL3 and GMTED, as described in (*24*)), long-term average and sd of MODIS-based EVI, bands 4 and 7, long-term monthly average and standard deviation in land surface temperature (MODIS LST), long-term averaged mean monthly hours of snow cover (MODIS 8-day snow occurrence), land cover (using GlobCover30, described in (*25*), who classified Landsat data from 2000 and 2010), precipitation data (averaging WorldClim and GPCP), lithology (using GLiM, (*26*)), Global water table depth (*27*), long-term averaged mean monthly MODIS Flood Water, Landsat-based distribution of mangroves (*28*) and average soil and sedimentary-deposit thickness (*29*) | Biodiversity-Ecosystem Function |
| Soil Organic Carbon Stock from 0.00m-0.05m | (*23*) | tonnes per ha | https://www.soilgrids.org ; https://www.isric.org/explore/wosis/accessin | (*23*) used data on 150,000 soil profiles worldwide to create a global map, integrating observations within profile to obtain datapoints at fixed depth.A machine learning approach was used, with input data on: Digital Elevation Model-derived characteristics (using the merged DEMS SRTMGL3 and GMTED, as described in (*24*)), long-term average and sd of MODIS- | Biodiversity-Ecosystem Function |

| | | | | | |
|---|---|---|---|---|---|
| | | | g-wosis-derived-datasets | based EVI, bands 4 and 7, long-term monthly average and standard deviation in land surface temperature (MODIS LST), long-term averaged mean monthly hours of snow cover (MODIS 8-day snow occurrence), land cover (using GlobCover30, described in (*25*), who classified Landsat data from 2000 and 2010), precipitation data (averaging WorldClim and GPCP), lithology (using GLiM, (*26*)), Global water table depth (*27*), long-term averaged mean monthly MODIS Flood Water, Landsat-based distribution of mangroves (*28*) and average soil and sedimentary-deposit thickness (*29*) | |
| Soil Organic Carbon Stock from 0.05m-0.15m | (*23*) | tonnes per ha | https://www.soilgrids.org ; https://www.isric.org/explore/wosis/accessing-wosis-derived-datasets | (*23*) used data on 150,000 soil profiles worldwide to create a global map, integrating observations within profile to obtain datapoints at fixed depth.A machine learning approach was used, with input data on: Digital Elevation Model-derived characteristics (using the merged DEMS SRTMGL3 and GMTED, as described in (*24*)), long-term average and sd of MODIS-based EVI, bands 4 and 7, long-term monthly average and standard deviation in land surface temperature (MODIS LST), long-term averaged mean monthly hours of snow cover (MODIS 8-day snow occurrence), land cover (using GlobCover30, described in (*25*), who classified Landsat data from 2000 and 2010), precipitation data (averaging WorldClim and GPCP), lithology (using GLiM, (*26*)), Global water table depth (*27*), long-term averaged mean monthly MODIS Flood Water, Landsat-based distribution of mangroves (*28*) and average soil and sedimentary-deposit thickness (*29*) | Biodiversity-Ecosystem Function |
| Soil Organic Carbon Stock from 0.15m-0.30m | (*23*) | tonnes per ha | https://www.soilgrids.org ; https://www.isric.org/explore/wosis/accessing-wosis-derived-datasets | (*23*) used data on 150,000 soil profiles worldwide to create a global map, integrating observations within profile to obtain datapoints at fixed depth.A machine learning approach was used, with input data on: Digital Elevation Model-derived characteristics (using the merged DEMS SRTMGL3 and GMTED, as described in (*24*)), long-term average and sd of MODIS-based EVI, bands 4 and 7, long-term monthly average and standard deviation in land surface temperature (MODIS LST), long-term averaged mean monthly hours of snow cover (MODIS 8-day snow occurrence), land cover (using GlobCover30, described in (*25*), who classified Landsat data from 2000 and 2010), precipitation data (averaging WorldClim and GPCP), lithology (using GLiM, (*26*)), Global water table depth (*27*), long-term averaged mean monthly MODIS Flood Water, Landsat-based | Biodiversity-Ecosystem Function |

| | | | | | |
|---|---|---|---|---|---|
| | | | | distribution of mangroves (*28*) and average soil and sedimentary-deposit thickness (*29*) | |
| Soil Organic Carbon Stock from 0.30m-0.60m | (*23*) | tonnes per ha | https://www.soilgrids.org ; https://www.isric.org/explore/wosis/accessing-wosis-derived-datasets | (*23*) used data on 150,000 soil profiles worldwide to create a global map, integrating observations within profile to obtain datapoints at fixed depth.A machine learning approach was used, with input data on: Digital Elevation Model-derived characteristics (using the merged DEMS SRTMGL3 and GMTED, as described in (*24*)), long-term average and sd of MODIS-based EVI, bands 4 and 7, long-term monthly average and standard deviation in land surface temperature (MODIS LST), long-term averaged mean monthly hours of snow cover (MODIS 8-day snow occurrence), land cover (using GlobCover30, described in (*25*), who classified Landsat data from 2000 and 2010), precipitation data (averaging WorldClim and GPCP), lithology (using GLiM, (*26*)), Global water table depth (*27*), long-term averaged mean monthly MODIS Flood Water, Landsat-based distribution of mangroves (*28*) and average soil and sedimentary-deposit thickness (*29*) | Biodiversity-Ecosystem Function |
| Soil Organic Carbon Stock from 0.60m-1.00m | (*23*) | tonnes per ha | https://www.soilgrids.org ; https://www.isric.org/explore/wosis/accessing-wosis-derived-datasets | (*23*) used data on 150,000 soil profiles worldwide to create a global map, integrating observations within profile to obtain datapoints at fixed depth.A machine learning approach was used, with input data on: Digital Elevation Model-derived characteristics (using the merged DEMS SRTMGL3 and GMTED, as described in (*24*)), long-term average and sd of MODIS-based EVI, bands 4 and 7, long-term monthly average and standard deviation in land surface temperature (MODIS LST), long-term averaged mean monthly hours of snow cover (MODIS 8-day snow occurrence), land cover (using GlobCover30, described in (*25*), who classified Landsat data from 2000 and 2010), precipitation data (averaging WorldClim and GPCP), lithology (using GLiM, (*26*)), Global water table depth (*27*), long-term averaged mean monthly MODIS Flood Water, Landsat-based distribution of mangroves (*28*) and average soil and sedimentary-deposit thickness (*29*) | Biodiversity-Ecosystem Function |
| Organic Carbon Stock from 1.00m-2.00m | (*23*) | tonnes per ha | https://www.soilgrids.org ; https://www.isric.org/explore/wosis/accessin | (*23*) used data on 150,000 soil profiles worldwide to create a global map, integrating observations within profile to obtain datapoints at fixed depth.A machine learning approach was used, with input data on: Digital Elevation Model-derived characteristics (using the merged DEMS SRTMGL3 and GMTED, as described in (*24*)), long-term average and sd of MODIS- | Biodiversity-Ecosystem Function |

| | | | | | |
|---|---|---|---|---|---|
| | | | g-wosis-derived-datasets | based EVI, bands 4 and 7, long-term monthly average and standard deviation in land surface temperature (MODIS LST), long-term averaged mean monthly hours of snow cover (MODIS 8-day snow occurrence), land cover (using GlobCover30, described in (*25*), who classified Landsat data from 2000 and 2010), precipitation data (averaging WorldClim and GPCP), lithology (using GLiM, (*26*)), Global water table depth (*27*), long-term averaged mean monthly MODIS Flood Water, Landsat-based distribution of mangroves (*28*) and average soil and sedimentary-deposit thickness (*29*) | |
| Sand content (50-2000 micro meter) at 0.00m | (*23*) | mass fraction in % | https://www.soilgrids.org ; https://www.isric.org/explore/wosis/accessing-wosis-derived-datasets | (*23*) used data on 150,000 soil profiles worldwide to create a global map, integrating observations within profile to obtain datapoints at fixed depth.A machine learning approach was used, with input data on: Digital Elevation Model-derived characteristics (using the merged DEMS SRTMGL3 and GMTED, as described in (*24*)), long-term average and sd of MODIS-based EVI, bands 4 and 7, long-term monthly average and standard deviation in land surface temperature (MODIS LST), long-term averaged mean monthly hours of snow cover (MODIS 8-day snow occurrence), land cover (using GlobCover30, described in (*25*), who classified Landsat data from 2000 and 2010), precipitation data (averaging WorldClim and GPCP), lithology (using GLiM, (*26*)), Global water table depth (*27*), long-term averaged mean monthly MODIS Flood Water, Landsat-based distribution of mangroves (*28*) and average soil and sedimentary-deposit thickness (*29*) | Biodiversity-Ecosystem Function |
| Sand content (50-2000 micro meter) at 0.05m | (*23*) | mass fraction in % | https://www.soilgrids.org ; https://www.isric.org/explore/wosis/accessing-wosis-derived-datasets | (*23*) used data on 150,000 soil profiles worldwide to create a global map, integrating observations within profile to obtain datapoints at fixed depth.A machine learning approach was used, with input data on: Digital Elevation Model-derived characteristics (using the merged DEMS SRTMGL3 and GMTED, as described in (*24*)), long-term average and sd of MODIS-based EVI, bands 4 and 7, long-term monthly average and standard deviation in land surface temperature (MODIS LST), long-term averaged mean monthly hours of snow cover (MODIS 8-day snow occurrence), land cover (using GlobCover30, described in (*25*), who classified Landsat data from 2000 and 2010), precipitation data (averaging WorldClim and GPCP), lithology (using GLiM, (*26*)), Global water table depth (*27*), long-term averaged mean monthly MODIS Flood Water, Landsat-based | Biodiversity-Ecosystem Function |

| | | | | | |
|---|---|---|---|---|---|
| | | | | distribution of mangroves (*28*) and average soil and sedimentary-deposit thickness (*29*) | |
| Sand content (50-2000 micro meter) at 0.15m | (*23*) | mass fraction in % | https://www.soilgrids.org ; https://www.isric.org/explore/wosis/accessing-wosis-derived-datasets | (*23*) used data on 150,000 soil profiles worldwide to create a global map, integrating observations within profile to obtain datapoints at fixed depth.A machine learning approach was used, with input data on: Digital Elevation Model-derived characteristics (using the merged DEMS SRTMGL3 and GMTED, as described in (*24*)), long-term average and sd of MODIS-based EVI, bands 4 and 7, long-term monthly average and standard deviation in land surface temperature (MODIS LST), long-term averaged mean monthly hours of snow cover (MODIS 8-day snow occurrence), land cover (using GlobCover30, described in (*25*), who classified Landsat data from 2000 and 2010), precipitation data (averaging WorldClim and GPCP), lithology (using GLiM, (*26*)), Global water table depth (*27*), long-term averaged mean monthly MODIS Flood Water, Landsat-based distribution of mangroves (*28*) and average soil and sedimentary-deposit thickness (*29*) | Biodiversity-Ecosystem Function |
| Sand content (50-2000 micro meter) at 0.30m | (*23*) | mass fraction in % | https://www.soilgrids.org ; https://www.isric.org/explore/wosis/accessing-wosis-derived-datasets | (*23*) used data on 150,000 soil profiles worldwide to create a global map, integrating observations within profile to obtain datapoints at fixed depth.A machine learning approach was used, with input data on: Digital Elevation Model-derived characteristics (using the merged DEMS SRTMGL3 and GMTED, as described in (*24*)), long-term average and sd of MODIS-based EVI, bands 4 and 7, long-term monthly average and standard deviation in land surface temperature (MODIS LST), long-term averaged mean monthly hours of snow cover (MODIS 8-day snow occurrence), land cover (using GlobCover30, described in (*25*), who classified Landsat data from 2000 and 2010), precipitation data (averaging WorldClim and GPCP), lithology (using GLiM, (*26*)), Global water table depth (*27*), long-term averaged mean monthly MODIS Flood Water, Landsat-based distribution of mangroves (*28*) and average soil and sedimentary-deposit thickness (*29*) | Biodiversity-Ecosystem Function |
| Sand content (50-2000 micro meter) at 0.60m | (*23*) | mass fraction in % | https://www.soilgrids.org ; https://www.isric.org/explore/wosis/accessin | (*23*) used data on 150,000 soil profiles worldwide to create a global map, integrating observations within profile to obtain datapoints at fixed depth.A machine learning approach was used, with input data on: Digital Elevation Model-derived characteristics (using the merged DEMS SRTMGL3 and GMTED, as described in (*24*)), long-term average and sd of MODIS- | Biodiversity-Ecosystem Function |

| | | | | | |
|---|---|---|---|---|---|
| | | | g-wosis-derived-datasets | based EVI, bands 4 and 7, long-term monthly average and standard deviation in land surface temperature (MODIS LST), long-term averaged mean monthly hours of snow cover (MODIS 8-day snow occurrence), land cover (using GlobCover30, described in (*25*), who classified Landsat data from 2000 and 2010), precipitation data (averaging WorldClim and GPCP), lithology (using GLiM, (*26*)), Global water table depth (*27*), long-term averaged mean monthly MODIS Flood Water, Landsat-based distribution of mangroves (*28*) and average soil and sedimentary-deposit thickness (*29*) | |
| Sand content (50-2000 micro meter) at 1.00m | (*23*) | mass fraction in % | https://www.soilgrids.org ; https://www.isric.org/explore/wosis/accessing-wosis-derived-datasets | (*23*) used data on 150,000 soil profiles worldwide to create a global map, integrating observations within profile to obtain datapoints at fixed depth.A machine learning approach was used, with input data on: Digital Elevation Model-derived characteristics (using the merged DEMS SRTMGL3 and GMTED, as described in (*24*)), long-term average and sd of MODIS-based EVI, bands 4 and 7, long-term monthly average and standard deviation in land surface temperature (MODIS LST), long-term averaged mean monthly hours of snow cover (MODIS 8-day snow occurrence), land cover (using GlobCover30, described in (*25*), who classified Landsat data from 2000 and 2010), precipitation data (averaging WorldClim and GPCP), lithology (using GLiM, (*26*)), Global water table depth (*27*), long-term averaged mean monthly MODIS Flood Water, Landsat-based distribution of mangroves (*28*) and average soil and sedimentary-deposit thickness (*29*) | Biodiversity-Ecosystem Function |
| Sand content (50-2000 micro meter) at 2.00m | (*23*) | mass fraction in % | https://www.soilgrids.org ; https://www.isric.org/explore/wosis/accessing-wosis-derived-datasets | (*23*) used data on 150,000 soil profiles worldwide to create a global map, integrating observations within profile to obtain datapoints at fixed depth.A machine learning approach was used, with input data on: Digital Elevation Model-derived characteristics (using the merged DEMS SRTMGL3 and GMTED, as described in (*24*)), long-term average and sd of MODIS-based EVI, bands 4 and 7, long-term monthly average and standard deviation in land surface temperature (MODIS LST), long-term averaged mean monthly hours of snow cover (MODIS 8-day snow occurrence), land cover (using GlobCover30, described in (*25*), who classified Landsat data from 2000 and 2010), precipitation data (averaging WorldClim and GPCP), lithology (using GLiM, (*26*)), Global water table depth (*27*), long-term averaged mean monthly MODIS Flood Water, Landsat-based | Biodiversity-Ecosystem Function |

| | | | | | |
|---|---|---|---|---|---|
| | | | | distribution of mangroves (*28*) and average soil and sedimentary-deposit thickness (*29*) | |
| Saturated water content (volumetric fraction) for depth 0 cm | (*23*) | % | https://www.soilgrids.org ; https://www.isric.org/explore/wosis/accessing-wosis-derived-datasets | (*23*) used data on 150,000 soil profiles worldwide to create a global map, integrating observations within profile to obtain datapoints at fixed depth.A machine learning approach was used, with input data on: Digital Elevation Model-derived characteristics (using the merged DEMS SRTMGL3 and GMTED, as described in (*24*)), long-term average and sd of MODIS-based EVI, bands 4 and 7, long-term monthly average and standard deviation in land surface temperature (MODIS LST), long-term averaged mean monthly hours of snow cover (MODIS 8-day snow occurrence), land cover (using GlobCover30, described in (*25*), who classified Landsat data from 2000 and 2010), precipitation data (averaging WorldClim and GPCP), lithology (using GLiM, (*26*)), Global water table depth (*27*), long-term averaged mean monthly MODIS Flood Water, Landsat-based distribution of mangroves (*28*) and average soil and sedimentary-deposit thickness (*29*) | Biodiversity-Ecosystem Function |
| Saturated water content (volumetric fraction) for depth 5 cm | (*23*) | % | https://www.soilgrids.org ; https://www.isric.org/explore/wosis/accessing-wosis-derived-datasets | (*23*) used data on 150,000 soil profiles worldwide to create a global map, integrating observations within profile to obtain datapoints at fixed depth.A machine learning approach was used, with input data on: Digital Elevation Model-derived characteristics (using the merged DEMS SRTMGL3 and GMTED, as described in (*24*)), long-term average and sd of MODIS-based EVI, bands 4 and 7, long-term monthly average and standard deviation in land surface temperature (MODIS LST), long-term averaged mean monthly hours of snow cover (MODIS 8-day snow occurrence), land cover (using GlobCover30, described in (*25*), who classified Landsat data from 2000 and 2010), precipitation data (averaging WorldClim and GPCP), lithology (using GLiM, (*26*)), Global water table depth (*27*), long-term averaged mean monthly MODIS Flood Water, Landsat-based distribution of mangroves (*28*) and average soil and sedimentary-deposit thickness (*29*) | Biodiversity-Ecosystem Function |
| Saturated water content (volumetric | (*23*) | % | https://www.soilgrids.org ; https://www.isric.org/explore/wosis/accessin | (*23*) used data on 150,000 soil profiles worldwide to create a global map, integrating observations within profile to obtain datapoints at fixed depth.A machine learning approach was used, with input data on: Digital Elevation Model-derived characteristics (using the merged DEMS SRTMGL3 and GMTED, as described in (*24*)), long-term average and sd of MODIS- | Biodiversity-Ecosystem Function |

| | | | | | |
|---|---|---|---|---|---|
| fraction) for depth 15 cm | | | g-wosis-derived-datasets | based EVI, bands 4 and 7, long-term monthly average and standard deviation in land surface temperature (MODIS LST), long-term averaged mean monthly hours of snow cover (MODIS 8-day snow occurrence), land cover (using GlobCover30, described in (*25*), who classified Landsat data from 2000 and 2010), precipitation data (averaging WorldClim and GPCP), lithology (using GLiM, (*26*)), Global water table depth (*27*), long-term averaged mean monthly MODIS Flood Water, Landsat-based distribution of mangroves (*28*) and average soil and sedimentary-deposit thickness (*29*) | |
| Saturated water content (volumetric fraction) for depth 30 cm | (*23*) | % | https://www.soilgrids.org ; https://www.isric.org/explore/wosis/accessing-wosis-derived-datasets | (*23*) used data on 150,000 soil profiles worldwide to create a global map, integrating observations within profile to obtain datapoints at fixed depth.A machine learning approach was used, with input data on: Digital Elevation Model-derived characteristics (using the merged DEMS SRTMGL3 and GMTED, as described in (*24*)), long-term average and sd of MODIS-based EVI, bands 4 and 7, long-term monthly average and standard deviation in land surface temperature (MODIS LST), long-term averaged mean monthly hours of snow cover (MODIS 8-day snow occurrence), land cover (using GlobCover30, described in (*25*), who classified Landsat data from 2000 and 2010), precipitation data (averaging WorldClim and GPCP), lithology (using GLiM, (*26*)), Global water table depth (*27*), long-term averaged mean monthly MODIS Flood Water, Landsat-based distribution of mangroves (*28*) and average soil and sedimentary-deposit thickness (*29*) | Biodiversity-Ecosystem Function |
| Saturated water content (volumetric fraction) for depth 60 cm | (*23*) | % | https://www.soilgrids.org ; https://www.isric.org/explore/wosis/accessing-wosis-derived-datasets | (*23*) used data on 150,000 soil profiles worldwide to create a global map, integrating observations within profile to obtain datapoints at fixed depth.A machine learning approach was used, with input data on: Digital Elevation Model-derived characteristics (using the merged DEMS SRTMGL3 and GMTED, as described in (*24*)), long-term average and sd of MODIS-based EVI, bands 4 and 7, long-term monthly average and standard deviation in land surface temperature (MODIS LST), long-term averaged mean monthly hours of snow cover (MODIS 8-day snow occurrence), land cover (using GlobCover30, described in (*25*), who classified Landsat data from 2000 and 2010), precipitation data (averaging WorldClim and GPCP), lithology (using GLiM, (*26*)), Global water table depth (*27*), long-term averaged mean monthly MODIS Flood Water, Landsat-based | Biodiversity-Ecosystem Function |

| | | | | | |
|---|---|---|---|---|---|
| | | | | distribution of mangroves (*28*) and average soil and sedimentary-deposit thickness (*29*) | |
| Saturated water content (volumetric fraction) for depth 100 cm | (*23*) | % | https://www.soilgrids.org ; https://www.isric.org/explore/wosis/accessing-wosis-derived-datasets | (*23*) used data on 150,000 soil profiles worldwide to create a global map, integrating observations within profile to obtain datapoints at fixed depth.A machine learning approach was used, with input data on: Digital Elevation Model-derived characteristics (using the merged DEMS SRTMGL3 and GMTED, as described in (*24*)), long-term average and sd of MODIS-based EVI, bands 4 and 7, long-term monthly average and standard deviation in land surface temperature (MODIS LST), long-term averaged mean monthly hours of snow cover (MODIS 8-day snow occurrence), land cover (using GlobCover30, described in (*25*), who classified Landsat data from 2000 and 2010), precipitation data (averaging WorldClim and GPCP), lithology (using GLiM, (*26*)), Global water table depth (*27*), long-term averaged mean monthly MODIS Flood Water, Landsat-based distribution of mangroves (*28*) and average soil and sedimentary-deposit thickness (*29*) | Biodiversity-Ecosystem Function |
| Saturated water content (volumetric fraction) for depth 200 cm | (*23*) | % | https://www.soilgrids.org ; https://www.isric.org/explore/wosis/accessing-wosis-derived-datasets | (*23*) used data on 150,000 soil profiles worldwide to create a global map, integrating observations within profile to obtain datapoints at fixed depth.A machine learning approach was used, with input data on: Digital Elevation Model-derived characteristics (using the merged DEMS SRTMGL3 and GMTED, as described in (*24*)), long-term average and sd of MODIS-based EVI, bands 4 and 7, long-term monthly average and standard deviation in land surface temperature (MODIS LST), long-term averaged mean monthly hours of snow cover (MODIS 8-day snow occurrence), land cover (using GlobCover30, described in (*25*), who classified Landsat data from 2000 and 2010), precipitation data (averaging WorldClim and GPCP), lithology (using GLiM, (*26*)), Global water table depth (*27*), long-term averaged mean monthly MODIS Flood Water, Landsat-based distribution of mangroves (*28*) and average soil and sedimentary-deposit thickness (*29*) | Biodiversity-Ecosystem Function |
| Silt content (2-50 micro meter) at 0.00m | (*23*) | mass fraction in % | https://www.soilgrids.org ; https://www.isric.org/explore/wosis/accessin | (*23*) used data on 150,000 soil profiles worldwide to create a global map, integrating observations within profile to obtain datapoints at fixed depth.A machine learning approach was used, with input data on: Digital Elevation Model-derived characteristics (using the merged DEMS SRTMGL3 and GMTED, as described in (*24*)), long-term average and sd of MODIS- | Biodiversity-Ecosystem Function |

| | | | | | |
|---|---|---|---|---|---|
| | | | g-wosis-derived-datasets | based EVI, bands 4 and 7, long-term monthly average and standard deviation in land surface temperature (MODIS LST), long-term averaged mean monthly hours of snow cover (MODIS 8-day snow occurrence), land cover (using GlobCover30, described in (*25*), who classified Landsat data from 2000 and 2010), precipitation data (averaging WorldClim and GPCP), lithology (using GLiM, (*26*)), Global water table depth (*27*), long-term averaged mean monthly MODIS Flood Water, Landsat-based distribution of mangroves (*28*) and average soil and sedimentary-deposit thickness (*29*) | |
| Silt content (2-50 micro meter) at 0.05m | (*23*) | mass fraction in % | https://www.soilgrids.org ; https://www.isric.org/explore/wosis/accessing-wosis-derived-datasets | (*23*) used data on 150,000 soil profiles worldwide to create a global map, integrating observations within profile to obtain datapoints at fixed depth.A machine learning approach was used, with input data on: Digital Elevation Model-derived characteristics (using the merged DEMS SRTMGL3 and GMTED, as described in (*24*)), long-term average and sd of MODIS-based EVI, bands 4 and 7, long-term monthly average and standard deviation in land surface temperature (MODIS LST), long-term averaged mean monthly hours of snow cover (MODIS 8-day snow occurrence), land cover (using GlobCover30, described in (*25*), who classified Landsat data from 2000 and 2010), precipitation data (averaging WorldClim and GPCP), lithology (using GLiM, (*26*)), Global water table depth (*27*), long-term averaged mean monthly MODIS Flood Water, Landsat-based distribution of mangroves (*28*) and average soil and sedimentary-deposit thickness (*29*) | Biodiversity-Ecosystem Function |
| Silt content (2-50 micro meter) at 0.15m | (*23*) | mass fraction in % | https://www.soilgrids.org ; https://www.isric.org/explore/wosis/accessing-wosis-derived-datasets | (*23*) used data on 150,000 soil profiles worldwide to create a global map, integrating observations within profile to obtain datapoints at fixed depth.A machine learning approach was used, with input data on: Digital Elevation Model-derived characteristics (using the merged DEMS SRTMGL3 and GMTED, as described in (*24*)), long-term average and sd of MODIS-based EVI, bands 4 and 7, long-term monthly average and standard deviation in land surface temperature (MODIS LST), long-term averaged mean monthly hours of snow cover (MODIS 8-day snow occurrence), land cover (using GlobCover30, described in (*25*), who classified Landsat data from 2000 and 2010), precipitation data (averaging WorldClim and GPCP), lithology (using GLiM, (*26*)), Global water table depth (*27*), long-term averaged mean monthly MODIS Flood Water, Landsat-based | Biodiversity-Ecosystem Function |

| | | | | | |
|---|---|---|---|---|---|
| | | | | distribution of mangroves (*28*) and average soil and sedimentary-deposit thickness (*29*) | |
| Silt content (2-50 micro meter) at 0.30m | (*23*) | mass fraction in % | https://www.soilgrids.org ; https://www.isric.org/explore/wosis/accessing-wosis-derived-datasets | (*23*) used data on 150,000 soil profiles worldwide to create a global map, integrating observations within profile to obtain datapoints at fixed depth.A machine learning approach was used, with input data on: Digital Elevation Model-derived characteristics (using the merged DEMS SRTMGL3 and GMTED, as described in (*24*)), long-term average and sd of MODIS-based EVI, bands 4 and 7, long-term monthly average and standard deviation in land surface temperature (MODIS LST), long-term averaged mean monthly hours of snow cover (MODIS 8-day snow occurrence), land cover (using GlobCover30, described in (*25*), who classified Landsat data from 2000 and 2010), precipitation data (averaging WorldClim and GPCP), lithology (using GLiM, (*26*)), Global water table depth (*27*), long-term averaged mean monthly MODIS Flood Water, Landsat-based distribution of mangroves (*28*) and average soil and sedimentary-deposit thickness (*29*) | Biodiversity-Ecosystem Function |
| Silt content (2-50 micro meter) at 0.60m | (*23*) | mass fraction in % | https://www.soilgrids.org ; https://www.isric.org/explore/wosis/accessing-wosis-derived-datasets | (*23*) used data on 150,000 soil profiles worldwide to create a global map, integrating observations within profile to obtain datapoints at fixed depth.A machine learning approach was used, with input data on: Digital Elevation Model-derived characteristics (using the merged DEMS SRTMGL3 and GMTED, as described in (*24*)), long-term average and sd of MODIS-based EVI, bands 4 and 7, long-term monthly average and standard deviation in land surface temperature (MODIS LST), long-term averaged mean monthly hours of snow cover (MODIS 8-day snow occurrence), land cover (using GlobCover30, described in (*25*), who classified Landsat data from 2000 and 2010), precipitation data (averaging WorldClim and GPCP), lithology (using GLiM, (*26*)), Global water table depth (*27*), long-term averaged mean monthly MODIS Flood Water, Landsat-based distribution of mangroves (*28*) and average soil and sedimentary-deposit thickness (*29*) | Biodiversity-Ecosystem Function |
| Silt content (2-50 micro meter) at 1.00m | (*23*) | mass fraction in % | https://www.soilgrids.org ; https://www.isric.org/explore/wosis/accessin | (*23*) used data on 150,000 soil profiles worldwide to create a global map, integrating observations within profile to obtain datapoints at fixed depth.A machine learning approach was used, with input data on: Digital Elevation Model-derived characteristics (using the merged DEMS SRTMGL3 and GMTED, as described in (*24*)), long-term average and sd of MODIS- | Biodiversity-Ecosystem Function |

| | | | | | |
|---|---|---|---|---|---|
| | | | g-wosis-derived-datasets | based EVI, bands 4 and 7, long-term monthly average and standard deviation in land surface temperature (MODIS LST), long-term averaged mean monthly hours of snow cover (MODIS 8-day snow occurrence), land cover (using GlobCover30, described in (*25*), who classified Landsat data from 2000 and 2010), precipitation data (averaging WorldClim and GPCP), lithology (using GLiM, (*26*)), Global water table depth (*27*), long-term averaged mean monthly MODIS Flood Water, Landsat-based distribution of mangroves (*28*) and average soil and sedimentary-deposit thickness (*29*) | |
| Silt content (2-50 micro meter) at 2.00m | (*23*) | mass fraction in % | https://www.soilgrids.org ; https://www.isric.org/explore/wosis/accessing-wosis-derived-datasets | (*23*) used data on 150,000 soil profiles worldwide to create a global map, integrating observations within profile to obtain datapoints at fixed depth.A machine learning approach was used, with input data on: Digital Elevation Model-derived characteristics (using the merged DEMS SRTMGL3 and GMTED, as described in (*24*)), long-term average and sd of MODIS-based EVI, bands 4 and 7, long-term monthly average and standard deviation in land surface temperature (MODIS LST), long-term averaged mean monthly hours of snow cover (MODIS 8-day snow occurrence), land cover (using GlobCover30, described in (*25*), who classified Landsat data from 2000 and 2010), precipitation data (averaging WorldClim and GPCP), lithology (using GLiM, (*26*)), Global water table depth (*27*), long-term averaged mean monthly MODIS Flood Water, Landsat-based distribution of mangroves (*28*) and average soil and sedimentary-deposit thickness (*29*) | Biodiversity-Ecosystem Function |
| Available soil water capacity (volumetric fraction) depth 0 cm | (*23*) | % | https://www.soilgrids.org ; https://www.isric.org/explore/wosis/accessing-wosis-derived-datasets | (*23*) used data on 150,000 soil profiles worldwide to create a global map, integrating observations within profile to obtain datapoints at fixed depth.A machine learning approach was used, with input data on: Digital Elevation Model-derived characteristics (using the merged DEMS SRTMGL3 and GMTED, as described in (*24*)), long-term average and sd of MODIS-based EVI, bands 4 and 7, long-term monthly average and standard deviation in land surface temperature (MODIS LST), long-term averaged mean monthly hours of snow cover (MODIS 8-day snow occurrence), land cover (using GlobCover30, described in (*25*), who classified Landsat data from 2000 and 2010), precipitation data (averaging WorldClim and GPCP), lithology (using GLiM, (*26*)), Global water table depth (*27*), long-term averaged mean monthly MODIS Flood Water, Landsat-based | Biodiversity-Ecosystem Function |

| | | | | | |
|---|---|---|---|---|---|
| | | | | distribution of mangroves (*28*) and average soil and sedimentary-deposit thickness (*29*) | |
| Available soil water capacity (volumetric fraction) for depth 5 cm | (*23*) | % | https://www.soilgrids.org ; https://www.isric.org/explore/wosis/accessing-wosis-derived-datasets | (*23*) used data on 150,000 soil profiles worldwide to create a global map, integrating observations within profile to obtain datapoints at fixed depth.A machine learning approach was used, with input data on: Digital Elevation Model-derived characteristics (using the merged DEMS SRTMGL3 and GMTED, as described in (*24*)), long-term average and sd of MODIS-based EVI, bands 4 and 7, long-term monthly average and standard deviation in land surface temperature (MODIS LST), long-term averaged mean monthly hours of snow cover (MODIS 8-day snow occurrence), land cover (using GlobCover30, described in (*25*), who classified Landsat data from 2000 and 2010), precipitation data (averaging WorldClim and GPCP), lithology (using GLiM, (*26*)), Global water table depth (*27*), long-term averaged mean monthly MODIS Flood Water, Landsat-based distribution of mangroves (*28*) and average soil and sedimentary-deposit thickness (*29*) | Biodiversity-Ecosystem Function |
| Available soil water capacity (volumetric fraction) for depth 15 cm | (*23*) | % | https://www.soilgrids.org ; https://www.isric.org/explore/wosis/accessing-wosis-derived-datasets | (*23*) used data on 150,000 soil profiles worldwide to create a global map, integrating observations within profile to obtain datapoints at fixed depth.A machine learning approach was used, with input data on: Digital Elevation Model-derived characteristics (using the merged DEMS SRTMGL3 and GMTED, as described in (*24*)), long-term average and sd of MODIS-based EVI, bands 4 and 7, long-term monthly average and standard deviation in land surface temperature (MODIS LST), long-term averaged mean monthly hours of snow cover (MODIS 8-day snow occurrence), land cover (using GlobCover30, described in (*25*), who classified Landsat data from 2000 and 2010), precipitation data (averaging WorldClim and GPCP), lithology (using GLiM, (*26*)), Global water table depth (*27*), long-term averaged mean monthly MODIS Flood Water, Landsat-based distribution of mangroves (*28*) and average soil and sedimentary-deposit thickness (*29*) | Biodiversity-Ecosystem Function |
| Available soil water capacity (volumetric | (*23*) | % | https://www.soilgrids.org ; https://www.isric.org/explore/wosis/accessin | (*23*) used data on 150,000 soil profiles worldwide to create a global map, integrating observations within profile to obtain datapoints at fixed depth.A machine learning approach was used, with input data on: Digital Elevation Model-derived characteristics (using the merged DEMS SRTMGL3 and GMTED, as described in (*24*)), long-term average and sd of MODIS- | Biodiversity-Ecosystem Function |

| | | | | | |
|---|---|---|---|---|---|
| fraction) for depth 30 cm | | | g-wosis-derived-datasets | based EVI, bands 4 and 7, long-term monthly average and standard deviation in land surface temperature (MODIS LST), long-term averaged mean monthly hours of snow cover (MODIS 8-day snow occurrence), land cover (using GlobCover30, described in (*25*), who classified Landsat data from 2000 and 2010), precipitation data (averaging WorldClim and GPCP), lithology (using GLiM, (*26*)), Global water table depth (*27*), long-term averaged mean monthly MODIS Flood Water, Landsat-based distribution of mangroves (*28*) and average soil and sedimentary-deposit thickness (*29*) | |
| Available soil water capacity (volumetric fraction) for depth 60 cm | (*23*) | % | https://www.soilgrids.org ; https://www.isric.org/explore/wosis/accessing-wosis-derived-datasets | (*23*) used data on 150,000 soil profiles worldwide to create a global map, integrating observations within profile to obtain datapoints at fixed depth.A machine learning approach was used, with input data on: Digital Elevation Model-derived characteristics (using the merged DEMS SRTMGL3 and GMTED, as described in (*24*)), long-term average and sd of MODIS-based EVI, bands 4 and 7, long-term monthly average and standard deviation in land surface temperature (MODIS LST), long-term averaged mean monthly hours of snow cover (MODIS 8-day snow occurrence), land cover (using GlobCover30, described in (*25*), who classified Landsat data from 2000 and 2010), precipitation data (averaging WorldClim and GPCP), lithology (using GLiM, (*26*)), Global water table depth (*27*), long-term averaged mean monthly MODIS Flood Water, Landsat-based distribution of mangroves (*28*) and average soil and sedimentary-deposit thickness (*29*) | Biodiversity-Ecosystem Function |
| Available soil water capacity (volumetric fraction) for depth 100 cm | (*23*) | % | https://www.soilgrids.org ; https://www.isric.org/explore/wosis/accessing-wosis-derived-datasets | (*23*) used data on 150,000 soil profiles worldwide to create a global map, integrating observations within profile to obtain datapoints at fixed depth.A machine learning approach was used, with input data on: Digital Elevation Model-derived characteristics (using the merged DEMS SRTMGL3 and GMTED, as described in (*24*)), long-term average and sd of MODIS-based EVI, bands 4 and 7, long-term monthly average and standard deviation in land surface temperature (MODIS LST), long-term averaged mean monthly hours of snow cover (MODIS 8-day snow occurrence), land cover (using GlobCover30, described in (*25*), who classified Landsat data from 2000 and 2010), precipitation data (averaging WorldClim and GPCP), lithology (using GLiM, (*26*)), Global water table depth (*27*), long-term averaged mean monthly MODIS Flood Water, Landsat-based | Biodiversity-Ecosystem Function |

| | | | | | |
|---|---|---|---|---|---|
| | | | | distribution of mangroves (*28*) and average soil and sedimentary-deposit thickness (*29*) | |
| Available soil water capacity (volumetric fraction) for depth 200 cm | (*23*) | % | https://www.soilgrids.org ; https://www.isric.org/explore/wosis/accessing-wosis-derived-datasets | (*23*) used data on 150,000 soil profiles worldwide to create a global map, integrating observations within profile to obtain datapoints at fixed depth.A machine learning approach was used, with input data on: Digital Elevation Model-derived characteristics (using the merged DEMS SRTMGL3 and GMTED, as described in (*24*)), long-term average and sd of MODIS-based EVI, bands 4 and 7, long-term monthly average and standard deviation in land surface temperature (MODIS LST), long-term averaged mean monthly hours of snow cover (MODIS 8-day snow occurrence), land cover (using GlobCover30, described in (*25*), who classified Landsat data from 2000 and 2010), precipitation data (averaging WorldClim and GPCP), lithology (using GLiM, (*26*)), Global water table depth (*27*), long-term averaged mean monthly MODIS Flood Water, Landsat-based distribution of mangroves (*28*) and average soil and sedimentary-deposit thickness (*29*) | Biodiversity-Ecosystem Function |
| Available soil water capacity (volumetric fraction) for depth 0 cm | (*23*) | % | https://www.soilgrids.org ; https://www.isric.org/explore/wosis/accessing-wosis-derived-datasets | (*23*) used data on 150,000 soil profiles worldwide to create a global map, integrating observations within profile to obtain datapoints at fixed depth.A machine learning approach was used, with input data on: Digital Elevation Model-derived characteristics (using the merged DEMS SRTMGL3 and GMTED, as described in (*24*)), long-term average and sd of MODIS-based EVI, bands 4 and 7, long-term monthly average and standard deviation in land surface temperature (MODIS LST), long-term averaged mean monthly hours of snow cover (MODIS 8-day snow occurrence), land cover (using GlobCover30, described in (*25*), who classified Landsat data from 2000 and 2010), precipitation data (averaging WorldClim and GPCP), lithology (using GLiM, (*26*)), Global water table depth (*27*), long-term averaged mean monthly MODIS Flood Water, Landsat-based distribution of mangroves (*28*) and average soil and sedimentary-deposit thickness (*29*) | Biodiversity-Ecosystem Function |
| Available soil water capacity (volumetric | (*23*) | % | https://www.soilgrids.org ; https://www.isric.org/explore/wosis/accessin | (*23*) used data on 150,000 soil profiles worldwide to create a global map, integrating observations within profile to obtain datapoints at fixed depth.A machine learning approach was used, with input data on: Digital Elevation Model-derived characteristics (using the merged DEMS SRTMGL3 and GMTED, as described in (*24*)), long-term average and sd of MODIS- | Biodiversity-Ecosystem Function |

| | | | | | |
|---|---|---|---|---|---|
| fraction) for depth 5 cm | | | g-wosis-derived-datasets | based EVI, bands 4 and 7, long-term monthly average and standard deviation in land surface temperature (MODIS LST), long-term averaged mean monthly hours of snow cover (MODIS 8-day snow occurrence), land cover (using GlobCover30, described in (*25*), who classified Landsat data from 2000 and 2010), precipitation data (averaging WorldClim and GPCP), lithology (using GLiM, (*26*)), Global water table depth (*27*), long-term averaged mean monthly MODIS Flood Water, Landsat-based distribution of mangroves (*28*) and average soil and sedimentary-deposit thickness (*29*) | |
| Available soil water capacity (volumetric fraction) for depth 15 cm | (*23*) | % | https://www.soilgrids.org ; https://www.isric.org/explore/wosis/accessing-wosis-derived-datasets | (*23*) used data on 150,000 soil profiles worldwide to create a global map, integrating observations within profile to obtain datapoints at fixed depth.A machine learning approach was used, with input data on: Digital Elevation Model-derived characteristics (using the merged DEMS SRTMGL3 and GMTED, as described in (*24*)), long-term average and sd of MODIS-based EVI, bands 4 and 7, long-term monthly average and standard deviation in land surface temperature (MODIS LST), long-term averaged mean monthly hours of snow cover (MODIS 8-day snow occurrence), land cover (using GlobCover30, described in (*25*), who classified Landsat data from 2000 and 2010), precipitation data (averaging WorldClim and GPCP), lithology (using GLiM, (*26*)), Global water table depth (*27*), long-term averaged mean monthly MODIS Flood Water, Landsat-based distribution of mangroves (*28*) and average soil and sedimentary-deposit thickness (*29*) | Biodiversity-Ecosystem Function |
| Available soil water capacity (volumetric fraction) for depth 30 cm | (*23*) | % | https://www.soilgrids.org ; https://www.isric.org/explore/wosis/accessing-wosis-derived-datasets | (*23*) used data on 150,000 soil profiles worldwide to create a global map, integrating observations within profile to obtain datapoints at fixed depth.A machine learning approach was used, with input data on: Digital Elevation Model-derived characteristics (using the merged DEMS SRTMGL3 and GMTED, as described in (*24*)), long-term average and sd of MODIS-based EVI, bands 4 and 7, long-term monthly average and standard deviation in land surface temperature (MODIS LST), long-term averaged mean monthly hours of snow cover (MODIS 8-day snow occurrence), land cover (using GlobCover30, described in (*25*), who classified Landsat data from 2000 and 2010), precipitation data (averaging WorldClim and GPCP), lithology (using GLiM, (*26*)), Global water table depth (*27*), long-term averaged mean monthly MODIS Flood Water, Landsat-based | Biodiversity-Ecosystem Function |

| | | | | | |
|---|---|---|---|---|---|
| | | | | distribution of mangroves (*28*) and average soil and sedimentary-deposit thickness (*29*) | |
| Available soil water capacity (volumetric fraction) for depth 60 cm | (*23*) | % | https://www.soilgrids.org ; https://www.isric.org/explore/wosis/accessing-wosis-derived-datasets | (*23*) used data on 150,000 soil profiles worldwide to create a global map, integrating observations within profile to obtain datapoints at fixed depth.A machine learning approach was used, with input data on: Digital Elevation Model-derived characteristics (using the merged DEMS SRTMGL3 and GMTED, as described in (*24*)), long-term average and sd of MODIS-based EVI, bands 4 and 7, long-term monthly average and standard deviation in land surface temperature (MODIS LST), long-term averaged mean monthly hours of snow cover (MODIS 8-day snow occurrence), land cover (using GlobCover30, described in (*25*), who classified Landsat data from 2000 and 2010), precipitation data (averaging WorldClim and GPCP), lithology (using GLiM, (*26*)), Global water table depth (*27*), long-term averaged mean monthly MODIS Flood Water, Landsat-based distribution of mangroves (*28*) and average soil and sedimentary-deposit thickness (*29*) | Biodiversity-Ecosystem Function |
| Available soil water capacity (volumetric fraction) for depth 100 cm | (*23*) | % | https://www.soilgrids.org ; https://www.isric.org/explore/wosis/accessing-wosis-derived-datasets | (*23*) used data on 150,000 soil profiles worldwide to create a global map, integrating observations within profile to obtain datapoints at fixed depth.A machine learning approach was used, with input data on: Digital Elevation Model-derived characteristics (using the merged DEMS SRTMGL3 and GMTED, as described in (*24*)), long-term average and sd of MODIS-based EVI, bands 4 and 7, long-term monthly average and standard deviation in land surface temperature (MODIS LST), long-term averaged mean monthly hours of snow cover (MODIS 8-day snow occurrence), land cover (using GlobCover30, described in (*25*), who classified Landsat data from 2000 and 2010), precipitation data (averaging WorldClim and GPCP), lithology (using GLiM, (*26*)), Global water table depth (*27*), long-term averaged mean monthly MODIS Flood Water, Landsat-based distribution of mangroves (*28*) and average soil and sedimentary-deposit thickness (*29*) | Biodiversity-Ecosystem Function |
| Available soil water capacity (volumetric | (*23*) | % | https://www.soilgrids.org ; https://www.isric.org/explore/wosis/accessin | (*23*) used data on 150,000 soil profiles worldwide to create a global map, integrating observations within profile to obtain datapoints at fixed depth.A machine learning approach was used, with input data on: Digital Elevation Model-derived characteristics (using the merged DEMS SRTMGL3 and GMTED, as described in (*24*)), long-term average and sd of MODIS- | Biodiversity-Ecosystem Function |

| | | | | | |
|---|---|---|---|---|---|
| fraction) for depth 200 cm | | | g-wosis-derived-datasets | based EVI, bands 4 and 7, long-term monthly average and standard deviation in land surface temperature (MODIS LST), long-term averaged mean monthly hours of snow cover (MODIS 8-day snow occurrence), land cover (using GlobCover30, described in (*25*), who classified Landsat data from 2000 and 2010), precipitation data (averaging WorldClim and GPCP), lithology (using GLiM, (*26*)), Global water table depth (*27*), long-term averaged mean monthly MODIS Flood Water, Landsat-based distribution of mangroves (*28*) and average soil and sedimentary-deposit thickness (*29*) | |
| Available soil water capacity (volumetric fraction) for depth 0 cm | (*23*) | % | https://www.soilgrids.org ; https://www.isric.org/explore/wosis/accessing-wosis-derived-datasets | (*23*) used data on 150,000 soil profiles worldwide to create a global map, integrating observations within profile to obtain datapoints at fixed depth.A machine learning approach was used, with input data on: Digital Elevation Model-derived characteristics (using the merged DEMS SRTMGL3 and GMTED, as described in (*24*)), long-term average and sd of MODIS-based EVI, bands 4 and 7, long-term monthly average and standard deviation in land surface temperature (MODIS LST), long-term averaged mean monthly hours of snow cover (MODIS 8-day snow occurrence), land cover (using GlobCover30, described in (*25*), who classified Landsat data from 2000 and 2010), precipitation data (averaging WorldClim and GPCP), lithology (using GLiM, (*26*)), Global water table depth (*27*), long-term averaged mean monthly MODIS Flood Water, Landsat-based distribution of mangroves (*28*) and average soil and sedimentary-deposit thickness (*29*) | Biodiversity-Ecosystem Function |
| Available soil water capacity (volumetric fraction) for depth 5 cm | (*23*) | % | https://www.soilgrids.org ; https://www.isric.org/explore/wosis/accessing-wosis-derived-datasets | (*23*) used data on 150,000 soil profiles worldwide to create a global map, integrating observations within profile to obtain datapoints at fixed depth.A machine learning approach was used, with input data on: Digital Elevation Model-derived characteristics (using the merged DEMS SRTMGL3 and GMTED, as described in (*24*)), long-term average and sd of MODIS-based EVI, bands 4 and 7, long-term monthly average and standard deviation in land surface temperature (MODIS LST), long-term averaged mean monthly hours of snow cover (MODIS 8-day snow occurrence), land cover (using GlobCover30, described in (*25*), who classified Landsat data from 2000 and 2010), precipitation data (averaging WorldClim and GPCP), lithology (using GLiM, (*26*)), Global water table depth (*27*), long-term averaged mean monthly MODIS Flood Water, Landsat-based | Biodiversity-Ecosystem Function |

| | | | | | |
|---|---|---|---|---|---|
| | | | | distribution of mangroves (*28*) and average soil and sedimentary-deposit thickness (*29*) | |
| Available soil water capacity (volumetric fraction) for depth 15 cm | (*23*) | % | https://www.soilgrids.org ; https://www.isric.org/explore/wosis/accessing-wosis-derived-datasets | (*23*) used data on 150,000 soil profiles worldwide to create a global map, integrating observations within profile to obtain datapoints at fixed depth.A machine learning approach was used, with input data on: Digital Elevation Model-derived characteristics (using the merged DEMS SRTMGL3 and GMTED, as described in (*24*)), long-term average and sd of MODIS-based EVI, bands 4 and 7, long-term monthly average and standard deviation in land surface temperature (MODIS LST), long-term averaged mean monthly hours of snow cover (MODIS 8-day snow occurrence), land cover (using GlobCover30, described in (*25*), who classified Landsat data from 2000 and 2010), precipitation data (averaging WorldClim and GPCP), lithology (using GLiM, (*26*)), Global water table depth (*27*), long-term averaged mean monthly MODIS Flood Water, Landsat-based distribution of mangroves (*28*) and average soil and sedimentary-deposit thickness (*29*) | Biodiversity-Ecosystem Function |
| Available soil water capacity (volumetric fraction) for depth 30 cm | (*23*) | % | https://www.soilgrids.org ; https://www.isric.org/explore/wosis/accessing-wosis-derived-datasets | (*23*) used data on 150,000 soil profiles worldwide to create a global map, integrating observations within profile to obtain datapoints at fixed depth.A machine learning approach was used, with input data on: Digital Elevation Model-derived characteristics (using the merged DEMS SRTMGL3 and GMTED, as described in (*24*)), long-term average and sd of MODIS-based EVI, bands 4 and 7, long-term monthly average and standard deviation in land surface temperature (MODIS LST), long-term averaged mean monthly hours of snow cover (MODIS 8-day snow occurrence), land cover (using GlobCover30, described in (*25*), who classified Landsat data from 2000 and 2010), precipitation data (averaging WorldClim and GPCP), lithology (using GLiM, (*26*)), Global water table depth (*27*), long-term averaged mean monthly MODIS Flood Water, Landsat-based distribution of mangroves (*28*) and average soil and sedimentary-deposit thickness (*29*) | Biodiversity-Ecosystem Function |
| Available soil water capacity (volumetric | (*23*) | % | https://www.soilgrids.org ; https://www.isric.org/explore/wosis/accessin | (*23*) used data on 150,000 soil profiles worldwide to create a global map, integrating observations within profile to obtain datapoints at fixed depth.A machine learning approach was used, with input data on: Digital Elevation Model-derived characteristics (using the merged DEMS SRTMGL3 and GMTED, as described in (*24*)), long-term average and sd of MODIS- | Biodiversity-Ecosystem Function |

| | | | | | |
|---|---|---|---|---|---|
| fraction) for depth 60 cm | | | g-wosis-derived-datasets | based EVI, bands 4 and 7, long-term monthly average and standard deviation in land surface temperature (MODIS LST), long-term averaged mean monthly hours of snow cover (MODIS 8-day snow occurrence), land cover (using GlobCover30, described in (*25*), who classified Landsat data from 2000 and 2010), precipitation data (averaging WorldClim and GPCP), lithology (using GLiM, (*26*)), Global water table depth (*27*), long-term averaged mean monthly MODIS Flood Water, Landsat-based distribution of mangroves (*28*) and average soil and sedimentary-deposit thickness (*29*) | |
| Available soil water capacity (volumetric fraction) for depth 100 cm | (*23*) | % | https://www.soilgrids.org ; https://www.isric.org/explore/wosis/accessing-wosis-derived-datasets | (*23*) used data on 150,000 soil profiles worldwide to create a global map, integrating observations within profile to obtain datapoints at fixed depth.A machine learning approach was used, with input data on: Digital Elevation Model-derived characteristics (using the merged DEMS SRTMGL3 and GMTED, as described in (*24*)), long-term average and sd of MODIS-based EVI, bands 4 and 7, long-term monthly average and standard deviation in land surface temperature (MODIS LST), long-term averaged mean monthly hours of snow cover (MODIS 8-day snow occurrence), land cover (using GlobCover30, described in (*25*), who classified Landsat data from 2000 and 2010), precipitation data (averaging WorldClim and GPCP), lithology (using GLiM, (*26*)), Global water table depth (*27*), long-term averaged mean monthly MODIS Flood Water, Landsat-based distribution of mangroves (*28*) and average soil and sedimentary-deposit thickness (*29*) | Biodiversity-Ecosystem Function |
| Available soil water capacity (volumetric fraction) for depth 200 cm | (*23*) | % | https://www.soilgrids.org ; https://www.isric.org/explore/wosis/accessing-wosis-derived-datasets | (*23*) used data on 150,000 soil profiles worldwide to create a global map, integrating observations within profile to obtain datapoints at fixed depth.A machine learning approach was used, with input data on: Digital Elevation Model-derived characteristics (using the merged DEMS SRTMGL3 and GMTED, as described in (*24*)), long-term average and sd of MODIS-based EVI, bands 4 and 7, long-term monthly average and standard deviation in land surface temperature (MODIS LST), long-term averaged mean monthly hours of snow cover (MODIS 8-day snow occurrence), land cover (using GlobCover30, described in (*25*), who classified Landsat data from 2000 and 2010), precipitation data (averaging WorldClim and GPCP), lithology (using GLiM, (*26*)), Global water table depth (*27*), long-term averaged mean monthly MODIS Flood Water, Landsat-based | Biodiversity-Ecosystem Function |

| | | | | | |
|---|---|---|---|---|---|
| | | | | distribution of mangroves (*28*) and average soil and sedimentary-deposit thickness (*29*) | |
| Soil pH x 10 in H2O at 0.00m | (*23*) | pH x 10 | https://www.soilgrids.org ; https://www.isric.org/explore/wosis/accessing-wosis-derived-datasets | (*23*) used data on 150,000 soil profiles worldwide to create a global map, integrating observations within profile to obtain datapoints at fixed depth.A machine learning approach was used, with input data on: Digital Elevation Model-derived characteristics (using the merged DEMS SRTMGL3 and GMTED, as described in (*24*)), long-term average and sd of MODIS-based EVI, bands 4 and 7, long-term monthly average and standard deviation in land surface temperature (MODIS LST), long-term averaged mean monthly hours of snow cover (MODIS 8-day snow occurrence), land cover (using GlobCover30, described in (*25*), who classified Landsat data from 2000 and 2010), precipitation data (averaging WorldClim and GPCP), lithology (using GLiM, (*26*)), Global water table depth (*27*), long-term averaged mean monthly MODIS Flood Water, Landsat-based distribution of mangroves (*28*) and average soil and sedimentary-deposit thickness (*29*) | Biodiversity-Ecosystem Function |
| Soil pH x 10 in H2O at 0.05m | (*23*) | pH x 10 | https://www.soilgrids.org ; https://www.isric.org/explore/wosis/accessing-wosis-derived-datasets | (*23*) used data on 150,000 soil profiles worldwide to create a global map, integrating observations within profile to obtain datapoints at fixed depth.A machine learning approach was used, with input data on: Digital Elevation Model-derived characteristics (using the merged DEMS SRTMGL3 and GMTED, as described in (*24*)), long-term average and sd of MODIS-based EVI, bands 4 and 7, long-term monthly average and standard deviation in land surface temperature (MODIS LST), long-term averaged mean monthly hours of snow cover (MODIS 8-day snow occurrence), land cover (using GlobCover30, described in (*25*), who classified Landsat data from 2000 and 2010), precipitation data (averaging WorldClim and GPCP), lithology (using GLiM, (*26*)), Global water table depth (*27*), long-term averaged mean monthly MODIS Flood Water, Landsat-based distribution of mangroves (*28*) and average soil and sedimentary-deposit thickness (*29*) | Biodiversity-Ecosystem Function |
| Soil pH x 10 in H2O at 0.15m | (*23*) | pH x 10 | https://www.soilgrids.org ; https://www.isric.org/explore/wosis/accessin | (*23*) used data on 150,000 soil profiles worldwide to create a global map, integrating observations within profile to obtain datapoints at fixed depth.A machine learning approach was used, with input data on: Digital Elevation Model-derived characteristics (using the merged DEMS SRTMGL3 and GMTED, as described in (*24*)), long-term average and sd of MODIS- | Biodiversity-Ecosystem Function |

| | | | | | |
|---|---|---|---|---|---|
| | | | g-wosis-derived-datasets | based EVI, bands 4 and 7, long-term monthly average and standard deviation in land surface temperature (MODIS LST), long-term averaged mean monthly hours of snow cover (MODIS 8-day snow occurrence), land cover (using GlobCover30, described in (*25*), who classified Landsat data from 2000 and 2010), precipitation data (averaging WorldClim and GPCP), lithology (using GLiM, (*26*)), Global water table depth (*27*), long-term averaged mean monthly MODIS Flood Water, Landsat-based distribution of mangroves (*28*) and average soil and sedimentary-deposit thickness (*29*) | |
| Soil pH x 10 in H2O at 0.30m | (*23*) | pH x 10 | https://www.soilgrids.org ; https://www.isric.org/explore/wosis/accessing-wosis-derived-datasets | (*23*) used data on 150,000 soil profiles worldwide to create a global map, integrating observations within profile to obtain datapoints at fixed depth.A machine learning approach was used, with input data on: Digital Elevation Model-derived characteristics (using the merged DEMS SRTMGL3 and GMTED, as described in (*24*)), long-term average and sd of MODIS-based EVI, bands 4 and 7, long-term monthly average and standard deviation in land surface temperature (MODIS LST), long-term averaged mean monthly hours of snow cover (MODIS 8-day snow occurrence), land cover (using GlobCover30, described in (*25*), who classified Landsat data from 2000 and 2010), precipitation data (averaging WorldClim and GPCP), lithology (using GLiM, (*26*)), Global water table depth (*27*), long-term averaged mean monthly MODIS Flood Water, Landsat-based distribution of mangroves (*28*) and average soil and sedimentary-deposit thickness (*29*) | Biodiversity-Ecosystem Function |
| Soil pH x 10 in H2O at 0.60m | (*23*) | pH x 10 | https://www.soilgrids.org ; https://www.isric.org/explore/wosis/accessing-wosis-derived-datasets | (*23*) used data on 150,000 soil profiles worldwide to create a global map, integrating observations within profile to obtain datapoints at fixed depth.A machine learning approach was used, with input data on: Digital Elevation Model-derived characteristics (using the merged DEMS SRTMGL3 and GMTED, as described in (*24*)), long-term average and sd of MODIS-based EVI, bands 4 and 7, long-term monthly average and standard deviation in land surface temperature (MODIS LST), long-term averaged mean monthly hours of snow cover (MODIS 8-day snow occurrence), land cover (using GlobCover30, described in (*25*), who classified Landsat data from 2000 and 2010), precipitation data (averaging WorldClim and GPCP), lithology (using GLiM, (*26*)), Global water table depth (*27*), long-term averaged mean monthly MODIS Flood Water, Landsat-based | Biodiversity-Ecosystem Function |

| | | | | | |
|---|---|---|---|---|---|
| | | | | distribution of mangroves (*28*) and average soil and sedimentary-deposit thickness (*29*) | |
| Soil pH x 10 in H2O at 1.00m | (*23*) | pH x 10 | https://www.soilgrids.org ; https://www.isric.org/explore/wosis/accessing-wosis-derived-datasets | (*23*) used data on 150,000 soil profiles worldwide to create a global map, integrating observations within profile to obtain datapoints at fixed depth.A machine learning approach was used, with input data on: Digital Elevation Model-derived characteristics (using the merged DEMS SRTMGL3 and GMTED, as described in (*24*)), long-term average and sd of MODIS-based EVI, bands 4 and 7, long-term monthly average and standard deviation in land surface temperature (MODIS LST), long-term averaged mean monthly hours of snow cover (MODIS 8-day snow occurrence), land cover (using GlobCover30, described in (*25*), who classified Landsat data from 2000 and 2010), precipitation data (averaging WorldClim and GPCP), lithology (using GLiM, (*26*)), Global water table depth (*27*), long-term averaged mean monthly MODIS Flood Water, Landsat-based distribution of mangroves (*28*) and average soil and sedimentary-deposit thickness (*29*) | Biodiversity-Ecosystem Function |
| Soil pH x 10 in H2O at 2.00m | (*23*) | pH x 10 | https://www.soilgrids.org ; https://www.isric.org/explore/wosis/accessing-wosis-derived-datasets | (*23*) used data on 150,000 soil profiles worldwide to create a global map, integrating observations within profile to obtain datapoints at fixed depth.A machine learning approach was used, with input data on: Digital Elevation Model-derived characteristics (using the merged DEMS SRTMGL3 and GMTED, as described in (*24*)), long-term average and sd of MODIS-based EVI, bands 4 and 7, long-term monthly average and standard deviation in land surface temperature (MODIS LST), long-term averaged mean monthly hours of snow cover (MODIS 8-day snow occurrence), land cover (using GlobCover30, described in (*25*), who classified Landsat data from 2000 and 2010), precipitation data (averaging WorldClim and GPCP), lithology (using GLiM, (*26*)), Global water table depth (*27*), long-term averaged mean monthly MODIS Flood Water, Landsat-based distribution of mangroves (*28*) and average soil and sedimentary-deposit thickness (*29*) | Biodiversity-Ecosystem Function |
| Soil pH x 10 in KCL at 0.00m | (*23*) | pH x 10 | https://www.soilgrids.org ; https://www.isric.org/explore/wosis/accessin | (*23*) used data on 150,000 soil profiles worldwide to create a global map, integrating observations within profile to obtain datapoints at fixed depth.A machine learning approach was used, with input data on: Digital Elevation Model-derived characteristics (using the merged DEMS SRTMGL3 and GMTED, as described in (*24*)), long-term average and sd of MODIS- | Biodiversity-Ecosystem Function |

| | | | | | |
|---|---|---|---|---|---|
| | | | g-wosis-derived-datasets | based EVI, bands 4 and 7, long-term monthly average and standard deviation in land surface temperature (MODIS LST), long-term averaged mean monthly hours of snow cover (MODIS 8-day snow occurrence), land cover (using GlobCover30, described in (*25*), who classified Landsat data from 2000 and 2010), precipitation data (averaging WorldClim and GPCP), lithology (using GLiM, (*26*)), Global water table depth (*27*), long-term averaged mean monthly MODIS Flood Water, Landsat-based distribution of mangroves (*28*) and average soil and sedimentary-deposit thickness (*29*) | |
| Soil pH x 10 in KCL at 0.05m | (*23*) | pH x 10 | https://www.soilgrids.org ; https://www.isric.org/explore/wosis/accessing-wosis-derived-datasets | (*23*) used data on 150,000 soil profiles worldwide to create a global map, integrating observations within profile to obtain datapoints at fixed depth.A machine learning approach was used, with input data on: Digital Elevation Model-derived characteristics (using the merged DEMS SRTMGL3 and GMTED, as described in (*24*)), long-term average and sd of MODIS-based EVI, bands 4 and 7, long-term monthly average and standard deviation in land surface temperature (MODIS LST), long-term averaged mean monthly hours of snow cover (MODIS 8-day snow occurrence), land cover (using GlobCover30, described in (*25*), who classified Landsat data from 2000 and 2010), precipitation data (averaging WorldClim and GPCP), lithology (using GLiM, (*26*)), Global water table depth (*27*), long-term averaged mean monthly MODIS Flood Water, Landsat-based distribution of mangroves (*28*) and average soil and sedimentary-deposit thickness (*29*) | Biodiversity-Ecosystem Function |
| Soil pH x 10 in KCL at 0.15m | (*23*) | pH x 10 | https://www.soilgrids.org ; https://www.isric.org/explore/wosis/accessing-wosis-derived-datasets | (*23*) used data on 150,000 soil profiles worldwide to create a global map, integrating observations within profile to obtain datapoints at fixed depth.A machine learning approach was used, with input data on: Digital Elevation Model-derived characteristics (using the merged DEMS SRTMGL3 and GMTED, as described in (*24*)), long-term average and sd of MODIS-based EVI, bands 4 and 7, long-term monthly average and standard deviation in land surface temperature (MODIS LST), long-term averaged mean monthly hours of snow cover (MODIS 8-day snow occurrence), land cover (using GlobCover30, described in (*25*), who classified Landsat data from 2000 and 2010), precipitation data (averaging WorldClim and GPCP), lithology (using GLiM, (*26*)), Global water table depth (*27*), long-term averaged mean monthly MODIS Flood Water, Landsat-based | Biodiversity-Ecosystem Function |

| | | | | | |
|---|---|---|---|---|---|
| | | | | distribution of mangroves (*28*) and average soil and sedimentary-deposit thickness (*29*) | |
| Soil pH x 10 in KCL at 0.30m | (*23*) | pH x 10 | https://www.soilgrids.org ; https://www.isric.org/explore/wosis/accessing-wosis-derived-datasets | (*23*) used data on 150,000 soil profiles worldwide to create a global map, integrating observations within profile to obtain datapoints at fixed depth.A machine learning approach was used, with input data on: Digital Elevation Model-derived characteristics (using the merged DEMS SRTMGL3 and GMTED, as described in (*24*)), long-term average and sd of MODIS-based EVI, bands 4 and 7, long-term monthly average and standard deviation in land surface temperature (MODIS LST), long-term averaged mean monthly hours of snow cover (MODIS 8-day snow occurrence), land cover (using GlobCover30, described in (*25*), who classified Landsat data from 2000 and 2010), precipitation data (averaging WorldClim and GPCP), lithology (using GLiM, (*26*)), Global water table depth (*27*), long-term averaged mean monthly MODIS Flood Water, Landsat-based distribution of mangroves (*28*) and average soil and sedimentary-deposit thickness (*29*) | Biodiversity-Ecosystem Function |
| Soil pH x 10 in KCL at 0.60m | (*23*) | pH x 10 | https://www.soilgrids.org ; https://www.isric.org/explore/wosis/accessing-wosis-derived-datasets | (*23*) used data on 150,000 soil profiles worldwide to create a global map, integrating observations within profile to obtain datapoints at fixed depth.A machine learning approach was used, with input data on: Digital Elevation Model-derived characteristics (using the merged DEMS SRTMGL3 and GMTED, as described in (*24*)), long-term average and sd of MODIS-based EVI, bands 4 and 7, long-term monthly average and standard deviation in land surface temperature (MODIS LST), long-term averaged mean monthly hours of snow cover (MODIS 8-day snow occurrence), land cover (using GlobCover30, described in (*25*), who classified Landsat data from 2000 and 2010), precipitation data (averaging WorldClim and GPCP), lithology (using GLiM, (*26*)), Global water table depth (*27*), long-term averaged mean monthly MODIS Flood Water, Landsat-based distribution of mangroves (*28*) and average soil and sedimentary-deposit thickness (*29*) | Biodiversity-Ecosystem Function |
| Soil pH x 10 in KCL at 1.00m | (*23*) | pH x 10 | https://www.soilgrids.org ; https://www.isric.org/explore/wosis/accessin | (*23*) used data on 150,000 soil profiles worldwide to create a global map, integrating observations within profile to obtain datapoints at fixed depth.A machine learning approach was used, with input data on: Digital Elevation Model-derived characteristics (using the merged DEMS SRTMGL3 and GMTED, as described in (*24*)), long-term average and sd of MODIS- | Biodiversity-Ecosystem Function |

| | | | | | |
|---|---|---|---|---|---|
| | | | g-wosis-derived-datasets | based EVI, bands 4 and 7, long-term monthly average and standard deviation in land surface temperature (MODIS LST), long-term averaged mean monthly hours of snow cover (MODIS 8-day snow occurrence), land cover (using GlobCover30, described in (*25*), who classified Landsat data from 2000 and 2010), precipitation data (averaging WorldClim and GPCP), lithology (using GLiM, (*26*)), Global water table depth (*27*), long-term averaged mean monthly MODIS Flood Water, Landsat-based distribution of mangroves (*28*) and average soil and sedimentary-deposit thickness (*29*) | |
| Soil pH x 10 in KCL at 2.00m | (*23*) | pH x 10 | https://www.soilgrids.org ; https://www.isric.org/explore/wosis/accessing-wosis-derived-datasets | (*23*) used data on 150,000 soil profiles worldwide to create a global map, integrating observations within profile to obtain datapoints at fixed depth.A machine learning approach was used, with input data on: Digital Elevation Model-derived characteristics (using the merged DEMS SRTMGL3 and GMTED, as described in (*24*)), long-term average and sd of MODIS-based EVI, bands 4 and 7, long-term monthly average and standard deviation in land surface temperature (MODIS LST), long-term averaged mean monthly hours of snow cover (MODIS 8-day snow occurrence), land cover (using GlobCover30, described in (*25*), who classified Landsat data from 2000 and 2010), precipitation data (averaging WorldClim and GPCP), lithology (using GLiM, (*26*)), Global water table depth (*27*), long-term averaged mean monthly MODIS Flood Water, Landsat-based distribution of mangroves (*28*) and average soil and sedimentary-deposit thickness (*29*) | Biodiversity-Ecosystem Function |
| Aboveground biomass density (carbon fraction) | (*37*) | MgC/ha | https://daac.ornl.gov/cgi-bin/dsviewer.pl?ds_id=1763 | (*37*) combined multiple biomass maps (of woody vegetation, (*33*), (*38*)) and using inverse techniques to derive biomass estimates from AVHRR NDVI for grasslands (*39*) and Landsat ETM for tundra (*40*). In addition land use cover data (*41*, *42*), temperature data (Worldclim) and an update of the Koeppen biome classification were used as well. | Biodiversity-Ecosystem Function |
| Belowground biomass density (carbon fraction) | (*37*) | MgC/ha | https://daac.ornl.gov/cgi-bin/dsviewer.pl?ds_id=1763 | (*37*) combined multiple biomass maps (of woody vegetation, (*33*), (*38*)) and using inverse techniques to derive biomass estimates from AVHRR NDVI for grasslands (*39*) and Landsat ETM for tundra (*40*). In addition land use cover data (*41*, *42*), temperature data (Worldclim) and an update of the Koeppen biome classification were used as well. Previous | Biodiversity-Ecosystem Function |

| | | | | | |
|---|---|---|---|---|---|
| | | | | approaches were followed to relate (satellite-derived) aboveground carbon estimates to belowground carbon estimates (*43–45*) | |
| Above and belowground biomass density (carbon fraction) | (*46*) | MgC/ha | https://data-gis.unep-wcmc.org/portal/home/item.html?id=8a8d4e24683a46e6b039aea78c8af20f | (*46*) combined the global aboveground biomass maps of (*33*, *37–39*), and then included belowground biomass using root to shoot ratios suggested in the IPCC 2006 guidelines for national greenhouse gas inventories. | Biodiversity-Ecosystem Function |
| Mean Annual Decomposition Humus#1 | (*21*) | 10^-3 year^-1 | https://zenodo.org/record/4041038#.ZAh_2nYo8uU, https://www.worldclim.org/data/worldclim21.html | (*21*) used the Yasso07 model, which is a function of mean temperature and precipitation, the decomposition constants in this case probably changing for each type of litter | Biodiversity-Ecosystem Function |
| Mean Annual Decomposition Humus#2 | (*21*) | 10^-3 year^-1 | https://zenodo.org/record/4041038#.ZAh_2nYo8uU, https://www.worldclim.org/data/worldclim21.html | (*21*) used the Yasso07 model, which is a function of mean temperature and precipitation, the decomposition constants in this case probably changing for each type of litter | Biodiversity-Ecosystem Function |
| Mean Annual Decomposition Humus#3 | (*21*) | 10^-3 year^-1 | https://zenodo.org/record/4041038#.ZAh_2nYo8uU, https://www.worldclim.org/da | (*21*) used the Yasso07 model, which is a function of mean temperature and precipitation, the decomposition constants in this case probably changing for each type of litter | Biodiversity-Ecosystem Function |

| | | | | | |
|---|---|---|---|---|---|
| | | | ta/worldclim21.html | | |
| Alpha diversity, relative to pristine conditions | (*47*, *48*) | Unitless (proportion) | https://data.nhm.ac.uk/dataset/the-2016-release-of-the-predicts-database | Created using other variables in the dataset as input (see descriptions in (*47*, *49*)). In brief, abundance is calculated using linear mixed-effects models with Human Population Density (from 2005, GPW4 (same database as included in this study), Road density (different dataset groads v1, while GRIP is used here) used here, Land use types (*50*) and Land use intensity (using the Global Land Systems dataset ((*51*), as in (*52*, *53*)). Inputs are obtained via statistical downscaling to 30 arc minutes, which is then further downscaled to 15 arcminutes | Biodiversity-Ecosystem Function |
| Amphibian species richness | | No. of species per (0.5 by 0.5 degree) grid cell | https://sedac.ciesin.columbia.edu/data/set/species-global-amphibian-richness-2015 | Original data collected by the Global Amphibian Assessment (*54*)contributing to the IUCN's Red Listing programme | Biodiversity-Ecosystem Function |
| Species richness, relative to pristine conditions | (*47*, *48*) | Unitless (proportion) | https://data.nhm.ac.uk/dataset/global-maps-of-biodiversity-intactness-index-sanchez-ortiz-et-al-2019-biorxiv | Dataset is an update by (*55*) improving modelling methods and specific consideration of island dynamics as compared to (*52*). Same framework as for alpha diversity described above; response of biodiversity to land use, land use intensity, population density and road density is modelled. | Biodiversity-Ecosystem Function |
| Tree cover | | % cover | https://data.globalforestwatch.org/datasets/tree-cover-2000 | Tree cover based on spectral data from the Landsat 7, thematic mapper plus (ETM+) sensor, using minimum growing season NDVI to indicate cover (*34*). | Biodiversity-Ecosystem Function |
| Mammal species richness | | No. of species per (0.5 | https://sedac.ciesin.columbia.edu/data/set/s | Original data from the IUCN Red List. | Biodiversity-Ecosystem Function |

| | | | | | |
|---|---|---|---|---|---|
| | | by 0.5 degree) grid cell | pecies-global-mammal-richness-2015 | | |
| Global AI (Aridity Index) | | AI Value | https://cgiarcsi.community/2019/01/24/global-aridity-index-and-potential-evapotranspiration-climate-database-v2/ | No data from Antarctica; original values have been multiplied by 10,000 (i.e., multiply by 0.0001 to retrieve original values) | Global Change |
| Global Potential Evapotranspiration (PET) | | PET Value (mm) | https://cgiarcsi.community/2019/01/24/global-aridity-index-and-potential-evapotranspiration-climate-database-v2/ | No data from Antarctica | Global Change |
| Human Modification Index | (*56*) | Modification index ranging between 0 and 1 | https://developers.google.com/earth-engine/datasets/catalog/CSP_HM_GlobalHumanModification | Described in (*56*); Modification index based on human population density (GPW4, as used in this study), settlement density (Global Human settlements, (*57*)), cropland (Unified cropland database, (*58*)), livestock density (Gridded livestock of the world database, (*59*)), road and railroad density (Open Street Map, gRoads v1 and Digital Chart of the World (*60*)), mining and energy production (Open Street Map) and electrical infrastructure (based on Open Street Map data and nighttime lights database (DMSP-OLS, as used in this study), | Global Change |
| Water table depth (below surface) | (*27*) | m | https://aquaknow.jrc.ec.europa.eu/content/global-patterns-groundwater- | As described in (*27*), world-wide observations of water table depth are used to calibrate and validate groundwater models, yielding a global, high-resolution dataset of water table depths. | Global Change |

| | | | | | |
|---|---|---|---|---|---|
| | | | table-depth-wtd | | |
| Proportion of area covered by regrowth forest | | Unitless (proportion) | https://doi.pangaea.de/10.1594/PANGAEA.889943 | Four Plant Functional Types from MODIS Collection 5.1 land cover dataset are used, crosswalking land cover types to PFT fractions. Age distributions are based on country-level forest inventories for temperate and high latitude regions, and based on biomass for tropical regions (see Poulter et al. 2016; Pugh et al. 2016) | Global Change |
| Average age of regrowth forest stands | | year | https://doi.pangaea.de/10.1594/PANGAEA.889943 | Four Plant Functional Types from MODIS Collection 5.1 land cover dataset are used, crosswalking land cover types to PFT fractions. Age distributions are based on country-level forest inventories for temperate and high latitude regions, and based on biomass for tropical regions. | Global Change |
| Standard deviation of the age of regrowth forest stands | | year | https://doi.pangaea.de/10.1594/PANGAEA.889943 | Four Plant Functional Types from MODIS Collection 5.1 land cover dataset are used, crosswalking land cover types to PFT fractions. Age distributions are based on country-level forest inventories for temperate and high latitude regions, and based on biomass for tropical regions. | Global Change |
| Fraction of incident photosynthetically active radiation (400-700nm) absorbed by the green elements of a vegetation canopy (fPAR) | | fPAR fraction | https://explorer.earthengine.google.com/#detail/MODIS%2F006%2FMCD15A3H | Masks substantial quantities of arid areas; averaged from the calendar years 2015-2019 | Global Change |
| Human footprint index | (*4*) | ranging between 0 and 50 | https://datadryad.org/stash/dataset/doi:10.5061/dryad.052q5 | An integrated index quantifying human pressure. Described in (*4*); pressure included settlement density (DMSP-OLS, as used in this study), human population density (GPW4, as used in this study), electric infrastructure (DMSP-OLS, as used in this study), agriculture (Globcover v.2.3, (*61*)), pasture lands (*41*), road density (gRoadsv1, https://sedac.ciesin.columbia.edu/data/set/groads-global-roads-open-access-v1) and railway density (*62*) | Global Change |

| | | | | | |
|---|---|---|---|---|---|
| Annual rate of decrease in forest cover between 2000 and 2019 | (*34*) | % cover | https://earthenginepartners.appspot.com/science-2013-global-forest/download_v1.7.html | Tree cover based on spectral data from the Landsat 7, thematic mapper plus (ETM+) sensor, using minimum growing season NDVI to indicate cover (*34*). Version 1.7 runs until 2019 | Global Change |
| Total increase in forest cover between 2000 and 2019 | (*34*) | % cover | https://earthenginepartners.appspot.com/science-2013-global-forest/download_v1.7.html | Tree cover based on spectral data from the Landsat 7, thematic mapper plus (ETM+) sensor, using minimum growing season NDVI to indicate cover (*34*). Version 1.7 runs until 2019 | Global Change |
| Proportion of impervious surface area within a grid cell | (*63*) | Unitless (proportion) | http://www.riverthreat.net/data.html | (*63*) upscaled the product of (*64*)(1km2), which modelled impervious surface as dependent variable in a regression equation including nighttime lights (DMSP-OLS, as used in this study), and human population density (LandScan 2004: https://geodata.lib.utexas.edu/catalog/nyu-2451-43669, which in turn uses MODIS land cover data (*65*), topographic data from the SRTM mission (*66*) and the NGA's Controlled Image Base: https://earth-info.nga.mil/index.php?dir=pi&action=pi) | Global Change |
| Proportion of wetlands occupied by croplands or urban areas | (*63*) | Unitless (proportion) | http://www.riverthreat.net/data.html | (*63*) intersected the wetland area of the Global Lakes and Wetlands database (*67*) | Global Change |
| Nitrogen loading index | (*63*) | Between 0 and 1 | http://www.riverthreat.net/data.html | Anthropogenic nitrogen loads to rivers and their catchments (nitrogen per area per time per unit discharge). (*63*) used the global nitrogen loading dataset of (*68*), who used results from atmospheric transport models, and FAO data on cropland areas, fertilizer production and waste treatment to model nitrogen loads | Global Change |

| | | | | | |
|---|---|---|---|---|---|
| Phosphorus loading index | (*63*) | Between 0 and 1 | http://www.riverthreat.net/data.html | Anthropogenic (and partly natural) phosphorus loads to rivers and their catchments (phosphorus per area per time per unit discharge). (*63*) combined data from point- and non-point pollution sources into rivers by (*69*), who in turn used the spatially explicit model HYDE using population density (*70*, *71*), fertilizer input and sewage output data from (*72*), and country GNP (world bank)) and data on anthropogenic atmospheric phosphorous deposition by (*73*), who used an atmospheric circulation model calibrated on observational data of phosphorous-containing aerosols, and data sources on fossil fuel combustion and other sources of aerosol production. | Global Change |
| Mercury loading index | (*63*) | Between 0 and 1 | http://www.riverthreat.net/data.html | Anthropogenic mercury loads to rivers and their catchments (mercury per area per time per unit discharge). (*63*) used the data from current and pre-industrial mercury deposition rates from (*74*), who in turn modelled global mercury dynamics using a 3D global circulation model, using input data on mercury sources including mines, volcanoes and biomass burning (assuming a fixed ratio between CO and Hg) | Global Change |
| Pesticide loading index | (*63*) | Between 0 and 1 | http://www.riverthreat.net/data.html | Pesticide loads to rivers and their catchments (pesticides per area per time per unit discharge). (*63*) combined country level data on pesticide use (Environmental Sustainability Index), with a land cover map specifying agricultural lands (*41*) | Global Change |
| Organic loading index | (*63*) | Between 0 and 1 | http://www.riverthreat.net/data.html | Biological oxygen demand of rivers due to addition of labile organic carbon from sewage. (*63*) used fixed ratios between BOD and Nitrogen loading, depending on sewage treatment level, which was taken from the GRUMP dataset: https://sedac.ciesin.columbia.edu/data/collection/grump-v1 | Global Change |
| Potential Acidification index | (*63*) | Between 0 and 1 | http://www.riverthreat.net/data.html | Loads of acidifying agents to rivers and their catchments (H+ equivalents per area per time per unit discharge). (*63*) used data on SOx and NOx emissions from (*75*), based on ensemble analysis of 23 atmospheric chemistry transport models), and determined susceptibility to acidification using pH data from SoilData database (v0; https://www.isric.org/explore/soilinfo) | Global Change |

| | | | | | |
|---|---|---|---|---|---|
| Dam density | (*63*) | Number of dams km-2 | http://www.riverthreat.net/data.html | Number of medium-sized, large and very large dams per area of the catchment. (*63*) used data on very large dams from the GWSP-GRAND database, smaller dam numbers taken from the ICOLD database, and distributed over each country, using a regression equation based on population density, discharge, cropland area, elevation and a cropland*elevation interaction term | Global Change |
| Water stress index. | (*63*) | Between 0 and 1 | http://www.riverthreat.net/data.html | Stress is quantified by (one minus) the (standardized) ratio between available water (river discharge) and human population density. (*63*) used population density data from the CIESIN-GRUMP database, and discharge data from the global hydrological modelling outputs of (*76*) and (*77*) | Global Change |
| Agricultural stress index. | (*63*) | Between 0 and 1 | http://www.riverthreat.net/data.html | Stress is quantified by (one minus) the (standardized) ratio between available water (river discharge) and cropland area. (*63*) used cropland area from (*41*) and discharge data from the global hydrological modelling outputs of (*76*) and (*77*) | Global Change |
| Fishing pressure stress index | (*63*) | Between 0 and 1 | http://www.riverthreat.net/data.html | Fishing pressure is calculated as the ratio between estimated fish catches and estimated fish production within the grid cell. (*63*) used country-level catch data of inland and diadromous fish (FAO FishStat Plus, 1997-2006), and distributed these catches over space using an empirical power function scaling-relationship with river discharge (and discharge data from the global hydrological modelling outputs of (*76*) and (*77*)). Potential production was quantified as the log(NPP), using outputs from the Global Ecosystem Model IBIS (*78*) | Global Change |
| Aquaculture pressure index | (*63*) | Between 0 and 1 | http://www.riverthreat.net/data.html | Aquaculture pressure quantified as the amount of fish produced by aquaculture within the grid cell. (*63*) used country-level aquaculture data of inland and diadromous fish (FAO FishStat Plus, 1997-2006), and distributed these catches over space using an empirical power function scaling-relationship with river discharge (from the global hydrological modelling outputs of (*76*) and (*77*)). | Global Change |
| Biodiversity threat index | (*63*) | Between 0 and 1 | http://www.riverthreat.net/data.html | Aggregate index summing the combined impact of 23 drivers on river-associated biodiversity. (*63*) summed the impacts of 23 drivers, with equal | Global Change |

| | | | | | |
|---|---|---|---|---|---|
| | | | | weighing of four themes, and expert-based weighing of drivers within themes | |
| Human Water Security threat index | (*63*) | Between 0 and 1 | http://www.riverthreat.net/data.html | Aggregate index summing the combined impact of 23 drivers on human water security (HWS). (*63*) summed the impacts of 23 drivers, with equal weighing of four themes, and expert-based weighing of drivers within themes | Global Change |
| Adjusted HWS threat index | (*63*) | Between 0 and 1 | http://www.riverthreat.net/data.html | Aggregate index summing the combined impact of 23 drivers on human water security (HWS), adjusted for security measures. (*63*) adjusted the incident water threat using an investment benefits factor (both these variables included in the dataset) | Global Change |
| Absolute change in annual mean temperature, comparing 1957-1986 to 1987-2016 | (*79*) | °C/10 | https://envicloud.wsl.ch/#/?prefix=chelsa%2Fchelsa_V1%2Fchelsa_cruts | (*79*) used ERA-interim (6-hour synoptic) reanalysis data to calculate minimum and maximum monthly temperature, the mean is then calculated as the mean of minimum and maximum temperature. | Global Change |
| Absolute change in the mean diurnal temperature range, comparing 1957-1986 to 1987-2016 | (*79*) | °C/10 | https://envicloud.wsl.ch/#/?prefix=chelsa%2Fchelsa_V1%2Fchelsa_cruts | (*79*) used ERA-interim (6-hour synoptic) reanalysis data to calculate minimum and maximum monthly temperature. Mean diurnal range is calculated as the mean of the monthly range (max. temp. minus min. temp) | Global Change |
| Absolute change in isothermality (mean diurnal:annual temperature range), comparing | (*79*) | °C/10 | https://envicloud.wsl.ch/#/?prefix=chelsa%2Fchelsa_V1%2Fchelsa_cruts | (*79*) used ERA-interim (6-hour synoptic) reanalysis data to calculate minimum and maximum monthly temperature. Isothermality is calculated as the ratio of BIOCLIM2 and BIOCLIM7 | Global Change |

| | | | | | |
|---|---|---|---|---|---|
| 1957-1986 to 1987-2016 | | | | | |
| Absolute change in temperature seasonality, comparing 1957-1986 to 1987-2016 | (*79*) | °C/10 | https://envicloud.wsl.ch/#/?prefix=chelsa%2Fchelsa_V1%2Fchelsa_cruts | (*79*) used ERA-interim (6-hour synoptic) reanalysis data to calculate minimum and maximum monthly temperature. Seasonality is calculated as the standard deviation * 100 | Global Change |
| Absolute change in the max. temperature of the warmest month, comparing 1957-1986 to 1987-2016 | (*79*) | °C/10 | https://envicloud.wsl.ch/#/?prefix=chelsa%2Fchelsa_V1%2Fchelsa_cruts | (*79*) used ERA-interim (6-hour synoptic) reanalysis data to calculate minimum and maximum monthly temperature. | Global Change |
| Absolute change in the min. temperature of the coldest month, comparing 1957-1986 to 1987-2016 | (*79*) | °C/10 | https://envicloud.wsl.ch/#/?prefix=chelsa%2Fchelsa_V1%2Fchelsa_cruts | (*79*) used ERA-interim (6-hour synoptic) reanalysis data to calculate minimum and maximum monthly temperature. | Global Change |
| Absolute change in temperature annual range, comparing | (*79*) | °C/10 | https://envicloud.wsl.ch/#/?prefix=chelsa%2Fchelsa_V1%2Fchelsa_cruts | (*79*) used ERA-interim (6-hour synoptic) reanalysis data to calculate minimum and maximum monthly temperature. Seasonality is calculated as the standard deviation * 100 | Global Change |

| | | | | | |
|---|---|---|---|---|---|
| 1957-1986 to 1987-2016 | | | | | |
| Absolute change in mean temperature of the wettest quarter, comparing 1957-1986 to 1987-2016 | (*79*) | °C/10 | https://envicloud.wsl.ch/#/?prefix=chelsa%2Fchelsa_V1%2Fchelsa_cruts | (*79*) used ERA-interim (6-hour synoptic) reanalysis data to calculate minimum and maximum monthly temperature. Temperature annual range is calculated as the difference between BIOCLIM5 and BIOCLIM6 | Global Change |
| Absolute change in mean temperature of the driest quarter, comparing 1957-1986 to 1987-2016 | (*79*) | °C/10 | https://envicloud.wsl.ch/#/?prefix=chelsa%2Fchelsa_V1%2Fchelsa_cruts | (*79*) used ERA-interim (6-hour synoptic) reanalysis data to calculate minimum and maximum monthly temperature. Seasonality is calculated as the standard deviation * 100 | Global Change |
| Absolute change in mean temperature of the warmest quarter, comparing 1957-1986 to 1987-2016 | (*79*) | °C/10 | https://envicloud.wsl.ch/#/?prefix=chelsa%2Fchelsa_V1%2Fchelsa_cruts | (*79*) used ERA-interim (6-hour synoptic) reanalysis data to calculate minimum and maximum monthly temperature, and precipitation. | Global Change |
| Absolute change in mean temperature of the coldest quarter, comparing | (*79*) | °C/10 | https://envicloud.wsl.ch/#/?prefix=chelsa%2Fchelsa_V1%2Fchelsa_cruts | (*79*) used ERA-interim (6-hour synoptic) reanalysis data to calculate minimum and maximum monthly temperature, and precipitation. | Global Change |

| | | | | | |
|---|---|---|---|---|---|
| 1957-1986 to 1987-2016 | | | | | |
| Absolute change in annual precipitation, comparing 1957-1986 to 1987-2016 | (*79*) | kg/m2 | https://envicloud.wsl.ch/#/?prefix=chelsa%2Fchelsa_V1%2Fchelsa_cruts | (*79*) used ERA-interim (6-hour synoptic) reanalysis data to calculate minimum and maximum monthly temperature, and precipitation. | Global Change |
| Absolute change in precipitation of the wettest month, comparing 1957-1986 to 1987-2016 | (*79*) | kg/m2 | https://envicloud.wsl.ch/#/?prefix=chelsa%2Fchelsa_V1%2Fchelsa_cruts | (*79*) used ERA-interim (6-hour synoptic) reanalysis data to calculate minimum and maximum monthly temperature, and precipitation. | Global Change |
| Absolute change in precipitation of the driest month, comparing 1957-1986 to 1987-2016 | (*79*) | kg/m2 | https://envicloud.wsl.ch/#/?prefix=chelsa%2Fchelsa_V1%2Fchelsa_cruts | (*79*) used ERA-interim (6-hour synoptic) reanalysis data to calculate minimum and maximum monthly temperature, and precipitation. | Global Change |
| Absolute change in precipitation seasonality comparing 1957-1986 to 1987-2016 | (*79*) | kg/m2 | https://envicloud.wsl.ch/#/?prefix=chelsa%2Fchelsa_V1%2Fchelsa_cruts | (*79*) used ERA-interim (6-hour synoptic) reanalysis data to calculate minimum and maximum monthly temperature, and precipitation. | Global Change |

| | | | | | |
|---|---|---|---|---|---|
| Absolute change in precipitation of the wettest quarter, comparing 1957-1986 to 1987-2016 | (*79*) | kg/m2 | https://envicloud.wsl.ch/#/?prefix=chelsa%2Fchelsa_V1%2Fchelsa_cruts | (*79*) used ERA-interim (6-hour synoptic) reanalysis data to calculate minimum and maximum monthly temperature, and precipitation. | Global Change |
| Absolute change in precipitation of the driest quarter, comparing 1957-1986 to 1987-2016 | (*79*) | kg/m2 | https://envicloud.wsl.ch/#/?prefix=chelsa%2Fchelsa_V1%2Fchelsa_cruts | (*79*) used ERA-interim (6-hour synoptic) reanalysis data to calculate minimum and maximum monthly temperature, and precipitation. | Global Change |
| Absolute change in precipitation of the warmest quarter, comparing 1957-1986 to 1987-2016 | (*79*) | kg/m2 | https://envicloud.wsl.ch/#/?prefix=chelsa%2Fchelsa_V1%2Fchelsa_cruts | (*79*) used ERA-interim (6-hour synoptic) reanalysis data to calculate minimum and maximum monthly temperature, and precipitation. | Global Change |
| Absolute change in precipitation of the coldest quarter, comparing 1957-1986 to 1987-2016 | (*79*) | kg/m2 | https://envicloud.wsl.ch/#/?prefix=chelsa%2Fchelsa_V1%2Fchelsa_cruts | (*79*) used ERA-interim (6-hour synoptic) reanalysis data to calculate minimum and maximum monthly temperature, and precipitation. | Global Change |

| | | | | | |
|---|---|---|---|---|---|
| Percentage of the pixel area covered by cultivated and managed vegetation (i.e., agriculture) | (*18*) | percentage (0-100) | https://www.earthenv.org/landcover | (*18*) created a global 1-km scale consensus land cover product, integrating the DISCover (*19*), GLC2000 (*20*), MODIS2005 (*80*) and GlobCover (*81*) land cover products | Ecosystem Service |
| Available soil water capacity (volumetric fraction) until wilting point for depth 0 cm | (*23*) | % | https://www.soilgrids.org ; https://www.isric.org/explore/wosis/accessing-wosis-derived-datasets | (*23*) used ~ data on 150,000 soil profiles worldwide to create a global map, integrating observations within profile to obtain datapoints at fixed depth.A machine learning approach was used, with input data on: Digital Elevation Model-derived characteristics (using the merged DEMS SRTMGL3 and GMTED, as described in (*24*), long-term average and sd of MODIS-based EVI, bands 4 and 7, long-term monthly average and standard deviation in land surface temperature (MODIS LST), long-term averaged mean monthly hours of snow cover (MODIS 8-day snow occurrence), land cover (using GlobCover30, described in (*25*), who classified Landsat data from 2000 and 2010), precipitation data (averaging WorldClim and GPCP), lithology (using GLiM, (*26*)), Global water table depth (*27*), long-term averaged mean monthly MODIS Flood Water, Landsat-based distribution of mangroves (*28*) and average soil and sedimentary-deposit thickness (*29*) | Ecosystem Service |
| Available soil water capacity (volumetric fraction) until wilting point for depth 5 cm | (*23*) | % | https://www.soilgrids.org ; https://www.isric.org/explore/wosis/accessing-wosis-derived-datasets | (*23*) used ~ data on 150,000 soil profiles worldwide to create a global map, integrating observations within profile to obtain datapoints at fixed depth.A machine learning approach was used, with input data on: Digital Elevation Model-derived characteristics (using the merged DEMS SRTMGL3 and GMTED, as described in (*24*), long-term average and sd of MODIS-based EVI, bands 4 and 7, long-term monthly average and standard deviation in land surface temperature (MODIS LST), long-term averaged mean monthly hours of snow cover (MODIS 8-day snow occurrence), land cover (using GlobCover30, described in (*25*), who classified Landsat data from 2000 and 2010), precipitation data (averaging WorldClim and GPCP), lithology (using GLiM, (*26*)), Global water table depth (*27*), long-term averaged mean monthly MODIS Flood Water, Landsat-based distribution of mangroves (*28*) and average soil and sedimentary-deposit thickness (*29*) | Ecosystem Service |

| | | | | | |
|---|---|---|---|---|---|
| Available soil water capacity (volumetric fraction) until wilting point for depth 15 cm | (*23*) | % | https://www.soilgrids.org ; https://www.isric.org/explore/wosis/accessing-wosis-derived-datasets | (*23*) used ~ data on 150,000 soil profiles worldwide to create a global map, integrating observations within profile to obtain datapoints at fixed depth.A machine learning approach was used, with input data on: Digital Elevation Model-derived characteristics (using the merged DEMS SRTMGL3 and GMTED, as described in (*24*), long-term average and sd of MODIS-based EVI, bands 4 and 7, long-term monthly average and standard deviation in land surface temperature (MODIS LST), long-term averaged mean monthly hours of snow cover (MODIS 8-day snow occurrence), land cover (using GlobCover30, described in (*25*), who classified Landsat data from 2000 and 2010), precipitation data (averaging WorldClim and GPCP), lithology (using GLiM, (*26*)), Global water table depth (*27*), long-term averaged mean monthly MODIS Flood Water, Landsat-based distribution of mangroves (*28*) and average soil and sedimentary-deposit thickness (*29*) | Ecosystem Service |
| Available soil water capacity (volumetric fraction) until wilting point for depth 30 cm | (*23*) | % | https://www.soilgrids.org ; https://www.isric.org/explore/wosis/accessing-wosis-derived-datasets | (*23*) used ~ data on 150,000 soil profiles worldwide to create a global map, integrating observations within profile to obtain datapoints at fixed depth.A machine learning approach was used, with input data on: Digital Elevation Model-derived characteristics (using the merged DEMS SRTMGL3 and GMTED, as described in (*24*), long-term average and sd of MODIS-based EVI, bands 4 and 7, long-term monthly average and standard deviation in land surface temperature (MODIS LST), long-term averaged mean monthly hours of snow cover (MODIS 8-day snow occurrence), land cover (using GlobCover30, described in (*25*), who classified Landsat data from 2000 and 2010), precipitation data (averaging WorldClim and GPCP), lithology (using GLiM, (*26*)), Global water table depth (*27*), long-term averaged mean monthly MODIS Flood Water, Landsat-based distribution of mangroves (*28*) and average soil and sedimentary-deposit thickness (*29*) | Ecosystem Service |
| Available soil water capacity (volumetric fraction) until wilting point for depth 60 cm | (*23*) | % | https://www.soilgrids.org ; https://www.isric.org/explore/wosis/accessing-wosis- | (*23*) used ~ data on 150,000 soil profiles worldwide to create a global map, integrating observations within profile to obtain datapoints at fixed depth.A machine learning approach was used, with input data on: Digital Elevation Model-derived characteristics (using the merged DEMS SRTMGL3 and GMTED, as described in (*24*), long-term average and sd of MODIS-based EVI, bands 4 and 7, long-term monthly average and standard deviation in land surface temperature (MODIS LST), long-term averaged mean monthly hours of snow cover (MODIS 8-day snow | Ecosystem Service |

| | | | | | |
|---|---|---|---|---|---|
| | | | derived-datasets | occurrence), land cover (using GlobCover30, described in (*25*), who classified Landsat data from 2000 and 2010), precipitation data (averaging WorldClim and GPCP), lithology (using GLiM, (*26*)), Global water table depth (*27*), long-term averaged mean monthly MODIS Flood Water, Landsat-based distribution of mangroves (*28*) and average soil and sedimentary-deposit thickness (*29*) | |
| Available soil water capacity (volumetric fraction) until wilting point for depth 100 cm | (*23*) | % | https://www.soilgrids.org ; https://www.isric.org/explore/wosis/accessing-wosis-derived-datasets | (*23*) used ~ data on 150,000 soil profiles worldwide to create a global map, integrating observations within profile to obtain datapoints at fixed depth.A machine learning approach was used, with input data on: Digital Elevation Model-derived characteristics (using the merged DEMS SRTMGL3 and GMTED, as described in (*24*), long-term average and sd of MODIS-based EVI, bands 4 and 7, long-term monthly average and standard deviation in land surface temperature (MODIS LST), long-term averaged mean monthly hours of snow cover (MODIS 8-day snow occurrence), land cover (using GlobCover30, described in (*25*), who classified Landsat data from 2000 and 2010), precipitation data (averaging WorldClim and GPCP), lithology (using GLiM, (*26*)), Global water table depth (*27*), long-term averaged mean monthly MODIS Flood Water, Landsat-based distribution of mangroves (*28*) and average soil and sedimentary-deposit thickness (*29*) | Ecosystem Service |
| Available soil water capacity (volumetric fraction) until wilting point for depth 200 cm | (*23*) | % | https://www.soilgrids.org ; https://www.isric.org/explore/wosis/accessing-wosis-derived-datasets | (*23*) used ~ data on 150,000 soil profiles worldwide to create a global map, integrating observations within profile to obtain datapoints at fixed depth.A machine learning approach was used, with input data on: Digital Elevation Model-derived characteristics (using the merged DEMS SRTMGL3 and GMTED, as described in (*24*), long-term average and sd of MODIS-based EVI, bands 4 and 7, long-term monthly average and standard deviation in land surface temperature (MODIS LST), long-term averaged mean monthly hours of snow cover (MODIS 8-day snow occurrence), land cover (using GlobCover30, described in (*25*), who classified Landsat data from 2000 and 2010), precipitation data (averaging WorldClim and GPCP), lithology (using GLiM, (*26*)), Global water table depth (*27*), long-term averaged mean monthly MODIS Flood Water, Landsat-based distribution of mangroves (*28*) and average soil and sedimentary-deposit thickness (*29*) | Ecosystem Service |

| | | | | | |
|---|---|---|---|---|---|
| Area harvested for food crops | (*82*) | ha | https://dataverse.harvard.edu/dataset.xhtml?persistentId=doi:10.7910/DVN/PRFF8V | Output from the Spatial Production Allocation Model (SPAM, (*83*). In brief, SPAM uses country-level data on harvested area and yield (FAOSTAT 2015), allocating this production over a cropland area obtained from IIASA-IFPRI, who made a composite cropland area map using GlobCover 2005 and MODIS v5 as global sources, further supplemented with regional and national satellite imagery. | Ecosystem Service |
| Riverine sediment flux | | kg/s | https://sdml.ua.edu/datasets-2/ | Sediment flux is calculated with the WBMsed model of (*84*), which includes the empirical BQART model, which uses topographic/elevation data, discharge data, upstream catchment area, temperature, geological factors and human-induced erosion (the latter being assigned different values based on low or high population density and low/high GNP within the river catchment, (*85*)) | Ecosystem Service |
| Proportion of the total grid cell area consisting of cropland | (*63*) | Rescaled to a Cropland index ranging between 0 and 1 | http://www.riverthreat.net/data.html | (*63*) used cropland area from (*41*). | Ecosystem Service |
| Percentage of urban/built-up areas summed with cultivated/managed vegetation | (*18*) | percentage (0-100) | https://www.earthenv.org/landcover | Percentage of urban/built-up areas summed with cultivated/managed vegetation (summed via composite code). (*18*) created a global 1-km scale consensus land cover product, integrating the DISCover (*19*), GLC2000 (*20*), MODIS2005 (*80*) and GlobCover (*81*) land cover products | Social-Economic Landscape |
| Human population density | | No people/km2 | https://sedac.ciesin.columbia.edu/data/collection/gpw-v4 | Population estimates from the World Population Prospects: 2015 revision (UN, 2015: https://www.un.org/en/development/desa/publications/world-population-prospects-2015-revision.html) were spatially allocated within administrative boundaries (GADMv2), homogeneously distributing the population over each census area | Social-Economic Landscape |

| | | | | | |
|---|---|---|---|---|---|
| Density of all road types combined | (*86*) | m/km2 | https://www.globio.info/download-grip-dataset#:~:text=The%20GRIP%20dataset%20consists%20of | (*86*) harmonized and integrated nearly 60 geospatial datasets of road infrastructure | Social-Economic Landscape |
| Density of primary roads | (*86*) | m/km2 | https://www.globio.info/download-grip-dataset#:~:text=The%20GRIP%20dataset%20consists%20of | (*86*) harmonized and integrated nearly 60 geospatial datasets of road infrastructure | Social-Economic Landscape |
| Density of secondary roads | (*86*) | m/km2 | https://www.globio.info/download-grip-dataset#:~:text=The%20GRIP%20dataset%20consists%20of | (*86*) harmonized and integrated nearly 60 geospatial datasets of road infrastructure | Social-Economic Landscape |
| Density of tertiary roads | (*86*) | m/km2 | https://www.globio.info/download-grip-dataset#:~:text=The%20GRIP%20dataset%20consists%20of | (*86*) harmonized and integrated nearly 60 geospatial datasets of road infrastructure | Social-Economic Landscape |

| | | | | | |
|---|---|---|---|---|---|
| Density of local roads | (*86*) | m/km2 | https://www.globio.info/download-grip-dataset#:~:text=The%20GRIP%20dataset%20consists%20of | (*86*) harmonized and integrated nearly 60 geospatial datasets of road infrastructure | Social-Economic Landscape |
| Investment index | (*63*) | An investment benefit factor that ranges between 0 and 1 | http://www.riverthreat.net/data.html | Index that quantifies the investments made into improving water infrastructure and management. (*63*) developed an aggregate index using their drivers dam density, flow disruption, wetland connectivity, supplemented with a Moderate Water Use variable (calculated as the ratio between human water use and discharge (using the outputs from global hydrological model outputs (*76*, *77*)) and Access to Clean Drinking water - the proportion of the population having access to clean drinking water (WHO 2004: https://apps.who.int/iris/handle/10665/42891) | Social-Economic Landscape |

**Table S3.** Proportional details for the colored bars in **Fig. 3**.

| Causal graph | Variable label | positive | neutral | negative | sum |
|---|---|---|---|---|---|
| 1 | Human Development Index->Cropland | 0.44 | 0.55 | 0.02 | 1.01 |
| 1 | Human Development Index->Tree_Density | 0.02 | 0.07 | 0.90 | 0.99 |
| 1 | Human Development Index -> Human_Footprint_2009 | 0.45 | 0.55 | 0.00 | 1.00 |
| 1 | Cropland-> Human Development Index | 0.44 | 0.57 | 0.05 | 1.06 |
| 1 | Cropland->Tree_Density | 0.13 | 0.27 | 0.59 | 0.99 |
| 1 | Cropland->Human_Footprint_2009 | 0.46 | 0.54 | 0.01 | 1.00 |
| 1 | Tree_Density-> Human Development Index | 0.00 | 1.00 | 0.00 | 1.00 |
| 1 | Tree_Density->Cropland | 0.00 | 0.95 | 0.05 | 1.00 |
| 1 | Tree_Density-> Human_Footprint_2009 | 0.00 | 0.99 | 0.02 | 1.00 |
| 1 | Human_Footprint_2009-> Human Development Index | 0.55 | 0.45 | 0.00 | 1.00 |
| 1 | Human_Footprint_2009->Cropland | 0.53 | 0.47 | 0.00 | 1.00 |
| 1 | Human_Footprint_2009-> Tree_Density | 0.00 | 0.24 | 0.74 | 0.99 |
| 2 | Species richness->Food area | 0.00 | 1.00 | 0.00 | 1.00 |
| 2 | Species richness->Investment_benefit_factor | 0.12 | 0.58 | 0.33 | 1.04 |
| 2 | Species richness->Phosphorus_Loading | 0.18 | 0.43 | 0.40 | 1.00 |
| 2 | Food area->biodiversity | 0.00 | 0.27 | 0.77 | 1.04 |
| 2 | Food area->Investment_benefit_factor | 0.08 | 0.78 | 0.15 | 1.00 |
| 2 | Food area->Phosphorus_Loading | 0.61 | 0.42 | 0.02 | 1.04 |
| 2 | Investment_benefit_factor->Species richness | 0.17 | 0.43 | 0.41 | 1.02 |
| 2 | Investment_benefit_factor->Food area | 0.00 | 1.00 | 0.00 | 1.00 |
| 2 | Investment_benefit_factor->Phosphorus_Loading | 0.43 | 0.41 | 0.18 | 1.02 |
| 2 | Phosphorus_Loading->Species richness | 0.18 | 0.46 | 0.42 | 1.07 |
| 2 | Phosphorus_Loading->Food area | 0.00 | 1.00 | 0.00 | 1.00 |
| 2 | Phosphorus_Loading->Investment_benefit_factor | 0.40 | 0.57 | 0.13 | 1.09 |

| | | | | | |
|---|---|---|---|---|---|
| 3 | Alpha diversity->AnnualForestLoss2000to2019 | 0.00 | 1.00 | 0.01 | 1.00 |
| 3 | Alpha diversity-> Population_Density | 0.07 | 0.40 | 0.53 | 1.00 |
| 3 | Alpha diversity->Water Holding Capacity | 0.05 | 0.38 | 0.62 | 1.05 |
| 3 | AnnualForestLoss2000to2019->Alpha diversity | 0.00 | 0.38 | 0.61 | 1.00 |
| 3 | AnnualForestLoss2000to2019-> Population_Density | 0.00 | 0.54 | 0.53 | 1.07 |
| 3 | AnnualForestLoss2000to2019->Water Holding Capacity | 0.31 | 0.68 | 0.01 | 0.99 |
| 3 | Population_Density->Alpha diversity | 0.00 | 0.78 | 0.25 | 1.03 |
| 3 | Population_Density->AnnualForestLoss2000to2019 | 0.00 | 0.99 | 0.02 | 1.01 |
| 3 | Population_Density->Water Holding Capacity | 0.06 | 0.86 | 0.07 | 1.00 |
| 3 | Water Holding Capacity->Alpha diversity | 0.03 | 0.63 | 0.34 | 1.00 |
| 3 | Water Holding Capacity->AnnualForestLoss2000to2019 | 0.00 | 1.00 | 0.00 | 1.00 |
| 3 | Water Holding Capacity-> Population_Density | 0.11 | 0.53 | 0.36 | 1.00 |

**Table S4.** Causal graphs and frequency of occurrence and associations on the complete text corpus resulting from our literature review (47665 words/terms in total entries of 356772525):

Tree density ($r_{\text{'tree' \& 'densiti'}}$=0.18; 'tree' n=3569, 'densiti' n=1171); Human development ($r_{\text{'human'\&'develop'}}$<0.1, 'human' n=863, 'develop' n=1701; $r_{\text{'econom'\&'develop'}}$=0.23, 'econom' n=581); Human footprint ($r_{\text{'footprint'\&'water'}}$=0.15, 'footprint' n= 24; 'water' n=1271); Species richness ($r_{\text{'speci'\&''rich'}}$=0.30, 'speci' n= 11860, 'rich' n=918); Food area ($r_{\text{'food'\&'feed'}}$=0.19, 'food' n=682, 'feed' n=403); Investment benefit factor ($r_{\text{'invest'\&'econometr'}}$=0.23, 'invest' n=113, 'econometr' n=2; $r_{\text{'benefit'\&'valuat'}}$=0.29, 'benefit' n=342, 'valuat' n=11); Phosphorus loading ('phosphorus'=143); Alpha diversity ($r_{\text{'alpha'\&'diver'}}$=0.19, 'alpha' n=66, 'diver' n=2162); Soil water content ($r_{\text{'soil'\&'moistur'}}$=0.16; 'soil'=2502, 'moistur'=125); Population density ($r_{\text{'populationlevel'\&'densiti'}}$<0.10, 'populationlevel' n=6, 'densiti' n=1171); Forest loss ($r_{\text{'forest'\&'log'}}$<0.43, 'forest' n=10030, 'log' n=2208).

| | Causal graph relation | Term frequency | Term relation | Correlation |
|---|---|---|---|---|

| | | | | |
|---|---|---|---|---|
| Causal graph 1 | 'Tree density'-'Cropland' | 'tree'=3569<br>'densiti'=1171<br>'crop'=247<br>'openland'=1<br>'sylvicultur'=1<br>'longfallow'=4<br>'forestdu'=4 | Direct<br>'tree'&'crop'<br>'densiti'&'crop'<br><br>Indirect<br>'tree'&'openland'<br>'tree'&'sylvicultur'<br>'densiti'&'longfallow'<br>'crop'&'forestdu' | <br><0.1<br><0.1<br><br><br>0.11<br>0.11<br>0.11<br>0.23 |
| | 'Cropland' -'Human developm ent' | 'crop'=247<br>'human'=863<br>'develop'=1701<br>'netpresentvalu'=1<br>'deregul'=1<br>'arabl'=6<br>'cash'=27<br>'yield'=396<br>'exportori'=2<br>'agroexport'=1<br>'nationallevel'=1 | Direct<br>'crop'&'human'<br>'crop'&'develop'<br><br>Indirect<br>'crop'&'netpresentvalu'<br>'crop'&'deregul'<br>'arabl'&'deregul'<br>'crop'&'cash'<br>'crop'&'yield'<br>'crop'&'exportori'<br>'crop'&'agroexport'<br>'crop'&'nationallevel' | <br><0.10<br><0.10<br><br><br>0.15<br>0.15<br>0.71<br>0.14<br>0.13<br>0.13<br>0.11<br>0.10 |
| | 'Human developm ent'-'Human footprint' | 'human'=863<br>'develop'=1701<br>'footprint'=24<br>'nitrit'=2<br>'nitrat'=22<br>'ammonia'=9<br>'stock'=307<br>'econom'=581<br>'industri'=361<br>'impact'=1048<br>'biodiv'=984 | Direct<br>'human'&'footprint'<br>'develop'&'footprint'<br><br>Indirect<br>'economicallyvalu'&'nitrit'<br>'economicallyvalu'&'nitrat'<br>'economicallyvalu'&'ammon ia'<br>'economicallyvalu'&'stock'<br>'develop'&'econom'<br>'develop'&'industr'<br>'develop'&'impact'<br>'develop'&'biodiv' | <br><0.10<br><0.10<br><br><br>0.71<br>0.17<br>0.15<br>0.14<br>0.23<br>0.16<br>0.11<br>0.10 |
| | 'Human footprint'-'Tree density' | 'footprint'=24<br>'tree'=3569<br>'densiti'=1171<br>'ecosystem'=853<br>'plantat'=1465<br>'deforest'=539 | Direct<br>'footprint'&'tree'<br>'footprint'&'densiti'<br><br>Indirect<br>'impact'&'forest'<br>'impact'&'biodiv'<br>'impact'&'ecosystem'<br>'impact'&'plantat'<br>'impact'&'deforest' | <br><0.10<br><0.10<br><0.10<br><0.10<br><br>0.20<br>0.17<br>0.13<br>0.12<br>0.11 |
| | 'Tree density'-'Human developm ent' | 'human'=863<br>'develop'=1701<br>'tree'=3569<br>'densiti'=1171<br>'harvest'=273<br>'disturbanceknowledg'=1 | Direct<br>'human'&'tree'<br>'human'&'densiti'<br>'develop'&'tree'<br>'develop'&'densiti'<br><br>Indirect | <br><0.10<br><0.10<br><0.10<br><0.10 |

| | | | | |
|---|---|---|---|---|
| | | | 'tree'&'harvest'<br>'densiti'&'<br>disturbanceknowledg'<br>'densiti'&' economicallyvalu' | 0.15<br>0.13<br>0.11 |
| | 'Cropland'<br>-'Human<br>footprint' | 'footprint'=24<br>'crop'=247<br>'weedi'=6<br>'extensif'=2 | Direct<br>'footprint'&'crop'<br><br>Indirect<br>'crop'&'weedi'<br>'crop'&'extensif' | <br><0.10<br><br><br>0.26<br>0.26 |
| Causal<br>graph 2 | 'Species<br>richness'-<br>'Food<br>area' | 'speci'=11860<br>'food'=682<br>'rich'=918 | Direct<br>'speci'&'food'<br>'rich'&'food'<br><br>Indirect | <br><0.10<br><0.10 |
| | 'Food<br>area'-<br>'Investme<br>nt benefit<br>factor' | 'benefit'=342<br>'invest'=113<br>'food'=682<br>'forestagricultur'=2<br>'soilconserv'=1 | Direct<br>'food'&'benefit'<br>'food'&'invest'<br><br>Indirect<br>'benefit'&'forestagricultur'<br>'invest'&'soilconserv' | <br><0.10<br><0.10<br><br><br>0.15<br>0.32 |
| | 'Investme<br>nt benefit<br>factor'-<br>'phosphor<br>us<br>loading' | 'phosphorus'=143<br>'benefit'=342<br>'invest'=113<br>'effici'=276 | Direct<br>'phosphorus'&'benefit'<br><br>Indirect<br>'phosphorus'&'effici' | <br><0.10<br><br><br>0.22 |
| | 'Phosphor<br>us<br>loading'-<br>'Species<br>richness' | 'phosphorus'=143<br>'speci'=11860<br>'rich'=918 | Direct<br>'phosphorus'&'speci'<br>'phosphorus'&'rich'<br><br>Indirect | <br><0.10<br><0.10 |
| | 'Species<br>richness'-<br>'Investme<br>nt benefit<br>factor' | 'speci'=11860<br>'rich'=918<br>'invest'=113<br>'benefit'=342 | Direct<br>'invest'&'speci'<br>'invest'&'rich'<br>'benefit'&'speci'<br>'benefit'&'rich'<br><br>Indirect | <br><0.10<br><0.10 |
| | 'Food<br>area'-<br>'Phosphor<br>us<br>loading' | 'food'=682<br>'phosphorus'=143 | Direct<br>'food'&'phosphorus'<br><br>Indirect | <br><0.10 |
| Causal<br>graph 3 | 'Alpha<br>diversity'-<br>'Soil<br>water<br>capacity' | 'alpha'=66<br>'diver'=2162<br>'soil'=2502<br>'water'=1271<br>'moistur'=125 | Direct<br>'alpha'&'soil'<br>'alpha'&'water'<br>'alpha'&'moistur'<br>'diver'&''soil'<br>'diver'&'water'<br>'diver'&'moistur' | <br><0.10<br><0.10<br><0.10 |

| | | | | |
|---|---|---|---|---|
| | | | Indirect | |
| | 'Soil water capacity'-'Population density' | 'soil'=2502<br>'water'=1271<br>'moistur'=125<br>'populationlevel'=6<br>'densiti'=1171<br><br>'wetter'=12 | Direct<br><br>Indirect<br>'populationlevel'&'wetter' | 0.11 |
| | 'Population density'-'Forest loss' | 'populationlevel'=6<br>'densiti'=1171<br>'forest'=10030<br>'loss'=705<br>'specieslevel'=16 | Direct<br><br>Indirect<br>'populationlevel'&'specieslevel' | 0.10 |
| | 'Forest loss'-'Alpha diversity' | 'forest'=10030<br>'loss'=705<br>'alpha'=66<br>'diver'=2162<br>'log'=2208<br>'biodiv'=984 | Direct<br><br>Indirect<br>'log'&'biodiv' | 0.14 |
| | 'Alpha diversity'-'Population density' | 'alpha'=66<br>'diver'=2162<br>'populationlevel'=6<br>'densiti'=1171 | Direct<br><br>Indirect | |
| | 'Soil water capacity'-'Forest loss' | 'soil'=2502<br>'water'=1271<br>'moistur'=125<br>'forest'=10030<br>'loss'=705 | Direct<br><br>Indirect | |

**Supporting Information References**

21. B. S. Steidinger, T. W. Crowther, J. Liang, M. E. Van Nuland, G. D. A. Werner, P. B. Reich, G. J. Nabuurs, S. de-Miguel, M. Zhou, N. Picard, B. Herault, X. Zhao, C. Zhang, D. Routh, K. G. Peay, M. Abegg, C. Y. Adou Yao, G. Alberti, A. Almeyda Zambrano, E. Alvarez-Davila, P. Alvarez-Loayza, L. F. Alves, C. Ammer, C. Antón-Fernández, A. Araujo-Murakami, L. Arroyo, V. Avitabile, G. Aymard, T. Baker, R. Bałazy, O. Banki, J. Barroso, M. Bastian, J.-F. Bastin, L. Birigazzi, P. Birnbaum, R. Bitariho, P. Boeckx, F. Bongers, O. Bouriaud, P. H. S. Brancalion, S. Brandl, F. Q. Brearley, R. Brienen, E. Broadbent, H. Bruelheide, F. Bussotti, R. Cazzolla Gatti, R. Cesar, G. Cesljar, R. Chazdon, H. Y. H. Chen, C. Chisholm, E. Cienciala, C. J. Clark, D. Clark, G. Colletta, R. Condit, D. Coomes, F. Cornejo Valverde, J. J. Corral-Rivas, P. Crim, J. Cumming, S. Dayanandan, A. L. de Gasper, M. Decuyper, G. Derroire, B. DeVries, I. Djordjevic, A. Iêda, A. Dourdain, N. L. E. Obiang, B. Enquist, T. Eyre, A. B. Fandohan, T. M. Fayle, T. R. Feldpausch, L. Finér, M. Fischer, C. Fletcher, J. Fridman, L. Frizzera, J. G. P. Gamarra, D. Gianelle, H. B. Glick, D. Harris, A. Hector, A. Hemp, G. Hengeveld, J. Herbohn, M. Herold, A. Hillers, E. N. Honorio Coronado, M. Huber, C. Hui, H. Cho, T. Ibanez, I. Jung, N. Imai, A. M. Jagodzinski, B. Jaroszewicz, V. Johannsen, C. A. Joly, T. Jucker, V. Karminov, K. Kartawinata, E. Kearsley, D. Kenfack, D. Kennard, S. Kepfer-Rojas, G. Keppel, M. L. Khan, T. Killeen, H. S. Kim, K. Kitayama, M. Köhl, H. Korjus, F. Kraxner, D. Laarmann, M. Lang, S. Lewis, H. Lu, N. Lukina, B. Maitner, Y. Malhi, E. Marcon, B. S. Marimon, B. H. Marimon-Junior, A. R. Marshall, E. Martin, O. Martynenko, J. A. Meave, O. Melo-Cruz, C. Mendoza, C. Merow, A. Monteagudo Mendoza, V. Moreno, S. A. Mukul, P. Mundhenk, M. G. Nava-Miranda, D. Neill, V. Neldner, R. Nevenic, M. Ngugi, P. Niklaus, J. Oleksyn, P. Ontikov, E. Ortiz-Malavasi, Y. Pan, A. Paquette, A. Parada-Gutierrez, E. Parfenova, M. Park, M. Parren, N. Parthasarathy, P. L. Peri, S. Pfautsch, O. Phillips, M. T. Piedade, D. Piotto, N. C. A. Pitman, I. Polo, L. Poorter, A. D. Poulsen, J. R. Poulsen, H. Pretzsch, F. Ramirez Arevalo, Z. Restrepo-Correa, M. Rodeghiero, S. Rolim, A. Roopsind, F. Rovero, E. Rutishauser, P. Saikia, P. Saner, P. Schall, M.-J. Schelhaas, D.

Schepaschenko, M. Scherer-Lorenzen, B. Schmid, J. Schöngart, E. Searle, V. Seben, J. M. Serra-Diaz, C. Salas-Eljatib, D. Sheil, A. Shvidenko, J. Silva-Espejo, M. Silveira, J. Singh, P. Sist, F. Slik, B. Sonké, A. F. Souza, K. Stereńczak, J.-C. Svenning, M. Svoboda, N. Targhetta, N. Tchebakova, H. ter Steege, R. Thomas, E. Tikhonova, P. Umunay, V. Usoltsev, F. Valladares, F. van der Plas, T. Van Do, R. Vasquez Martinez, H. Verbeeck, H. Viana, S. Vieira, K. von Gadow, H.-F. Wang, J. Watson, B. Westerlund, S. Wiser, F. Wittmann, V. Wortel, R. Zagt, T. Zawila-Niedzwiecki, Z.-X. Zhu, I. C. Zo-Bi, G. consortium, Climatic controls of decomposition drive the global biogeography of forest-tree symbioses. *Nature*. **569**, 404–408 (2019).

22. J. van den Hoogen, S. Geisen, D. Routh, H. Ferris, W. Traunspurger, D. A. Wardle, R. G. M. de Goede, B. J. Adams, W. Ahmad, W. S. Andriuzzi, R. D. Bardgett, M. Bonkowski, R. Campos-Herrera, J. E. Cares, T. Caruso, L. de Brito Caixeta, X. Chen, S. R. Costa, R. Creamer, J. Mauro da Cunha Castro, M. Dam, D. Djigal, M. Escuer, B. S. Griffiths, C. Gutiérrez, K. Hohberg, D. Kalinkina, P. Kardol, A. Kergunteuil, G. Korthals, V. Krashevska, A. A. Kudrin, Q. Li, W. Liang, M. Magilton, M. Marais, J. A. R. Martín, E. Matveeva, E. H. Mayad, C. Mulder, P. Mullin, R. Neilson, T. A. D. Nguyen, U. N. Nielsen, H. Okada, J. E. P. Rius, K. Pan, V. Peneva, L. Pellissier, J. Carlos Pereira da Silva, C. Pitteloud, T. O. Powers, K. Powers, C. W. Quist, S. Rasmann, S. S. Moreno, S. Scheu, H. Setälä, A. Sushchuk, A. V Tiunov, J. Trap, W. van der Putten, M. Vestergård, C. Villenave, L. Waeyenberge, D. H. Wall, R. Wilschut, D. G. Wright, J. Yang, T. W. Crowther, Soil nematode abundance and functional group composition at a global scale. *Nature*. **572**, 194–198 (2019).

23. T. Hengl, J. Mendes de Jesus, G. B. M. Heuvelink, M. Ruiperez Gonzalez, M. Kilibarda, A. Blagotić, W. Shangguan, M. N. Wright, X. Geng, B. Bauer-Marschallinger, M. A. Guevara, R. Vargas, R. A. MacMillan, N. H. Batjes, J. G. B. Leenaars, E. Ribeiro, I. Wheeler, S. Mantel, B. Kempen, SoilGrids250m: Global gridded soil information based on machine learning. *PLoS One*. **12**, e0169748 (2017).

24. J. J. Danielson, D. Gesch, "Global multi-resolution terrain elevation data 2010 (GMTED2010)." (2011).

30. T. W. Crowther, H. B. Glick, K. R. Covey, C. Bettigole, D. S. Maynard, S. M. Thomas, J. R. Smith, G. Hintler, M. C. Duguid, G. Amatulli, M.-N. Tuanmu, W. Jetz, C. Salas, C. Stam,

D. Piotto, R. Tavani, S. Green, G. Bruce, S. J. Williams, S. K. Wiser, M. O. Huber, G. M. Hengeveld, G.-J. Nabuurs, E. Tikhonova, P. Borchardt, C.-F. Li, L. W. Powrie, M. Fischer, A. Hemp, J. Homeier, P. Cho, A. C. Vibrans, P. M. Umunay, S. L. Piao, C. W. Rowe, M. S. Ashton, P. R. Crane, M. A. Bradford, Mapping tree density at a global scale. *Nature*. **525**, 201–205 (2015).

31. A. M. Wilson, W. Jetz, Remotely Sensed High-Resolution Global Cloud Dynamics for Predicting Ecosystem and Biodiversity Distributions. *PLOS Biol.* **14**, e1002415 (2016).

32. B. Poulter, L. Aragão, N. Andela, V. Bellassen, P. Ciais, T. Kato, X. Lin, B. Nachin, S. Luyssaert, N. Pederson, P. Peylin, S. Piao, T. Pugh, S. Saatchi, D. Schepaschenko, M. Schelhaas, A. Shivdenko, "The global forest age dataset and its uncertainties (GFADv1.1)." (2019).

33. M. Santoro, GlobBiomass - global datasets of forest biomass (2018), , doi:10.1594/PANGAEA.894711.

34. M. C. Hansen, P. V. Potapov, R. Moore, M. Hancher, S. A. Turubanova, A. Tyukavina, D. Thau, S. V. Stehman, S. J. Goetz, T. R. Loveland, A. Kommareddy, A. Egorov, L. Chini, C. O. Justice, J. R. G. Townshend, High-resolution global maps of 21st-century forest cover change. *Science (80-. ).* (2013), doi:10.1126/science.1244693.

35. S. Running, M. Zhao, "MOD17A3HGF MODIS/Terra Net Primary Production Gap-Filled Yearly L4 Global 500 m SIN Grid V006." (2019).

48. L. N. Hudson, T. Newbold, S. Contu, S. L. L. Hill, I. Lysenko, A. De Palma, H. R. P. Phillips, R. A. Senior, D. J. Bennett, H. Booth, A. Choimes, D. L. P. Correia, J. Day, S. Echeverría-Londoño, M. Garon, M. L. K. Harrison, D. J. Ingram, M. Jung, V. Kemp, L. Kirkpatrick, C. D. Martin, Y. Pan, H. J. White, J. Aben, S. Abrahamczyk, G. B. Adum, V. Aguilar-Barquero, M. A. Aizen, M. Ancrenaz, E. Arbeláez-Cortés, I. Armbrecht, B. Azhar, A. B. Azpiroz, L. Baeten, A. Báldi, J. E. Banks, J. Barlow, P. Batáry, A. J. Bates, E. M. Bayne, P. Beja, Å. Berg, N. J. Berry, J. E. Bicknell, J. H. Bihn, K. Böhning-Gaese, T. Boekhout, C. Boutin, J. Bouyer, F. Q. Brearley, I. Brito, J. Brunet, G. Buczkowski, E. Buscardo, J. Cabra-García, M. Calviño-Cancela, S. A. Cameron, E. M. Cancello, T. F. Carrijo, A. L. Carvalho, H. Castro, A. A. Castro-Luna, R. Cerda, A. Cerezo, M. Chauvat, F. M. Clarke, D. F. R. Cleary, S. P. Connop, B. D'Aniello, P. G. da Silva, B. Darvill, J. Dauber, A. Dejean, T. Diekötter, Y. Dominguez-Haydar, C. F. Dormann, B. Dumont, S. G. Dures, M. Dynesius, L. Edenius, Z. Elek, M. H. Entling, N. Farwig, T. M. Fayle, A. Felicioli, A. M. Felton, G. F. Ficetola, B. K. C. Filgueiras, S. J. Fonte, L. H. Fraser, D. Fukuda, D. Furlani, J. U. Ganzhorn, J. G. Garden, C. Gheler-Costa, P. Giordani, S. Giordano, M. S. Gottschalk, D. Goulson, A. D. Gove, J. Grogan, M. E. Hanley, T. Hanson, N. R. Hashim, J. E. Hawes, C. Hébert, A. J. Helden, J. A. Henden, L. Hernández, F. Herzog, D. Higuera-Diaz, B. Hilje, F. G. Horgan, R. Horváth, K. Hylander, P. Isaacs-Cubides, M. Ishitani, C. T. Jacobs, V. J. Jaramillo, B. Jauker, M. Jonsell, T. S. Jung, V. Kapoor, V. Kati, E. Katovai, M. Kessler, E. Knop, A. Kolb, Á. Korösi, T. Lachat, V. Lantschner, V. Le Féon, G. Lebuhn, J. P. Légaré, S. G. Letcher, N. A. Littlewood, C. A. López-Quintero, M. Louhaichi, G. L. Lövei, M. E. Lucas-Borja, V. H. Luja, K. Maeto, T. Magura, N. A. Mallari, E. Marin-Spiotta, E. J. P. Marshall, E. Martínez, M. M. Mayfield, G. Mikusinski, J. C. Milder, J. R. Miller, C. L. Morales, M. N. Muchane, M. Muchane, R. Naidoo, A. Nakamura, S. Naoe, G. Nates-Parra, D. A. Navarrete Gutierrez, E. L. Neuschulz, N. Noreika, O. Norfolk, J. A. Noriega, N. M. Nöske, N. O'Dea, W. Oduro, C. Ofori-Boateng, C. O. Oke, L. M. Osgathorpe, J. Paritsis, A. Parra-H, N. Pelegrin, C. A. Peres, A. S. Persson, T. Petanidou, B. Phalan, T. K. Philips, K. Poveda, E. F. Power, S. J. Presley, V. Proença, M. Quaranta, C. Quintero, N. A. Redpath-Downing, J. L. Reid, Y. T. Reis, D. B. Ribeiro, B. A. Richardson, M. J. Richardson, C. A. Robles, J. Römbke, L. P. Romero-Duque, L. Rosselli, S. J. Rossiter, T. H. Roulston, L. Rousseau, J. P. Sadler, S. Sáfián, R. A. Saldaña-Vázquez, U. Samnegård, C. Schüepp, O. Schweiger, J. L. Sedlock, G. Shahabuddin, D. Sheil, F. A. B. Silva, E. M. Slade, A. H. Smith-Pardo, N. S. Sodhi, E. J. Somarriba, R. A. Sosa, J. C. Stout, M. J. Struebig, Y. H. Sung, C. G. Threlfall, R. Tonietto, B. Tóthmérész, T. Tscharntke, E. C.

Turner, J. M. Tylianakis, A. J. Vanbergen, K. Vassilev, H. A. F. Verboven, C. H. Vergara, P. M. Vergara, J. Verhulst, T. R. Walker, Y. Wang, J. I. Watling, K. Wells, C. D. Williams, M. R. Willig, J. C. Z. Woinarski, J. H. D. Wolf, B. A. Woodcock, D. W. Yu, A. S. Zaitsev, B. Collen, R. M. Ewers, G. M. Mace, D. W. Purves, J. P. W. Scharlemann, A. Purvis, The PREDICTS database: A global database of how local terrestrial biodiversity responds to human impacts. *Ecol. Evol.* (2014), doi:10.1002/ece3.1303.

49. A. De Palma, K. Sanchez-Ortiz, P. A. Martin, A. Chadwick, G. Gilbert, A. E. Bates, L. Börger, S. Contu, S. L. L. Hill, A. Purvis, in *Next Generation Biomonitoring: Part 1*, D. A. Bohan, A. J. Dumbrell, G. Woodward, M. B. T.-A. in E. R. Jackson, Eds. (Academic Press, 2018; https://www.sciencedirect.com/science/article/pii/S0065250417300296), vol. 58, pp. 163–199.

50. A. J. Hoskins, A. Bush, J. Gilmore, T. Harwood, L. N. Hudson, C. Ware, K. J. Williams, S. Ferrier, Downscaling land-use data to provide global 30″ estimates of five land-use classes. *Ecol. Evol.* **6**, 3040–3055 (2016).

51. S. van Asselen, P. H. Verburg, Land cover change or land-use intensification: simulating land system change with a global-scale land change model. *Glob. Chang. Biol.* **19**, 3648–3667 (2013).

52. T. Newbold, L. N. Hudson, A. P. Arnell, S. Contu, A. De Palma, S. Ferrier, S. L. L. Hill, A. J. Hoskins, I. Lysenko, H. R. P. Phillips, V. J. Burton, C. W. T. Chng, S. Emerson, D. Gao, G. P. Hale, J. Hutton, M. Jung, K. Sanchez-Ortiz, B. I. Simmons, S. Whitmee, H. Zhang, J. P. W. Scharlemann, A. Purvis, Has land use pushed terrestrial biodiversity beyond the planetary boundary? A global assessment. *Science (80-. ).* (2016), doi:10.1126/science.aaf2201.

53. T. Newbold, L. N. Hudson, S. L. L. Hill, S. Contu, I. Lysenko, R. A. Senior, L. Börger, D. J. Bennett, A. Choimes, B. Collen, J. Day, A. De Palma, S. Díaz, S. Echeverria-Londoño, M. J. Edgar, A. Feldman, M. Garon, M. L. K. Harrison, T. Alhusseini, D. J. Ingram, Y. Itescu, J. Kattge, V. Kemp, L. Kirkpatrick, M. Kleyer, D. L. P. Correia, C. D. Martin, S. Meiri, M. Novosolov, Y. Pan, H. R. P. Phillips, D. W. Purves, A. Robinson, J. Simpson, S. L. Tuck, E. Weiher, H. J. White, R. M. Ewers, G. M. MacE, J. P. W. Scharlemann, A. Purvis, Global effects of land use on local terrestrial biodiversity. *Nature* (2015), doi:10.1038/nature14324.

54. C. Gascon, *Amphibian conservation action plan: proceedings IUCN/SSC Amphibian Conservation Summit 2005* (IUCn, 2007).

60. D. M. Danko, "The Digital Chart of the World" (1992).

61. Ucl. & ESA, "GLOBCOVER 2009 Products Description and Validation Report" (2011).

62. N. I. and M. Agency, "Department of Defense World Geodetic System 1984: Its Definition and Relationships with Local Geodetic Systems" (1997).

81. P. Bicheron, V. Amberg, L. Bourg, D. Petit, M. Huc, B. Miras, C. Brockmann, S. Delwart, F. Ranéra, O. Hagolle, in *Proceedings of MERIS/AATSR Colloque, Frascati* (2008).

82. Q. Yu, L. You, U. Wood-Sichra, Y. Ru, A. K. B. Joglekar, S. Fritz, W. Xiong, M. Lu, W. Wu, P. Yang, A cultivated planet in 2010 -- Part 2: The global gridded agricultural-production maps. *Earth Syst. Sci. Data*. **12**, 3545–3572 (2020).

83. U. Wood-Sichra, A. B. Joglekar, L. You, "Spatial Production Allocation Model (SPAM) 2005" (2016).

84. S. Cohen, A. J. Kettner, J. P. M. Syvitski, B. M. Fekete, WBMsed, a distributed global-scale riverine sediment flux model: Model description and validation. *Comput. Geosci.* **53**, 80–93 (2013).

85. J. P. M. Syvitski, J. D. Milliman, Geology, geography, and humans battle for dominance over the delivery of fluvial sediment to the coastal ocean. *J. Geol.* **115**, 1–19 (2007).

86. J. R. Meijer, M. A. J. Huijbregts, K. C. G. J. Schotten, A. M. Schipper, Global patterns of current and future road infrastructure. *Environ. Res. Lett.* **13**, 64006 (2018).

87. S. Shimizu, T. Inazumi, Y. Sogawa, A. Hyvärinen, Y. Kawahara, T. Washio, P. O. Hoyer, K. Bollen, DirectLiNGAM: A Direct Method for Learning a Linear Non-Gaussian Structural Equation Model. *J. Mach. Learn. Res.* **12**, 1225–1248 (2011).

88. S. Shimizu, P. O. Hoyer, A. Hyvärinen, A. Kerminen, A linear non-gaussian acyclic model

for causal discovery. *J. Mach. Learn. Res.* **7** (2006).
